# Distinct Tubulin C-Terminal Tails Control the Efficiency of a Microtubule Severing Machine

**DOI:** 10.64898/2026.05.28.728271

**Authors:** Madhavie Ranpati Dewage, Shehani Kahawatte, Jennifer L. Ross, Ruxandra I. Dima

## Abstract

Microtubule severing enzymes of the AAA+ family are essential regulators of cytoskeletal remodeling, extracting subunits from the microtubule lattice through ATP-driven conformational changes. Among them, katanin assembles into hexameric structures and binds the negatively charged carboxy-terminal tails (CTTs) of tubulin through its central pore. Experimental studies have shown that different tubulin CTT isotypes can act as either inhibitors or non-inhibitors of katanin-mediated severing, with increased CTT hydrophobicity associated with reduced inhibition. However, the molecular basis underlying this selective behavior remains poorly understood. Here, we employed molecular dynamics simulations, and quantitative analysis combining principal component analysis, clustering, and distance distribution analysis, to investigate how natural tubulin CTTs (beta5, beta4b, and beta3), and engineered CTTs (beta5-A+Y, beta5-cterm, beta5-midpoint, and poly-E) influence katanin structure and dynamics in spiral and ring conformations. Our results show that inhibitory CTTs form stronger interactions with the terminal protomers and increase flexibility in the inner protomers, resulting in coordinated motions associated with pore narrowing. In contrast, non-inhibitory CTTs preferentially interact with inner protomers, disrupt interprotomer coordination, weakening the collective grip of the hexamer on the substrate. Analyses of the engineered constructs further revealed that inhibitory behavior is governed primarily by the spatial distribution of acidic residues rather than the overall charge. Additionally, we identified species-specific responses of different CTT isotypes toward the katanin ring state. Together, these findings provide molecular-level insight into how tubulin CTT sequence organization regulates substrate recognition, pore dynamics, and severing efficiency, leading to predictive design of CTTs with a desired action on katanin.

## I. INTRODUCTION

Microtubules (MTs) are highly dynamic, hollow polymers found in nearly all eukaryotic cells. They are composed of *α* and *β*-tubulin dimers, which align longitudinally to form protofilaments that are arranged laterally into a cylindrical MT structure.^1–4^ As key components of the cytoskeleton, MTs are essential for maintaining cell shape, intracellular transport, and mitotic spindle assembly during cell division.^1,5,6^ Their functions are regulated by microtubule-associated proteins (MAPs), including microtubule-severing enzymes. These severing enzymes belong to the AAA+ (ATPases Associated with diverse cellular Activities) superfamily and use ATP hydrolysis to exert mechanical force and create internal breaks in the MT lattice^3–5,7,8^. The severing proteins play critical roles in cellular processes such as spindle formation, cytoskeletal remodeling, ciliogenesis, and axon branching during neurodevelopment.^3,9–11^

Katanin was the first MT-severing enzyme to be identified and plays a key role in neurodevelopment by regulating axonal growth, dendritic branching, and synaptic maintenance.^12,13^ Mutations in katanin have been linked to severe neurodevelopmental disorders, including microcephaly and lissencephaly, highlighting its importance in maintaining proper brain structure and function.^14–16^ The severing activity of katanin is mediated by its AAA+ ATPase motor domain, which is composed of two subdomains: the nucleotide-binding domain (NBD) and the helical bundle domain (HBD). The NBD contains several functionally conserved motifs, including Walker A, Walker B, and the Arg-finger, all of which are essential for ATP binding and hydrolysis. In addition, pore loop 1 (PL1) and pore loop 2 (PL2), located within the NBD, are critical for substrate binding and contribute directly to the severing mechanism, while pore loop 3 (PL3) acts as a bridge connecting substrate binding PLs to the ATP-binding sites.^14,17,18^.

For katanin from *Caenorhabditis elegans* two conformations were solved by cryo-EM: the spiral conformation (PDB ID: 6UGD) and the ring conformation (PDB ID: 6UGE). The spiral conformation has a right-handed spiral arrangement of subunits, with a ∼ 40 Å wide gate between the terminal protomers A and F. All the nucleotide-binding pockets in this state are occupied by ATP. Upon ATP hydrolysis, protomer A becomes more flexible and closes the gate between the terminal protomers, leading to the ring conformation. The mechanical force generated during the transition from spiral to ring, driven by ATP hydrolysis, is thought to act on the MT lattice, ultimately resulting in severing.^14,17,19^.

Studies showed that, in the presence of ATP, katanin oligomerizes into hexamers and interacts with the carboxy terminal tails (CTTs) of tubulin, which are disordered and negatively charged amino acid sequences extending from the MT surface.^3,14,17,20^ PL1 and PL2 form an electropositive double spiral, which wraps around the negatively charged CTT and grips it through the central pore leading to the extraction of tubulin dimers from the MT lattice.^14,17^

According to experimental findings, CTTs play a significant role in the severing mechanism, as severing enzymes directly bind to these tails. Due to variations in genetic coding and post-translational modifications, CTTs adopt different sequences, called isotypes, while the tubulin core remains much more conserved.^21–24^ The CTTs differ in their length, the number of acidic residues, and the distribution of these acidic residues throughout the tail. These tails are also the primary sites where most post-translational modifications, such as polyglutamylation and polyglycylation, occur. This sequence variability gives rise to multiple *α* and *β*-tubulin isotypes, including beta3 (TUBB3), beta5 (TUBB5), beta4b (TUBB4B), which display distinct tissue distributions and functional specializations.^25,26^ Previous expression analyses have revealed that beta3 is enriched in neuronal tissues, beta5 is broadly expressed and essential during brain development, and beta4b is strongly associated with motile cilia and multiciliated epithelial cells, indicating isotype-specific roles in specialized MT architectures.^27–30^ Biochemical and structural studies have demonstrated that severing efficiency is strongly influenced by interactions between enzyme pore loops and the tubulin CTTs, as well as by tail posttranslational modifications, which can either enhance or antagonize enzyme activity.^31–36^

Recent experimental evidence suggests that different *β* tubulin isotypes show varying levels of susceptibility to severing by katanin.^14,36–39^ Also, in experimental setups with free CTTs, MTs, and katanin, some CTT isotypes serve as inhibitors, while others act as non-inhibitors and allow MT severing to proceed.^36^ A proposed explanation behind the selective behavior of CTTs is that inhibitors are strongly bound to katanin, while non-inhibitors bind only weakly to katanin hexamers. Among the naturally occurring beta-CTT sequences, beta5 and beta4b were found to bind strongly to katanin and to act as inhibitors. In the presence of such CTTs, MT severing is significantly reduced. On the other hand, beta3 acts as a non-inhibitor with low affinity for katanin, allowing the severing activity to occur.

Engineered constructs, such as beta5-A+Y, also function as non-inhibitors, further highlighting the importance of the final amino acid in the sequence. Additionally, experimental results have shown that polyglutamate (poly-E) and polyaspartate (poly-D) peptides act as inhibitors likely due to their strong interaction with katanin.^36^

A characteristic of the various CTTs is their highly negative charge content. However, despite variations in the total number of charged amino acids, the experimental findings did not correlate well with the total charge of the CTTs. Instead, the average hydrophobicity of the tails was found to explain better some of the results. More importantly, the lack of understanding of the mechanisms behind the inhibitory or non-inhibitory behavior of the CTTs from various tubulin isotypes represents a major gap in knowledge of the severing action. In this work, we address this outstanding problem by probing the molecular level behavior of E14, the CTT used to solve the cryo-EM structure of hexameric katanin, along with the naturally occurring human *β* CTT sequences beta5, beta4b, and beta3, and the engineered construct beta5-A+Y used in the experimental work. In addition, to investigate how the location of negatively charged residues influences severing activity, we designed two modified beta5 constructs: beta5-cterm, with the negative charges from beta5 all shifted to the C-terminal end, and beta5-midpoint, with all the charges shifted to the central region of the tail. We used a combination of docking, molecular dynamics simulations, and quantitative analyses, including contact mapping, salt bridge persistence time tracking, principal component analysis, RMSD-based clustering, center-of-mass (COM) measurements, and divergence in distance distributions, to compare natural and artificial CTT isotypes in both the spiral and ring states of katanin.

Specifically, we aimed to determine how CTT sequence organization affects the interactions with the terminal and inner protomers, how these interactions influence pore engagement and inter-protomer coordination, and how they regulate the collective motions associated with severing. In addition, we explored how these mechanisms change between the spiral and ring conformations, by focusing on the role of pore loop interactions (PL1 and PL2) in stabilizing substrate binding and enabling docking in a species-dependent manner. We found that inhibitory and non-inhibitory CTTs influence the flexibility of terminal and inner protomers in different ways. Inhibitory CTTs form stronger contacts with the terminal protomers, and increase the flexibility in the inner regions of katanin, whereas non-inhibitory CTTs tend to interact more strongly with the inner protomers. This difference leads to clear changes in how the hexamer moves and how the central pore behaves. In particular, non-inhibitory CTTs appear to disrupt the coordinated motion of the protomers and the pore stability resulting in the widening of the pore and the likely loss of the katanin’s hexamer grip on the CTT. One important finding from the artificially constructed CTT sequences is that charge positioning, rather than the total charge, determines function: concentrating acidic residues at the C-terminal end preserves the inhibitor-like behavior seen in the wild type sequence, while moving them toward the midpoint leads to non-inhibitory dynamics. In the ring state, this mechanism becomes more interesting, as CTT sequences show species-dependent behavior, with inhibitory CTTs binding to both the human and C. elegans katanin, while non-inhibitors fail to dock to the C. elegans ring. The contacts and motions in the ring state highlight the differences between inhibitors and non-inhibitors with the ring katanin across different species. Together, these results show how the CTT sequence controls substrate binding, katanin hexamer motion and activity, providing a clear picture of how substrates are recognized.

## II. METHODS

### A. System Preparation and Simulation Details

#### 1. Starting Structures

The starting structures of the katanin spiral and ring hexamers from *C. elegans* were obtained from our previous work^7^. These structures were derived from the original cryo-EM structures^14,17^ from the Protein Data Bank^40^ (PDB ID: 6UGD and PDB ID: 6UGE), with missing residues modeled using Modeller^7^. As this study focuses on katanin’s interaction with various CTT isotypes, we first generated different CTT isotypes using the PyMOL software (version2.5)^41^. Additionally, we included the minimal substrate (E14) utilized in the original cryo-EM structure. Table I lists the CTT isotypes used in the study.

**TABLE I.**
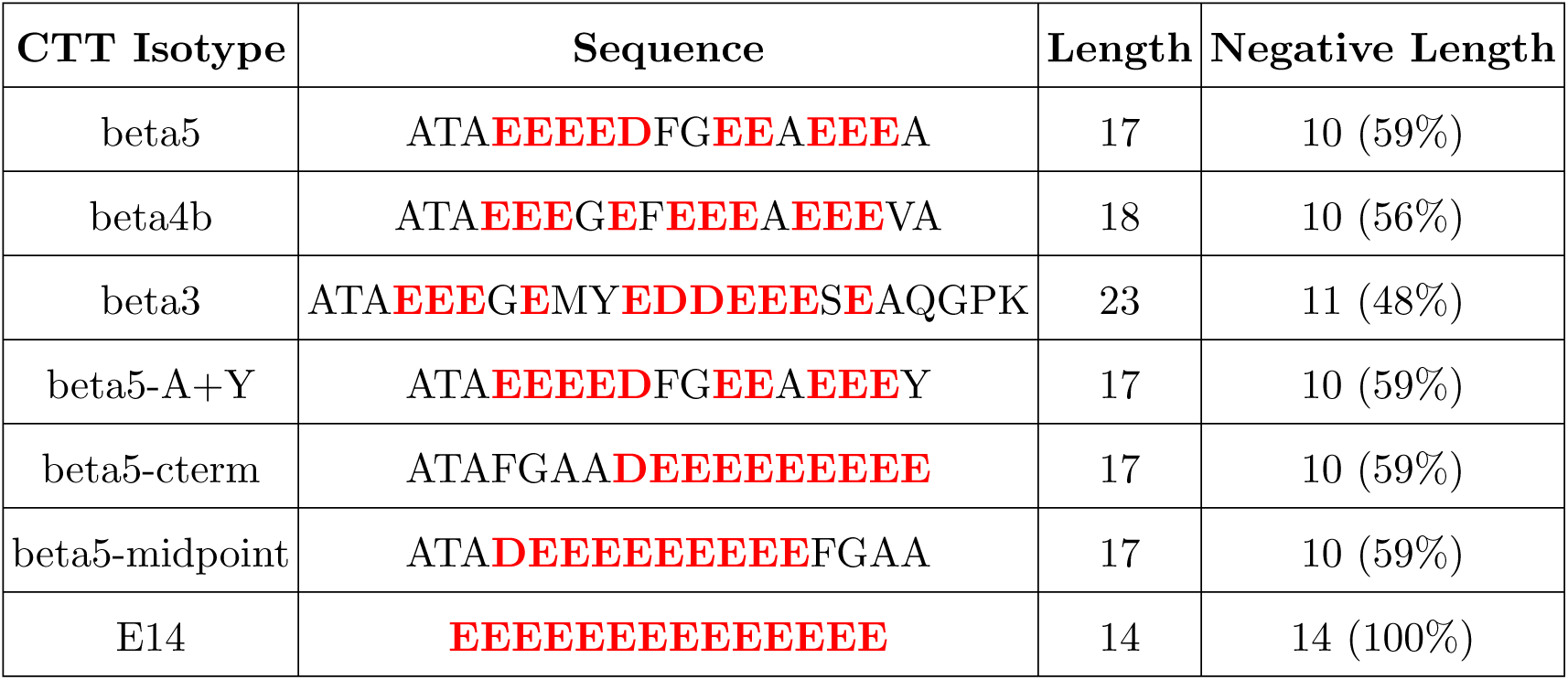
The CTT isotypes used in the study (Electro negative charged residues are in red)

The sequences of the CTT isotypes were obtained from the UniProt website^42^. After constructing each CTT isotype, we performed energy minimization using the GROMACS molecular modeling software version 2022 to ensure structural stability^43^. Each energy-minimized CTT isotype was then docked separately into the katanin structure using the GRAMM-X docking web server^44^. From the docking server, we obtained 100 docked structures. The best structure, in which the CTT was correctly oriented and positioned in the central pore of katanin, was identified by visual inspection and comparison with the orientation of the CTT present in the cryo-EM structure. A correct orientation refers to the CTT being aligned in the N- to C-terminal direction, consistent with how the peptide threads through the central pore in the cryo-EM model. Proper positioning means that the docked CTT populates states closely aligned with the experimentally resolved density, located near the center of the pore, without steric clashes. Note that all the CTT isotypes listed in Table I were successfully docked with the *C. elegans* spiral structure. In contrast, for the C. elegans ring structure, some of the CTT isotypes selected for ring-state analysis did not successfully dock, but did so with the human ring structure instead. For this analysis, we specifically focused on three CTT constructs: beta5, the natural isotype with the highest experimentally observed affinity for katanin, which docked successfully with both the *C. elegans* and human ring structures; beta3, a natural isotype that failed to dock with the *C. elegans* ring but docked successfully with the human ring structure and is known to have low binding affinity; and beta5-A+Y, an artificial isotype which also failed to dock with the *C. elegans* ring but docked with the human ring and has low binding affinity in experiments.

#### 2. Modeling the Human Katanin Ring Hexamer

Since no cryo-EM structure is available for the human katanin hexamer, we used AlphaFold3 to predict the structure of a homo-hexamer of the human KATNA1 (p60 subunit) sequence using ATP as the nucleotide to represent the ATP-bound state of the hexamer^9,45^. We note that AlphaFold3 cannot predict hetero-oligomeric structures. The predicted hexamer was composed of six identical human protomers, but the resulting conformation did not resemble either the spiral or the ring state observed in the experimental structures of *C. elegans* katanin. Therefore, this model was not considered biologically relevant and was not used for the functional analysis.

To construct the ring assembly of the human katanin, we first extracted a monomer from the AlphaFold3 hexamer, ensuring its conformation reflected the correct geometry of a monomer. This human KATNA1 monomer was then aligned to each protomer (A–F) of the *C. elegans* ring structure (PDB ID: 6UGE) using PyMOL, generating a human ring conformation with six protomers. We note that, in the cryo-EM structure of the *C. elegans* katanin ring, protomer A exhibits a distinct arrangement due to the nucleotide loss in its binding pocket, resulting in a 44 Å rotation of the NBD and an 18*°* increase in the angle between the NBD and HBD domains^17^. As a result, the human monomer structure could not be directly aligned to protomer A of the *C. elegans* ring. Instead, the sequence of the human monomer was aligned to the structure of *C. elegans* protomer A using MODELLER^46^ via ChimeraX^47^, to generate the human ring katanin protomer A suitable for inclusion in the ring model. With this modeled protomer A and the aligned protomers B to F, the human katanin ring hexamer was successfully assembled. The CTTs we selected, as mentioned in the previous section, beta5, beta5-A+Y, and beta3 were then docked into the central pore of the human ring hexamer using the GRAMM-X docking server^44^.

#### 3. Simulation Details

We performed molecular dynamics (MD) simulations using GROMACS version 2022 with the GROMOS96 54a7 force field^43,48^, employing the automated topology builder server to generate force field parameters for ATP. Each system was centered in a dodecahedral box (approximately 150 × 150 × 150 Å^3^) and solvated using the single point charge (SPC) explicit solvent model, with system neutralization achieved by adding NaCl ions and applying three-dimensional periodic boundary conditions (PBC). The simulation protocol began with energy minimization using the Verlet cutoff scheme and the steepest descent algorithm, running for 50,000 steps until reaching a convergence criterion of maximum force below 23.9 kcal/mol/Å (1000 kJ/mol/nm)^49^. Next, the system was equilibrated in two steps: first, an NVT equilibration was performed to raise the temperature to 300 K using a velocity-rescaling thermostat^50^ and the leapfrog integrator; second, an NPT equilibration was carried out to maintain the pressure at 1.0 bar, employing the Parrinello-Rahman coupling scheme with the same leapfrog integrator^51^. Both equilibration steps were run for 500 ps. For the production phase, we performed at least five independent simulations, each running for a minimum of 200 ns, resulting in a total simulation time of at least 1 microsecond per system. Throughout these simulations, we employed the LINCS algorithm^52^ to constrain hydrogen-containing bonds, allowing 2 fs integration steps, and used the Particle Mesh Ewald (PME) method to calculate electrostatic and nonbonded interactions with a 10.0 Å cutoff^7,53^. For the *C.elegans* katanin spiral structure we ran MD simulations for two setups namely the SUB state, which has just the CTT isotype bound to the katanin, and the COMPLEX state, which has both the the CTT isotype and the nucleotides (ATP) bound to katanin, while for both the ring structures (*C.elegans* and human) we only ran the COMPLEX setup. Further details about the simulations are in Tables S1 and S2.

### B. Data Analysis

#### 1. Root Mean Square Deviation (RMSD) and Root Mean Square Fluctuation (RMSF)

To assess the convergence of our simulations and the overall stability of each system, we calculated the root mean-square deviation (RMSD) of the protein backbone for each MD simulation relative to the initial structure using GROMACS. Table S3 presents the global average RMSD values for the katanin spiral conformation in both setups, comparing the different CTT isotypes, while Table S4 provides the global average RMSD values for the ring conformation of both species with various CTT isotypes. The individual RMSD plots for the katanin spiral conformation in each state are shown in Figures S1 and S2, while the RMSD plots for the ring conformation are displayed in Figure S3.

To assess how residue-level fluctuations in katanin are affected by the binding to different CTT isotypes, we calculated the root mean square fluctuation (RMSF) of the C*_α_* atoms for each MD simulation and averaged the results across all the simulations corresponding to each CTT isotype. To evaluate the changes in residue flexibility upon binding different CTT isotypes in the COMPLEX state, we computed the difference between the average RMSF of each CTT-bound system and the average RMSF of the ATP-only state, in which katanin is bound solely to ATP^7^. This comparison highlights how residue fluctuations are altered due to CTT binding in the presence of ATP. The equations used to compute RMSF are provided in Equations S1 and S2 of the Supporting Information. RMSF difference plots for the COMPLEX state are shown in Figures S4 and S5.

#### 2. RMSD based Double Clustering Analysis

To evaluate structural changes in the katanin spiral conformation induced by the binding to various CTT isotypes, we performed a two-step RMSD-based clustering analysis^10,54^. This approach allowed us to systematically compare the ATP state, where only ATP is bound, with the COMPLEX state, in which katanin is bound to both ATP and each of the CTT isotypes, revealing isotype-specific structural adaptations. For the clustering procedure, we first concatenated all the MD simulations for each setup separately (ATP state and for the COMPLEX state katanin bound to each individual CTT isotype separately). Then, the initial clustering was performed on these concatenated trajectories using an RMSD cutoff of 0.14 nm^54,55^. Clustering was performed using the ‘cluster’ tool in GROMACS, employing the GROMOS algorithm^56^.

For the second clustering step, we analyzed the central structures (those with the smallest average RMSD compared to other structures within their respective clusters) identified from the first clustering step. We determined the optimal RMSD cutoff for this second clustering by evaluating values between 0.15 nm and 0.55 nm. Our selection criterion required that at least 70% of the central structures from the first clustering step be captured within the top two clusters of the second clustering step across all setups^54^. Based on this evaluation, we selected 0.55 nm as the appropriate cutoff value for the second clustering step, with the detailed evaluation provided in Table S5. Since the top two clusters covered at least 70% of the conformations, we considered these two clusters as representative of the simulation data and the central structures of these clusters were employed for further analysis. For each CTT isotype, we separately aligned the central structure of the COMPLEX state with that of the ATP state, performing the alignment individually for each cluster. Then, RMSD values for different functional regions were calculated using the Visual Molecular Dynamics (VMD) software^57^. We performed alignments with respect to the whole protein as well as separately for the functional regions (listed in Table S6), to account for changes in overall orientation and to capture structural changes inside the functional regions. The resulting RMSD values, presented in Figures 1, 2, S6, and S7, enabled us to quantify structural changes resulting from katanin binding to different CTT isotypes. For clarity, we denote the RMSD values between the COMPLEX state with various CTT isotypes and the ATP state as RMSD_COMPLEX(CTT Isotype)→ATP._

**FIG. 1.**
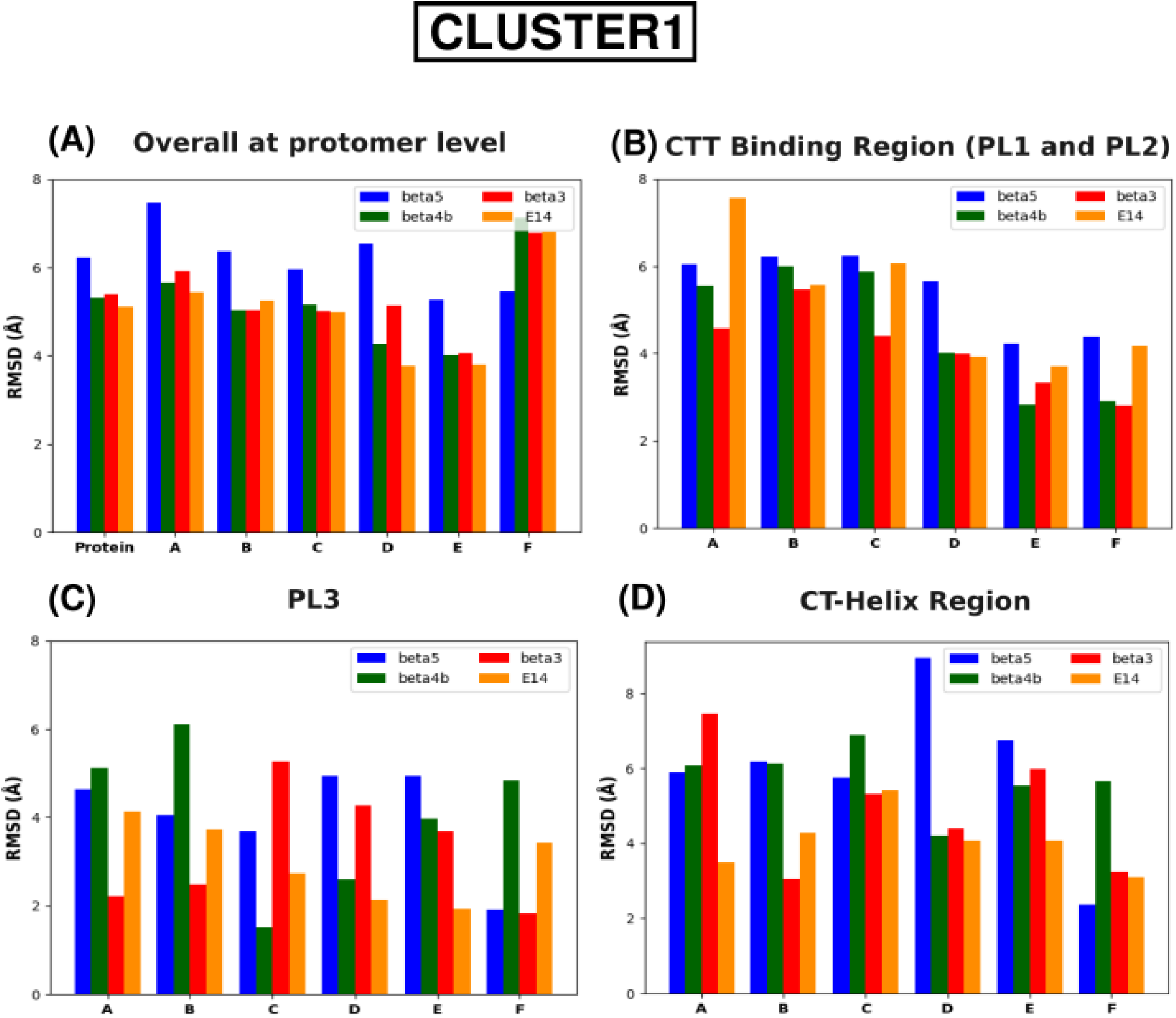
RMSD values of the central structures of the katanin spiral COMPLEX with natural CTT isotypes compared with the ATP only state for cluster1. (These values were obtained by aligning the whole protein). (A) The entire protein and each protomer (B) The CTT binding region (both PL1 and PL2) (C) The PL3 region (D) The CT-helix region. (The residue numbers of the functional regions are in Table S6).

#### 3. Essential Dynamics Analysis

To analyze the conformational dynamics of the katanin hexamer with various CTT isotypes, we conducted a detailed principal component analysis (PCA) using the MD simulations. We concatenated all the trajectories for each system separately and computed the covariance matrix using the GROMACS ‘covar’ function, considering only the C*_α_* atoms. Next, the ‘aneig’ function was used to extract the first two principal components representing the dominant motions^58–60^. This approach provided a reduced representation of the trajectory matrix, allowing us to identify the most significant motions corresponding to PC1 and PC2, which account for the largest variance observed in the MD simulations (Tables S7-S8 and S18)^60–62^. Following our previous work on MT-severing motors with minimal substrates, we analyzed the dynamics of the entire motor and the local motions of PL1 segments from each protomer, defining PL1 as the region spanning five residues before and after the positions identified experimentally (residues 258 - 277)^7,14,17,18^. To compare the essential spaces, we calculated the root-mean-square inner product (RMSIP) between the first 10 PCs, as described in Equation S3. Using RMSIP we compared the essential spaces of the natural CTT isotypes with the E14 subspace derived from the cryo-EM structure and similarly compared the beta5 modified CTT isotypes with the beta5 subspace. The results of these comparisons are provided in Tables S11–S12 and S20. Additionally, we visualized the individual PC1 and PC2 subspaces for each system, with the corresponding motions depicted using porcupine plots to highlight the observed dynamics. Global motions along PC1 are shown in Figure 3, S8 and S14, while the PL1 segment motions are illustrated in Figures 4 , S9 and S15. We also calculated the center-of-mass (COM) distances between the NBDs as well as the NBD and HBD of the terminal protomers, to characterize the distinct motion patterns of inhibitors and non-inhibitors (Table S13). The distributions of these COM distances were plotted, and Kullback–Leibler (KL) divergence values were computed to quantify similarities between the distributions, enabling comparison across CTT isotypes. The probability distribution plots of the COM distances are in Figures S10 and S16.

#### 4. Salt Bridge Analysis

Salt bridges are non-covalent electrostatic interactions between positively charged amino acids, such as lysine (Lys) or arginine (Arg), and negatively charged amino acids, such as aspartic acid (Asp) or glutamic acid (Glu). The stability and interactions between different regions of a protein can be assessed by calculating the lifetimes of salt bridges. To examine how the lifetimes of various salt bridges change due to binding of different CTT isotypes, inter- and intra-protomer salt bridge lifetimes were calculated using a maximum distance cutoff of 4 Å between N-O atoms, following our previous work^7^. This analysis was conducted using the VMD program^57^ and for comparison, 200 ns simulations were used for all systems. Specific regions, including PL1 (residues 258–278) and PL2 (residues 296–318), along with previously reported salt bridges between E14 and katanin, were only selected^7^. In this analysis, for each of the PLs, we included five residues before and after the positions reported in the literature, ensuring complete coverage of the loop region observed in the experimental structure. The lifetimes were calculated by multiplying the number of frames where the distance was less than or equal 4 Å by the time step of the simulations. Significant salt bridges were defined as those with lifetimes equal to or exceeding 1 ns and present in at least three protomers during the simulation. The significant salt bridges were then averaged over the trajectories and over the protomers. Tables S14-S17 and S21-S22 provide information about the inter and intra salt bridges with lifetimes greater than 10 ns in the katanin spiral and ring conformations when bound to different CTT isotypes. This approach provided insights into the structural stability and dynamic interactions of the protein through the identification and evaluation of significant salt bridges.

#### 5. Contact Map Analysis

We performed contact map analysis to investigate residue level interactions between the CTTs and the PL1 and PL2 regions of each protomer. We only calculated the contacts for the COMPLEX state to compare how the contacts change due to the binding of various CTT isotypes. Contact maps were generated using the ‘MDAnalysis’ python package^63^ by calculating inter-residue distances between C*α* atoms. Here, a contact was defined if the C*α* distances were within a 10Å cutoff^64^. Upon loading the trajectories and the corresponding topology file, contact frequencies were calculated and normalized across all trajectories. The resulting contact frequencies were visualized as a 2D matrix, where each element represents the fraction of frames of the simulations in which the corresponding residue pair was within 10 Å where 1 indicating that all the frames were in contact and 0 indicating that none of the frames were in contact. Figures 7, S13, and S17 show the contact maps between the various CTT isotypes with katanin’s spiral and ring conformations.

## III. RESULTS AND DISCUSSION

### A. Global and Local Structural Changes of the Katanin Spiral Conformation Upon the Binding of Different CTT Isotypes

To understand how katanin’s structure responds to the binding of CTTs from different tubulin isotypes, we examined both global and local structural changes using two quantitative approaches described in the Methods section. The RMSD-based clustering analysis identifies the dominant conformations adopted by katanin upon CTT binding, capturing both global and local structural deviations and highlighting isotype-specific effects. Complementing this, the RMSF analysis provides residue-level insights into local flexibility, specifically revealing regions sensitive to isotype-specific variations. Together, these analyses offer a multi-scale perspective on how different CTT sequences modulate katanin’s structural behavior, shedding light on the molecular basis of its substrate-specific severing activity.

When comparing the structural deviations of the entire protein between the COMPLEX state and the ATP state (i.e., due to binding different CTT isotypes) of the spiral configuration, we found an RMSD value of 6.22 Å for beta5 RMSD_COMPLEX(beta5)*→*ATP_ in cluster 1 (See Figure 1). This was the highest RMSD among the natural CTT isotypes, indicating that beta5, an experimentally validated strong katanin target, induces the most pronounced global structural change in the katanin spiral hexamer. In contrast, the RMSD values for the other natural CTT isotypes (beta4b, beta3 and the E14 from the cryo-EM structure) were similar, suggesting that their overall structural impact on katanin is comparable. At the individual protomer level, the RMSD-based clustering analysis using the whole-protein alignment, which accounts for changes in the orientation of a subunit with respect to the rest of the hexamer, shows that katanin bound to beta5 triggers substantial structural changes across all protomers except protomer F in cluster 1.

The largest structural deviation is observed in protomer A. For the other two natural CTT isotypes and for the E14 substrate, the overall structural impact at the protomer level appears to be similar for all the protomers except for protomer D where beta3 binding results in a higher RMSD value than beta4b and E14. Also, in both clusters, beta3 shows a larger structural deviation for protomer F indicating that the presence of beta3 induces significant changes in the orientation of protomer F with respect to the rest of the hexamer (See Figure 1A). When each protomer is aligned individually, which accounts for structural changes inside a subunit, we found that the binding of beta5 to the katanin spiral hexamer causes significant changes in the inner protomers C and D, which function as structural scaffolds^8^. These changes are considerably greater than those seen in the presence of the other natural CTT isotypes, indicating that the binding of beta5 induces major rearrangements in the core protomers of the complex compared to the ATP-bound state, which lacks CTTs. In contrast, when analyzing the terminal protomers, we found that beta3, an experimentally identified poor target for katanin, induces the most pronounced structural changes. Specifically, protomer F exhibits an RMSD value of 5.39 Å RMSD_COMPLEX(beta3)*→*ATP_, which is significantly higher than the corresponding RMSD value of 3.79 Å RMSD_COMPLEX(beta5)*→*ATP_ observed when katanin is bound to the strongly binding beta5. These results suggest that, while beta5 induces significant structural changes in the central protomers C and D, beta3 leads to pronounced structural deviations in the terminal protomer F (See Figure S6A). These findings apply to both changes in the orientation of the protomers with respect to the rest of the hexamer as well as to internal changes in the protomers.

Next, we aimed to identify specific functional regions within the individual protomers that undergo the most significant structural changes upon binding to the different CTT isotypes. We focused initially on PL1 and PL2, as these regions interact directly with the CTT and are likely to exhibit substrate-dependent conformational changes. In cluster 1, after aligning the entire protein, comparing the CTT-binding regions across natural CTT isotypes, we observed that all the protomers of the katanin bound to beta5 exhibit greater structural changes than in the presence of beta3. The largest differences in RMSD values between beta5 and beta3 were in protomers C and D, mirroring our previous findings for the entire protomers. Another interesting observation is that, when bound to beta4b (a moderate target for katanin), the RMSD values are similar to beta5 in protomers A, B, and C, while being similar to beta3 in protomers D, E, and F (Figure 1B). This suggests that, when katanin is bound to beta4b, the CTT-binding regions display an intermediate behavior between those seen with beta5 and beta3, respectively. When the CTT-binding regions were aligned separately, we observed a similar pattern as before. The largest RMSD differences between beta5 and beta3 were again found in protomers C, D, and E, which are all inner protomers (Figure S6B). This finding is further supported by the RMSF analysis where, in protomers D and E, both PL1 and PL2 show higher fluctuations when bound to beta5 compared to the ATP-bound state, while undergoing only negligible fluctuations when bound to beta3 (Figure S4). These results indicate that the CTT-binding regions of the inner protomers respond distinctly to the type of the bound CTT isotype, with beta5 inducing greater structural rearrangements and flexibility than beta3.

The other two functional regions of interest were PL3 and the CT-helix, which play important roles in the oligomerization of katanin. When comparing the RMSD values of PL3, the most pronounced differences between beta5 and beta3 were observed in protomers A and C. In protomer A, beta5 showed a higher RMSD than beta3, while in protomer C, beta3 exhibited a higher RMSD (Figure 1C). These findings are supported by the RMSF, where PL3 in protomer C had increased fluctuations when bound to beta3, indicating greater flexibility (Figure S4C). For the CT-helix, the largest differences in RMSD between beta5 and beta3 were in protomers A, B, and D. Specifically, beta3 had a higher RMSD than beta5 in protomer A, while beta5 led to higher RMSD values in protomers B and D (Figure 1D). Interestingly, protomer F showed minimal changes in the RMSD values of PL3 and the CT-helix for both beta5 and beta3. In terms of flexibility, RMSF analysis revealed that, in protomer A, both beta5 and beta3 reduced the flexibility of the PL3 and the CT-helix (Figure S4A). In protomer B, binding to beta5 made these regions more rigid, while in protomer D they became more flexible (Figures S4B and S4D). Together, these results suggest that, when katanin is bound to beta5, a significant structural change occurs in the PL3 of protomer A, with minimal impact on the PL3 of the other protomers. In contrast, when bound to beta3, the inner protomer C shows the greatest structural change in the PL3. For the CT-helix, the trend is reversed where binding to beta5 results in substantial structural changes in the inner protomers, while beta3 induces significant changes in protomer A.

Next, we performed a comparison between the artificially constructed beta5 variants and the original beta5 CTT. Note that the beta5-Cterm construct was not simulated in the COMPLEX state. This decision was based on its behavior in the SUB state, where essential dynamics analysis revealed strong similarities to the E14 substrate. Since E14 represents a minimal severing substrate, and beta5-Cterm exhibited similar global motions in the SUB state (as discussed in the next section), we concluded that its COMPLEX-state dynamics would also closely resemble that of E14. The RMSD values obtained when katanin was bound to either beta5-A+Y or beta5-midpoint were comparable indicating similar global structural changes in katanin due to the binding of either of these two modified CTT isotypes.

However, when compared with the RMSD changes due to beta5, the artificially constructed CTT isotypes induced less global change in katanin. At the protomer level, when aligning either the whole protein or just the individual protomers separately, protomer F showed significantly higher RMSD when bound to beta5-A+Y or beta5-midpoint compared to beta5. Specifically, when bound to beta5-A+Y, which was experimentally identified as a poor target for katanin, protomer F had a higher RMSD of 7.1 Å RMSD_COMPLEX(beta5-A+Y)*→*ATP_, compared to 5.4 Å RMSD_COMPLEX(beta5)*→*ATP_ when bound to beta5, with a similar difference in the presence of beta5-midpoint. In contrast, protomer A in katanin bound to beta5 shows considerably higher changes in the orientation with respect to the rest of the hexamer than when katanin is bound to either beta5-A+Y or beta5-midpoint (Figure 2A). However, when aligning each protomer separately, protomer A in katanin bound to beta5-A+Y behaves similarly to when katanin is bound to beta5 (Figure S7A). These results indicate that the beta5-A+Y variant induces a large change in the orientation of F with respect to the rest of the hexamer not seen in the presence of beta5, while it arrests most of the orientational motion of protomer A seen in the presence of beta5, without altering the internal structure of the protomer. Furthermore, beta5 leads to considerably higher RMSD values in protomers C and D, while the constructed CTT isotypes result in lower RMSD values, indicating that these inner protomers have less internal structural changes when bound to the artificial CTT isotypes than when bound to beta5. We note that, based on this effect on the central protomers and on the terminal protomer F, the behavior of the 2 beta constructs resembles the behavior induced by the binding of beta3 (natural non-inhibitor) discussed above.

**FIG. 2.**
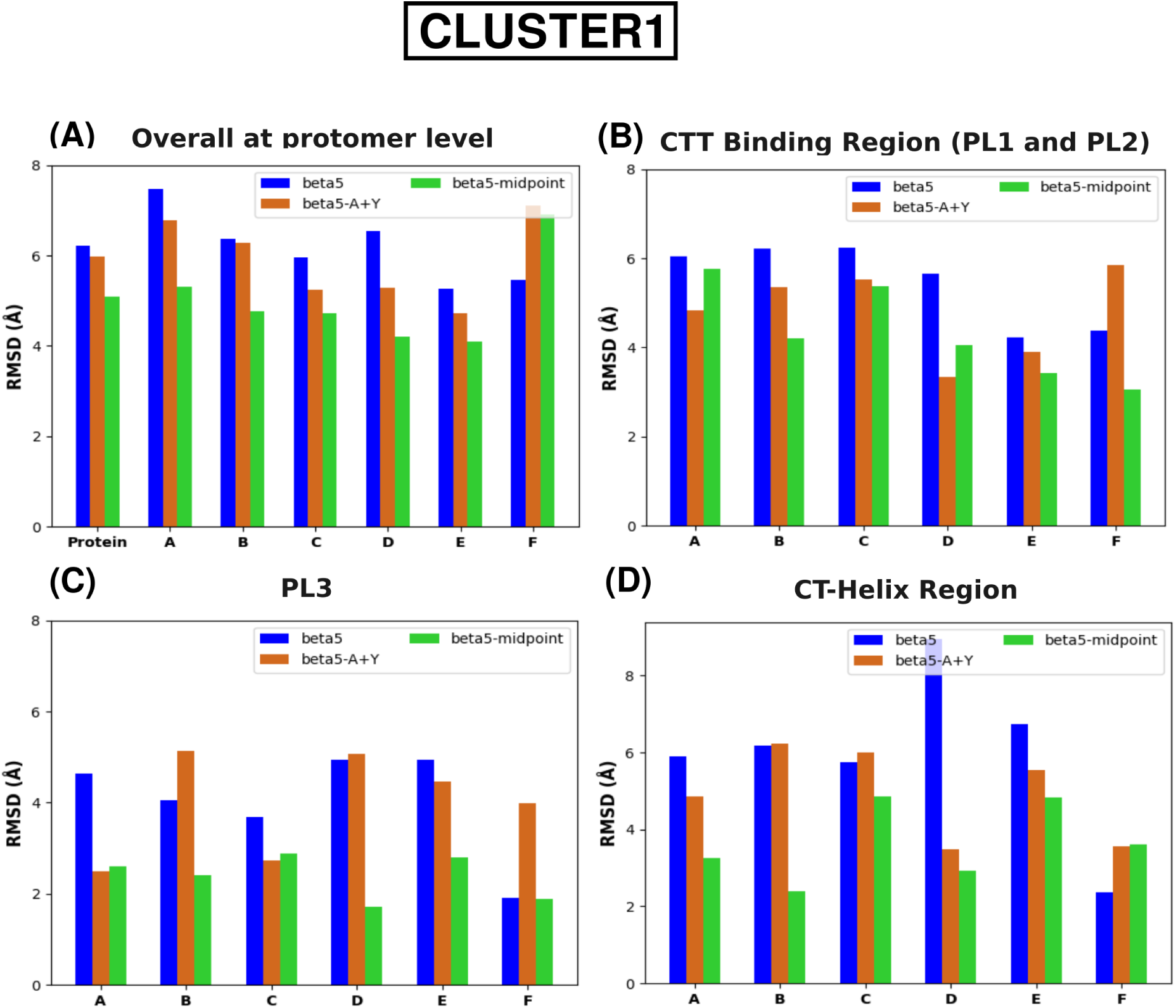
RMSD values of the central structures of the katanin spiral COMPLEX with artificial CTT isotypes compared with the ATP only state for cluster1. (These values were obtained by aligning the whole protein). (A) The entire protein and each protomer (B) The CTT binding region (both PL1 and PL2) (C) The PL3 region (D) The CT-helix region. (The residue numbers of the functional regions are in Table S6).

When analyzing PL1 and PL2, we found that, in cluster 1, beta5-A+Y results in lower RMSD values for all the protomers, except for protomer F, compared to beta5 (Figure 2B). This suggests that the CTT-binding regions from protomers A to E undergo fewer structural changes, while the CTT binding regions of protomer F show pronounced structural changes when bound to beta5-A+Y compared to beta5. RMSF analysis of protomer F further supports this finding, revealing that the PL1 and PL2 regions, when bound to beta5-A+Y, exhibit greater flexibility compared to the ATP-bound state, whereas beta5 leads to lower fluctuations. Additionally, the higher structural changes in protomers A to E of the CTT-binding regions when bound to beta5, compared to when bound to beta5-A+Y, are also confirmed by the higher RMSF values observed in the PL1 and PL2 regions of these protomers (See Figure S5). Importantly, compared to beta5, beta5-midpoint results in smaller RMSD values for the CTT-binding regions of all the protomers (Figure 2B). This indicates that shifting the negatively charged amino acids to the middle of the sequence is less consequential towards the stability of the PL1 and PL2 compared to switching the final amino acid in the sequence.

Focusing on PL3, the most differences between beta5 and the two constructs were observed in protomers A, C, and F (Figure 2C). Comparing beta5-A+Y with beta5, we found that PL3 of protomers A and C show significantly higher RMSD values when bound to beta5 and vice versa for PL3 of protomer F. When comparing beta5 with beta5-midpoint, the PL3 regions of protomers A through E exhibit lower RMSD values indicating less structural changes upon binding with beta5-midpoint. We then compared the CT-helix region and observed that protomers A, D, E, and F show significant structural differences between beta5 and beta5-A+Y. In contrast, the CT-helix regions of protomers B and C undergo similar changes with both beta5 and beta5-A+Y. When bound to beta5-midpoint, the CT-helix regions follow a similar pattern to PL3 where beta5-midpoint induces smaller structural re-arrangements than beta5 in protomers A through E (Figure 2D). In contrast, protomer F shows a higher RMSD for the CT-helix when bound to beta5-midpoint, indicating increased flexibility, which is confirmed by the RMSF analysis (Figures 2D and S5F). Also, another interesting observation is that the HBD region of protomers A, C, and E when bound to beta5-A+Y becomes more flexible (compared to the ATP-bound state) than when bound to beta5.

### B. The Essential Dynamics of the Katanin Spiral Conformation is Regulated by the CTT Isotypes

To uncover how the essential motions of katanin vary when bound to different CTT isotypes, we employed Principal Component Analysis (PCA) as described in the Methods section. For the spiral conformation of katanin, in the COMPLEX state, results show that beta5 and beta3 capture similar variance (55.22% and 52.84%, respectively) in the first two PCs, indicating comparable dominant motions, whereas beta4b captures less variance (44.42%), suggesting reduced coverage of the conformational space. However, in the ATP-free SUB state, beta4b and beta3 cover similar variance in the first two PCs, while beta5 covers a lower proportion of the conformational space. For the artificially constructed beta5 isotypes, beta5-A+Y and beta5-midpoint, we found similar conformational complexity in the COMPLEX state (50% variance), while in the SUB state beta5-midpoint and beta5-Cterm behave similarly and beta5-A+Y captures higher variance (63%), indicating that their dynamics are driven primarily by the dominant modes (Tables S7-S8).

The essential motions captured by PC1, when katanin is bound to beta5 in the COMPLEX state, reveal a swing-like movement of the HBD in protomer F shifting away from protomer A, with the NBD also moving horizontally away from A. Simultaneously, protomer A executes an up-and-down motion. These combined movements lead to the opening of the gap between the terminal protomers A and F, primarily driven by the larger horizontal displacement of the NBD of protomer F. When katanin is bound to beta4b, PC1 shows that the HBD of protomer F undergoes a more pronounced horizontal motion than in the presence of beta5, while the NBD exhibits a minimal motion. As shown in Figure 3B, although the porcupine arrows indicate that the HBD of protomer F is moving downward, this motion corresponds to it moving it away from protomer A, resulting in the opening of the gap between the terminal protomers.

**FIG. 3.**
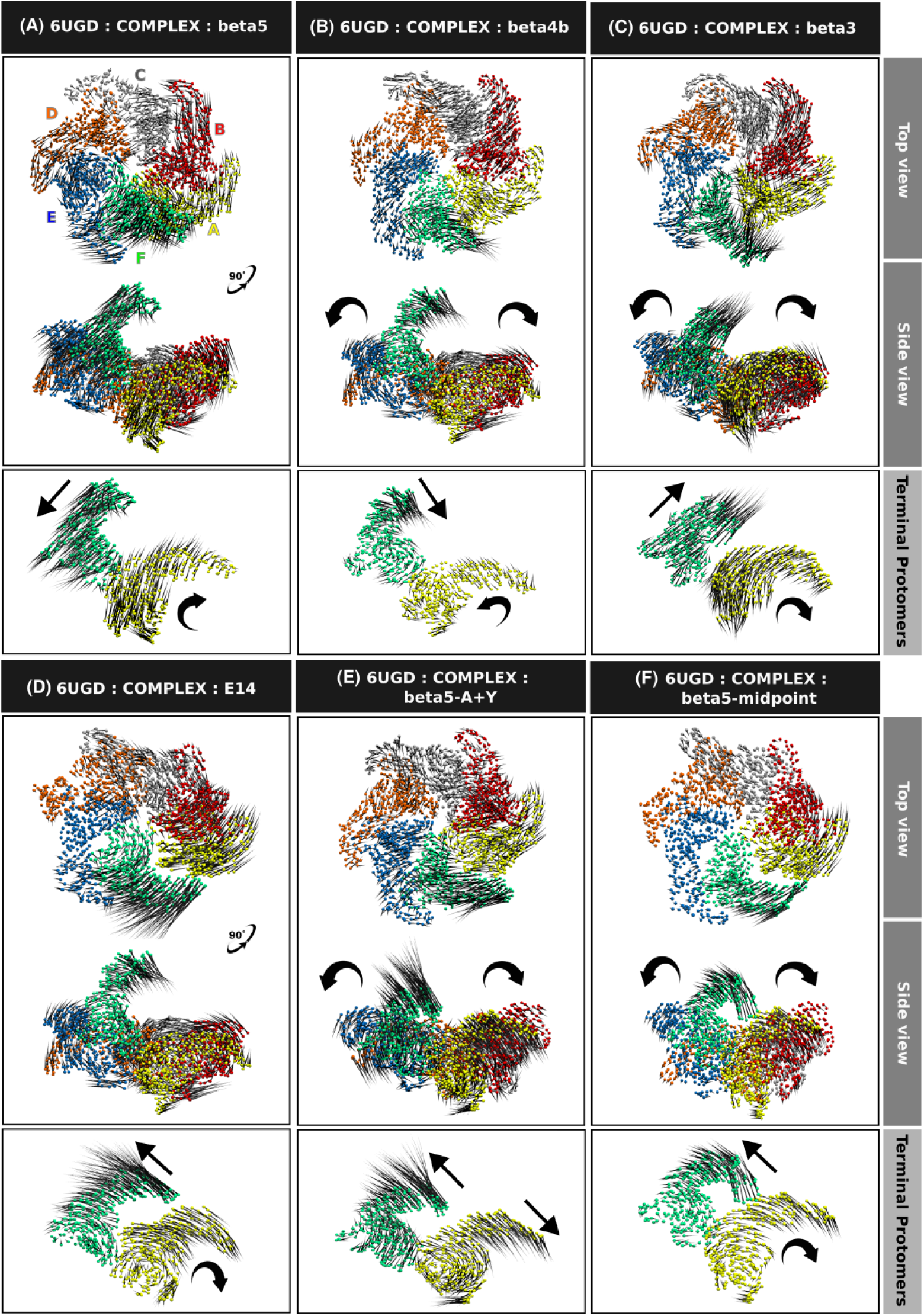
Porcupine plots illustrate the global motions corresponding to PC1 of the COMPLEX state of the katanin spiral conformation with different CTT isotypes. (A) beta5 (B) beta4b (C) beta3 (D) E14 (E) beta5-A+Y (F) beta5-midpoint. The variance covered by PC1 is in Tables S7-S8.

In this case the gap formation is driven primarily by the motion of the HBD of protomer F, unlike for katanin bound to beta5 where the NBD of F is involved. Interestingly, when bound to beta3, a poor target for katanin, PC1 shows a pronounced up-and-down movement of the HBD in protomer F, accompanied by an inward rotation in protomer A, but with a very small amplitude, in direct contrast to the motions in the presence of beta5. These motions result in a significantly larger gap opening between the terminal protomers A and F compared to the one seen in the presence of either beta5 or beta4b. These distinct motions suggest that katanin adopts a different dynamics when bound to beta3 than in the presence of good targets, supporting beta3’s unique behavior among natural CTTs as a poor target for katanin. Namely, they show a large opening of the gap between the protomers and the likely loss of the grip of the machine on the beta3 substrate. Importantly, this finding correlates well with the above results from the RMSD clustering that showed beta3 inducing the largest changes for protomer F both internally and with respect to its orientation vs the rest of the hexameric spiral. To better understand how natural CTT isotypes compare with the minimal substrate from the cryo-EM structure, we also examined the behavior of katanin bound to E14. We observed that the HBD of protomer F moves diagonally away from protomer A unlike our above findings for the natural CTT isotypes. We then quantified the subspace overlap between the natural CTT isotypes and the E14 bound system. All three natural CTT isotypes had similarly low overlap values, indicating weak similarity in their essential motions compared to the E14 minimal substrate (Table S11).

In the SUB state, where ATP is absent, the PC1 motions of katanin bound to beta5 show a swing-like movement of the HBD in protomer F moving away from protomer A, similar to the motion observed in the COMPLEX state with beta5. However, when bound to beta4b, the HBD of protomer F moves upward with high amplitude, while protomer A rotates outward, widening the gap between the terminal protomers in contrast to the horizontal motion seen in the COMPLEX state. When katanin is bound to beta3, PC1 motions indicate that the HBD of protomer F moves horizontally toward protomer A, while protomer A rotates inward, which is similar to its motion in PC1 of the COMPLEX state. These combined motions result in the closure of the gap between the terminal protomers, driven primarily by the motion of F, which is the opposite effect compared to the COMPLEX state. When bound to E14, we observe that the HBD of protomer F is moving away from protomer A, similar to the pattern seen with beta4b and in accord with the behavior seen above in the COMPLEX state. Therefore, beta5 and E14, which are both inhibitors, lead to similar motions in the COMPLEX and SUB states resulting in the opening of the gate between the end protomers. The non-inhibitor beta3 induces completely different motions in the COMPLEX and SUB states: a gate opening vs. a gate closing, respectively (See Figure S8).

Next, we evaluated the global dynamic behavior of the artificially constructed beta5 isotypes. In comparison to beta5, both beta5-A+Y and beta5-midpoint exhibited significantly different PC1 motions in the COMPLEX state. Specifically, the HBD of protomer F shifted diagonally upward away from protomer A, while protomer A rotated outward when bound to both the CTT constructs. The motion of the HBD in protomer F is similar to the PC1 motion observed when katanin is bound to E14. However the PC motions were less pronounced when katanin was bound to beta5-midpoint compared to beta5-A+Y. In the SUB state, all three beta5 constructs showed markedly different behaviors compared to beta5. When katanin was bound to beta5-A+Y, PC1 motions showed that the HBD of protomer F was moving vertically upward, while protomer A moved downward, resulting in an axial separation between the terminal protomers. When katanin was bound to beta5-midpoint, PC1 showed that the HBD of protomer F is moving diagonally away from protomer A, similar to the motion observed in beta4b, but with lower amplitude. When bound to beta5-Cterm, the HBD of protomer F moves horizontally away from protomer A, resembling the PC1 motion seen when katanin is bound to E14.

To further quantify how the motion patterns of the terminal protomers change upon binding different CTT isotypes, we calculated the center-of-mass (COM) distances between the terminal protomers, as described in the Methods section. This analysis was performed only for the COMPLEX state, as it represents the functional state of katanin during severing. By comparing the distributions of COM distances between the NBDs as well as the NBD of A to the HBD of F (shown in Figure S10), we found that both beta3 and beta5-A+Y exhibit higher KL divergence values compared to beta5, while the beta4b distributions have minimal divergence from the beta5 distributions. Consistent with these observations, the COM distance differences reported in Table S13 further demonstrate that variations between NBD–NBD and NBD–HBD distances are minimal for beta4b and beta5, but substantially larger for beta3 and beta5-A+Y. This indicates that the pattern of the motions for the terminal protomers changes between the experimentally observed inhibitors and non-inhibitors. Furthermore, we found that the distributions for beta5-midpoint follow the same trend as the non-inhibitors when compared to the beta5 distributions.

Next, we focused on the collective motions of the pore loops because the central pore serves as the active site for applying mechanical forces to the substrate, and these motions dynamically regulate the pore width, which is essential for substrate interaction. Following our previous work, we analyzed the PL1 motions along the top two principal components, which together account for more than 40% of the motion in the COMPLEX state and 50% in the SUB state across all CTT isotypes as listed in Tables S9 and S10^7^ . We found that, when katanin is bound to beta5, the main motions for PC1 in both the COMPLEX and SUB states involve a narrowing of the central pore, which suggests that beta5 has a strong grip on the hexamer, which is inhibitory. In the COMPLEX state, PC1 motions show PL1 of protomer A moving up, while PL1 of protomer F moving vertically down. PL1 in protomers B and C exhibits minimal movement, whereas protomers D and E display significant motions. In the SUB state, although a similar pore-closing motion is observed, the PL1 regions of the inner protomers show much larger displacements compared to the COMPLEX state, which we assign to the lack of ATP. However when katanin is bound to beta4b, we observe that the PL1 of both the terminal protomers moving in the same direction in both the states.

In the COMPLEX state, PC1 motions show a reduced movement of PL1 in the terminal protomers compared to beta5. Also, PL1 in protomers B and C shows increased motion while PL1 in protomers D and E exhibit much smaller displacements than those observed with beta5 (See Figure 4).

**FIG. 4.**
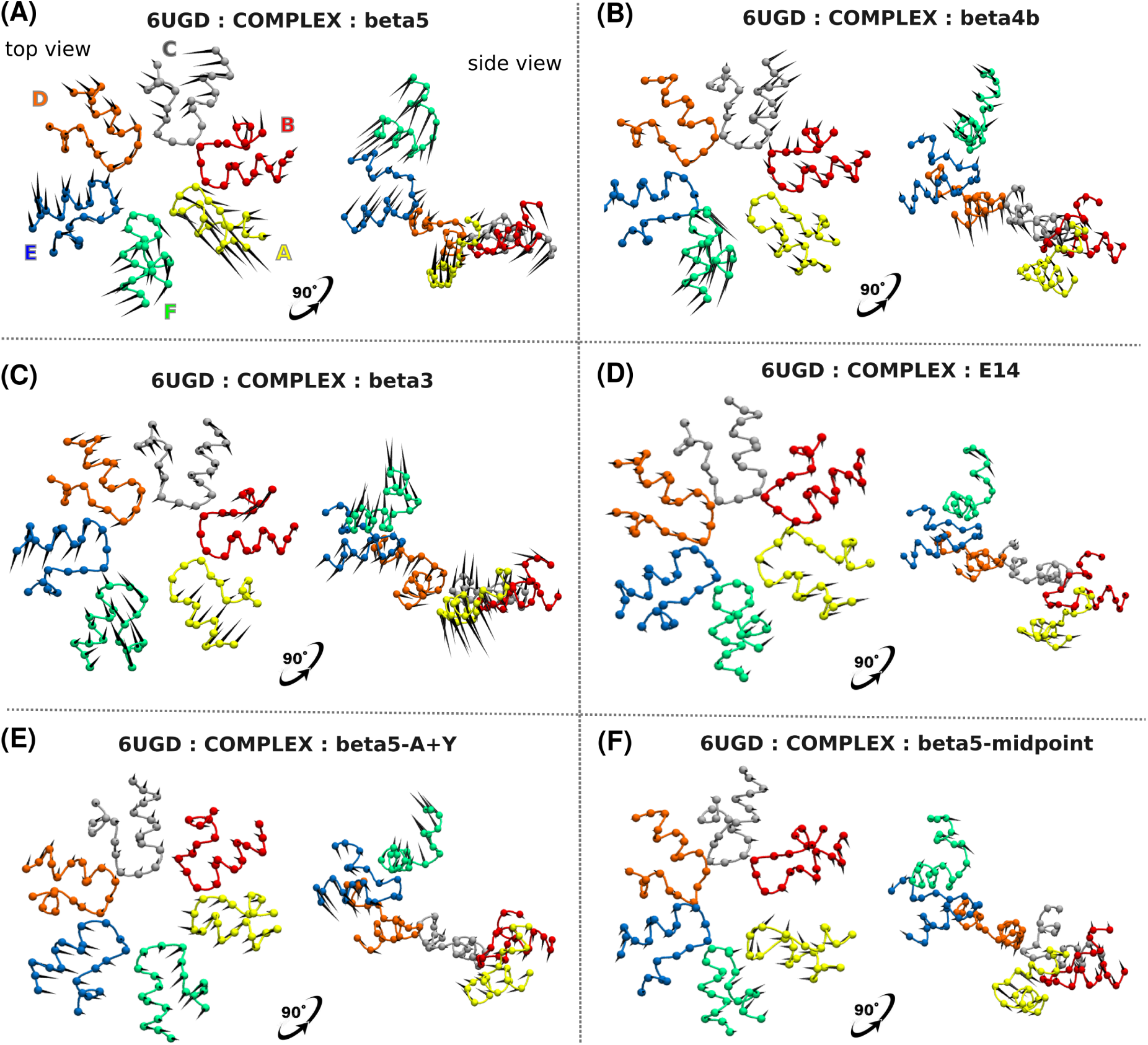
Porcupine plots illustrating the PL1 motions corresponding to PC1 of the COMPLEX state of the katanin spiral conformation with different CTT isotypes. (A) beta5 (B) beta4b (C) beta3 (D) E14 (E) beta5-A+Y (F) beta5-midpoint. The protomer labels, as indicated in (A), are used consistently across the entire figure. The variance covered by PC1 is in Tables S9-S10.

The presence of E14 arrests the motion of the pore loops in all protomers resulting in the maintenance of the original gap separation between the end protomers. Unlike beta5 and beta4b, katanin bound to beta3 shows different patterns of PL1 motion due to ATP binding.

In the COMPLEX state, PC1 indicates a widening of the pore, whereas in the SUB state lacking ATP, PC1 shows a narrowing of the pore, which recapitulates the motions discussed above for the full protomers. Thus, the non-inhibitor beta3 does not exert a strong grip on the pore loops like beta5. When comparing the PL1 motions in the SUB state, although both beta5 and beta3 lead to pore narrowing, the magnitude of PL1 motions across all protomers is reduced when bound to beta3 than to beta5. In the COMPLEX state as well, we observe reduced PL1 motion in the inner protomers compared to beta5, which is consistent with our earlier RMSD clustering results showing that the PL1 and PL2 loops of the inner protomers exhibit the lowest fluctuations in the presence of beta3, while displaying the largest changes in the presence of beta5. These results provide further evidence that, when katanin is bound to beta3, the motions of the PL1 regions display dynamics that are significantly different from those observed with high-affinity targets, highlighting the unique behavior of beta3 as a non-inhibitory substrate.

Comparing the PL1 motions of beta5 with those of the beta5 constructs, we observe that beta5-A+Y promotes pore widening in both the COMPLEX and SUB states. In contrast, when bound to beta5-midpoint, the system exhibits pore narrowing in the COMPLEX state and pore widening in the SUB state (Figure S9F). Notably, this pore narrowing differs from that observed with beta5, both in the direction of motion of the terminal protomers and in its magnitude, recapitulating the behavior seen in the presence of beta3. Similarly, as seen with beta3, both beta5-A+Y and beta5-midpoint show reduced PL1 motions in the inner protomers. When bound to beta5-Cterm, we observe pore expansion in the SUB state, with minimal PL1 motions of the interior protomers similar to the other constructs, but the PL1 motion of protomer A is significantly larger than that of the other constructs. Overall, all three beta5 constructs promote pore expansion with lack of coordinated and directional PL1 motions required for stable substrate engagement, unlike the behavior observed with beta5. These results suggest that redistribution of charge along the CTT disrupts the coordinated pore dynamics.

We also compared the overlap between the PL1 motions of the natural CTT isotypes and the cryo-EM E14 substrate. In the COMPLEX state, beta5 and beta3 exhibited similar RMSIP values, while beta4b showed a lower value. In the SUB state, all three CTT isotypes displayed comparable RMSIP values. Considering the beta5 constructs, in both the states, beta5-A+Y and beta5-midpoint exhibited comparable overlap with beta5. However, in the SUB state, beta5-Cterm showed a lower RMSIP, indicating that beta5-Cterm altered the native-like dynamics of PL1 seen in beta5 more than in the other two constructs.

### C. Salt Bridge Analysis Highlights Distinct Interaction Patterns Induced by Various CTT Isotypes

Salt bridges, formed between oppositely charged residues, are important for maintaining the stability of protein structures. As described in the Methods section, we performed a salt bridge analysis to identify any CTT-dependent inter- and intra-protomer salt bridge networks. Following our previous work, we focused primarily on the CTT-binding pore loop region to gain a broader understanding of how these interactions contribute to pore dynamics and interprotomer communication.

In the COMPLEX state, when katanin is bound to beta5, we observed two main interprotomer salt bridges, ARG275-ASP261 and ASP269-LYS265, formed between the PL1 regions of neighboring protomers. When bound to beta4b, the same two salt bridges were present. However, the persistence time of the ARG275-ASP261 salt bridge was shorter, while the ASP269-LYS265 salt bridge lasted longer compared to when bound to beta5. Additionally, a new salt bridge was observed between residues ASP171 in the fishhook and LYS265 in the PL1, extending the interaction from PL1 to the fishhook, a functional element specific to severing proteins that plays a role in hexamerization. These results suggest that binding to beta4b has changed the interprotomer communication network of the pore loop region of katanin compared to when beta5 is bound. Furthermore, when bound to beta3, the same two salt bridges as in beta5 were present, but, similar to the beta4b case, the ASP269-LYS265 salt bridge was more persistent than in the beta5. Also, we compared the inter-protomer salt bridges in the cryo-EM structure with the minimal E14 substrate. We found that the same two salt bridges observed in all the natural CTT isotypes were also present in E14, with persistence times similar to those in the beta5-bound case. In addition, an extra short-lived salt bridge was observed between PL2 residue ARG301 and WB residue GLU293. Thus, the long-lived inter-protomer salt bridges in the katanin pore of the spiral COMPLEX behave similarly in the presence of both inhibitory and non-inhibitory CTTs (See Figure 5).

**FIG. 5.**
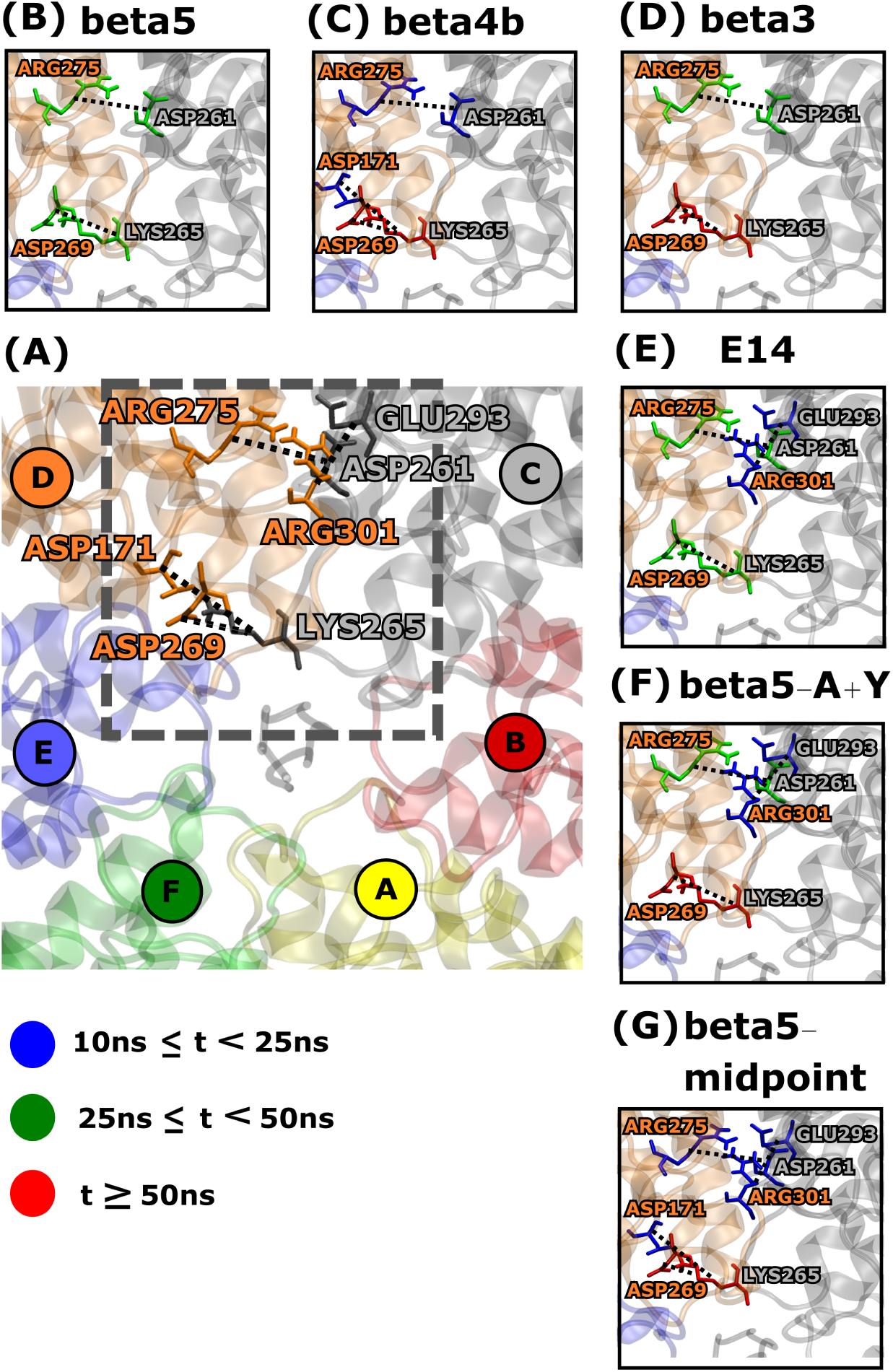
Interprotomer salt bridges observed for more than 10 ns in at least three protomer pairs of the katanin spiral conformation in the COMPLEX state. For visualization, the interface between protomers C and D is shown. (A) All interprotomer salt bridges. Residues in grey represent protomer i–1, and those in orange represent protomer i. Panels (B) – (G) show zoomed-in views of the interprotomer salt bridges specific to each CTT isotype. Colors (blue, green, red) indicate persistence time (t) as defined in the legend. If a residue forms multiple salt bridges, its color represents the salt bridge with the highest persistence time.

Compared to the COMPLEX state, in the SUB state we observed that the two common interprotomer salt bridges present across all CTT isotypes were still maintained, although their persistence times changed. Interestingly, additional salt bridges were observed in the SUB state when katanin was bound to beta4b and beta3 that were not present in the COMPLEX state. One such additional salt bridge was GLU308-LYS314, observed when bound to both beta4b and E14, which is located within the PL2 region, suggesting a broader role for PL2 in interprotomer stabilization under ATP-free conditions. The other additional salt bridge was ASP171-LYS265, which is present when bound to beta3, and had previously been observed in the beta4b COMPLEX system. Detailed information on the persistence times of these salt bridges can be found in Table S14. Therefore, unlike in the COMPLEX, the binding to beta3 has changed the interprotomer communication network of the pore loop region of katanin spiral SUB compared to when beta5 is bound. These two findings suggest that the katanin spiral is particularly stable when bound to beta5 as even the absence of ATP does not alter its inter-protomer salt-bridges. In contrast, when ATP is lost, additional salt-bridges need to form to ensure the integrity of the katanin spiral hexamer bound to beta3. Thus, the SUB state of the katanin spiral exhibits differences between the inhibitory and the non-inhibitory wild type CTTs.

Analyzing the intra protomer salt bridges, we identified two main salt bridge networks involving PL1, PL2, and PL3 in both the COMPLEX and SUB states. Importantly, these networks include complex salt bridges where certain charged residues form multiple interactions. As shown in Figures 6 and S12 , one salt bridge network is centered around residue LYS314 and the other around residue ARG312, both located in the PL2 region. The first salt bridge network involves four residues: ARG301, LYS314, GLU306 in PL2 and ASP346 in PL3 coupling PL2 to PL3. The other salt bridge network involves GLU308, ARG312 in PL2 and GLU271 in PL1 coupling PL2 to PL1. Apart from these two salt bridge networks, another salt bridge (GLU316 - ARG275) was observed in both states, coupling PL2 to PL1. Although these salt bridges were observed across all CTT isotypes, their persistence times varied depending on the specific isotype. In the COMPLEX state, when katanin is bound to beta5, we observed that the ASP346-LYS314 and GLU306-LYS314 salt bridges within the first network are long-lived, while the ASP346-ARG301 salt bridge is short-lived. However, when bound to either beta4b or beta3, the previously short-lived ASP346-ARG301 salt bridge becomes significantly more stable, whereas the ASP346-LYS314 salt bridge, which was long-lived in the beta5-bound system, becomes less stable. In the second salt bridge network, both the GLU308-ARG312 and GLU271-ARG312 salt bridges are short-lived when katanin is bound to beta5. When bound to either beta4b or beta3, the GLU308-ARG312 salt bridge remains short-lived, while the GLU271-ARG312 salt bridge no longer forms. These results indicate that the interactions between PL2 with PL1/PL3 found when katanin is bound to beta5 are affected by the binding of either beta4b or beta3 to katanin. We also found that the other independent salt bridge exhibited similar persistence times across all three natural CTT isotypes. In the SUB state, katanin bound to beta5 and beta3 showed comparable persistence times for the salt bridges within the two main networks. However, when bound to beta4b, the ASP346-ARG301 salt bridge was not present, indicating that, in the absence of ATP, this salt bridge is unstable, in contrast with the results in the COMPLEX state.

**FIG. 6.**
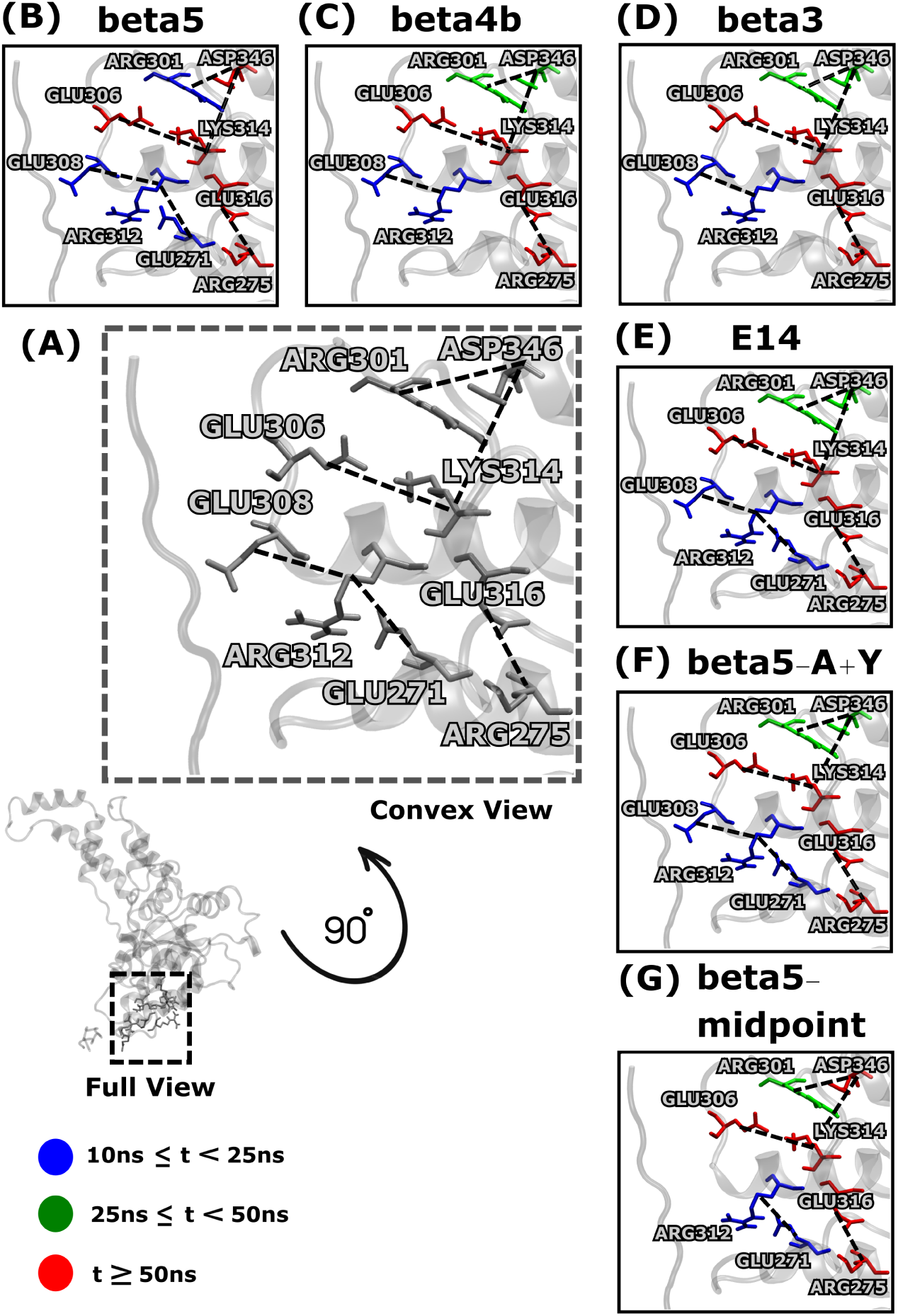
Intraprotomer salt bridges observed for more than 10 ns in at least three protomers of the katanin spiral conformation in the COMPLEX state. (A) All intra protomer salt bridges. The convex view is zoomed in to provide better visualization of the salt bridges. Panels (B) – (G) show zoomed-in views of the intra protomer salt bridges specific to each CTT isotype. Colors (blue, green, red) indicate persistence time (t) as defined in the legend. If a residue forms multiple salt bridges, its color represents the salt bridge with the highest persistence time.

We are now turning to the effect of the artificial beta5 constructs. In the COMPLEX state, beta5 displayed long-lived interprotomer salt bridges, including ASP261–ARG275 and ASP269–LYS265 within the PL1 region. Beta5-A+Y and beta5-midpoint preserved most of these interactions across several interfaces and maintained PL1–PL1 coordination and interfacial communication across the pore loops comparable to beta5.

Within the individual protomers, beta5 consistently formed the GLU316–ARG275 salt bridge in the COMPLEX state, which anchors PL2 to the adjacent convex interface. This intraprotomer interaction was also detected in beta5-A+Y and beta5-midpoint. Other PL2-related salt bridges, such as GLU308–ARG312 and GLU306–LYS314, were also observed in these constructs, further stabilizing the PL2 domain. ASP346–ARG301 and ASP346–LYS314 were found in beta5, beta5-A+Y, and beta5-midpoint. However, ASP346–LYS314 occurred across fewer protomers in beta5-A+Y and beta5-midpoint, suggesting a subtle loss in HBD stability. ASP269–LYS265 was also conserved among all three variants and contributed to both intra- and inter-chain coordination. These conserved salt bridge patterns in beta5-A+Y and beta5-midpoint support their capacity to maintain electrostatic integrity within individual protomers and across the entire hexamer in the ATP-bound state.

In the SUB state, beta5 exhibited a robust network of interprotomer salt bridges, particularly ASP261–ARG275 and ASP269–LYS265 across multiple chains, reflecting how the PL1 regions of each chain coordinate in the absence of ATP. These two key salt bridges were consistently observed in all three engineered isotypes: beta5-A+Y, beta5-midpoint, and beta5-cterm. This suggests that their formation is structurally favored and independent of the CTT charge redistribution. However, other stabilizing interactions such as GLU293–ARG301 and GLU308–LYS314 were absent in beta5-cterm but present in beta5 and the other two variants. This indicates that some important electrostatic connections near the pore loops are either missing or become weaker in beta5-cterm.

The intra protomer salt bridge network in the SUB state further supports these distinctions. Beta5 formed a recurring intra protomer salt bridge between GLU316 and ARG275, a long-lived interaction that links PL2 to the adjacent NBD region. This interaction was also present in beta5-A+Y, beta5-midpoint, and beta5-cterm, indicating that this internal structural feature is conserved across all variants. An additional PL2-related intra protomer salt bridge, GLU306–LYS314, was also shared among beta5, beta5-A+Y, and beta5-midpoint. However, in beta5-cterm this salt bridge was short-lived compared to the other beta5 variants and resembled the pattern observed in beta4b, which has moderate affinity to katanin. Other stabilizing salt bridges, including ASP346–ARG301 and ASP346–LYS314, were detected in all beta5 variants. Notably, ASP346–LYS314 persisted longer in beta5-cterm, similar to beta4b, while in beta5-A+Y and beta5-midpoint it was comparatively unstable, suggesting a loss of electrostatic rigidity along the upper HBD surface in those two variants. These findings indicate that although beta5-cterm lacks some key interprotomer salt bridges, it retains complementary internal salt bridges that help maintain structural cohesion.

Overall, beta5-A+Y and beta5-midpoint maintain most of the key salt bridges that characterize the native beta5 system, although with reduced intra protomer rigidity in certain regions. In contrast, beta5-cterm preserves only a subset of essential interactions in the SUB state and lacks several interprotomer salt bridges required for maintaining global pore-loop stability. This weakened connectivity likely contributes to altered pore behavior in beta5-cterm and may reduce the efficiency of force transmission during CTT engagement.

### D. Contact Map Analysis Reveals Distinct Binding Patterns of Inhibitory vs. Non-inhibitory CTTs with the Katanin Pore

Contacts between the CTTs and the central pore of katanin are crucial for its function, as they provide the mechanical grip needed for pulling and unfolding the substrate. These interactions help keep the substrate stable inside the pore, allowing katanin to apply the forces necessary to sever MTs. If these contacts are weakened or disrupted, katanin may not hold onto the substrate as effectively, which can shorten how long it stays attached to an MT and reduce its ability to carry out severing efficiently. To assess these interactions, we employed the contact map analysis from the Methods section, which quantitatively tracks CTT to pore contacts over the duration of the simulation for each system. Figures 7 and S13 show the contact maps obtained for the katanin COMPLEX system with the various CTT isotypes.

**FIG. 7.**
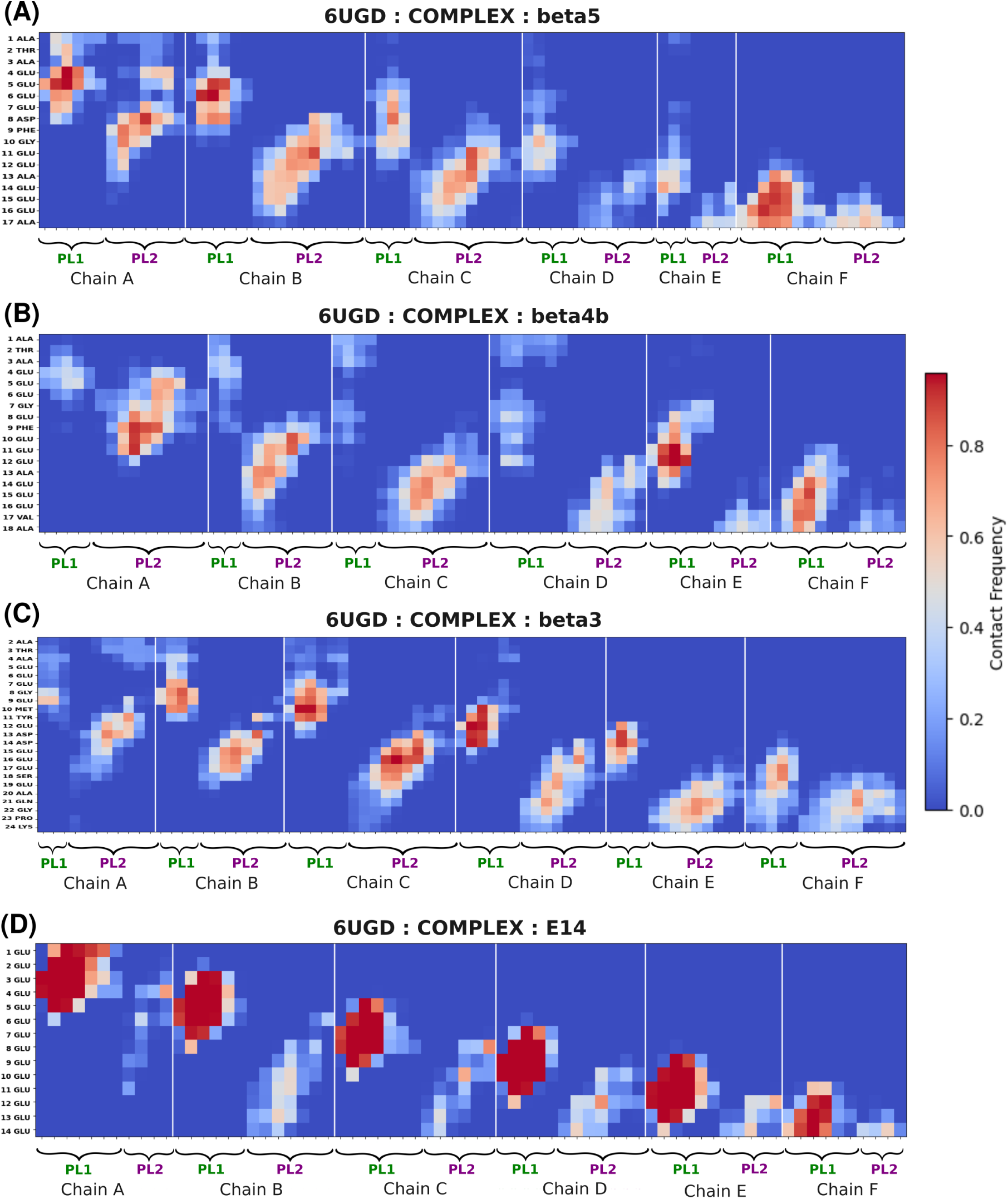
Contact maps showing the fraction of contacts between the natural CTT isotypes and the pore loop region (PL1 and PL2) of the katanin spiral conformation in the COMPLEX state. Here, red indicates that residues are highly in contact during the simulations and blue indicates less contacts. (A) Contact map with beta5 (B) Contact map with beta4b (C) Contact map with beta3 (D) Contact map with E14.

When katanin is bound to the beta5 CTT, we found strong and localized contacts with protomers A, B, C, and F. In protomers A and B, beta5 CTT formed strong contacts with both PL1 and PL2. In protomer C, these contacts were reduced compared to protomers A and B, while in protomer F the contacts were mostly with PL1.

When bound to the beta4b CTT isotype, the main contacts were made with the PL2 of protomers A, B, and C. There were no significant contacts with PL1, except in protomers E and F. In these two protomers, beta4b had more contacts with PL2. Overall, beta4b formed fewer contacts with the pore compared to beta5. Beta3 is forming contacts with both PL1 and PL2 of the internal protomers B, C, and D, but only minimal contacts with the pore loops of the terminal protomers, in striking contrast to beta5 which favors contacts with the terminal protomers.

As described above, katanin binding to the beta5 CTT isotype showed strong and localized contacts with chains A, B, C, and F, with notably high interaction frequencies with the PL1 of chains A, B, and F. These contacts were substantially persistent compared to the interactions observed in the beta5-A+Y complex. Notably, beta5-A+Y had reduced contact frequencies with both PL1 and PL2 of the terminal protomers, similar to the behavior of beta3, which helps explain its experimentally observed lower binding affinity for katanin. Instead, beta5-A+Y engaged more frequently with inner protomers, particularly the PL regions of chains C and E, indicating a shift in substrate engagement toward the internal regions of the hexamer. This finding aligns well with the above results from the RMSD clustering that showed that the PL1 and PL2 regions of the central protomers have the lowest fluctuations in the presence of beta3 and beta5-A+Y, which we can ascribe to the increase in the contacts with these non-inhibitory CTTs. In contrast, RMSD clustering showed that PL1 and PL2 of the central protomers have the largest fluctuations in the presence of beta5, which is likely due to the reduced contacts with this inhibitory CTT.

A similar interaction pattern as the one seen with the non-inhibitory CTTs was present in the beta5-midpoint construct. While wild-type beta5 preferentially contacted the pore loops of terminal protomers, especially PL1, beta5-midpoint showed enhanced contacts with PL2 of chain B and PL1 of chain D, both of which are inner protomers. In contrast, the other pore loops across the hexamer had only limited interactions with beta5-midpoint. These findings suggest that redistributing the negatively charged residues away from the wild-type C-terminal configuration alters the spatial engagement of the CTT with katanin, favoring inner protomer binding over terminal pore loop contacts. In summary, our results show that the inhibitory CTTs, such as beta5 and beta4b, prefer to make strong contacts with the pore loops of the terminal protomers while the non-inhibitory CTTs and beta5-midpoint all favor contacts with the pore loops of the internal protomers.

### E. Katanin rings from different species show distinct behaviors when bound to the beta5 isotype

We compared the ring structures of katanin from *C. elegans* and human when bound to beta5 to determine their behavior with the same CTT isotype and to identify potential differences that limit the non-inhibiting CTT isotypes from docking with the *C. elegans* katanin ring. First, we analyzed the essential motions using principal component analysis to establish the patterns of motion of the two structures.

PC1 motions of the *C. elegans* katanin ring conformation bound to beta5 show that the NBDs of both terminal protomers move toward each other, while the HBD of protomer F exhibits an up and down motion and the HBD of protomer A rotates outward. The inner protomers undergo a coordinated, butterfly-like pushing down motion, suggesting axial excursions and compression that likely facilitate substrate translocation. However, when beta5 is bound to the human katanin ring, although protomers A and F move toward each other, similar to what we found in the *C. elegans* ring, the motion of the HBD of protomer F and the disordered movement of the HBD of protomer A and of the inner protomers differ from the butterfly-like motion observed in the *C. elegans* ring. The dominant motion of protomer F is similar to that in the *C. elegans* ring, but protomer A and the inner protomers have uncorrelated motions along PC1, with relatively small displacements (See Figures S14A and S14B). The COM distributions further quantify these results indicating that, for both the distances between the two NBDs and between the NBD and HBD of the terminal protomers, the *C. elegans* and human rings have high KL divergence values, indicative of differences in their motions (Figure S16).

At the pore loop level, PC1 motions of the *C. elegans* katanin ring show that PL1 of protomer A is moving away from the other protomers, while the pore loops of the remaining protomers are moving together in a coordinated manner. These collective motions of the inner protomers suggest modulation of the pore width while maintaining an organized loop behavior. However, due the displacement of PL1 of protomer A, the pore opens widely. In the human ring, PL1 of protomer A also moves away from the neighboring protomers, but the motions of PL1s of the other protomers are different: the PL1 loops of protomers E and F move together, while the pore loops of protomers B, C, and D move together. This grouped pattern differs from the coordinated motions observed in the C. elegans ring, which is indicative of a different dynamic behavior in the pore loop region, where the substrate binds, which is species-specific (See Figures S15A, and S15B).

Next, we analyzed the contacts between the central pore and beta5 to identify any species dependent changes in the interactions. As shown in Figure 8, when beta5 binds to the *C. elegans* katanin ring, the inner protomers make more contacts with beta5 compared to when beta5 binds to the human katanin ring. Specifically, the PL regions of inner protomers C and D form strong contacts in the *C. elegans* ring, which are much weaker in the human ring. In contrast, contacts between the pore loop regions of the protomers A and B are similar in both species. In the human ring the contacts between protomers E and F and beta5 are stronger than in the *C. elegans* case, which supports the above finding that the pore loops of these protomers move together in the human ring. These observations highlight species-specific differences in how the central pore interacts with the substrate, with the inner protomers playing a more prominent role in the *C. elegans* katanin ring conformation.

**FIG. 8.**
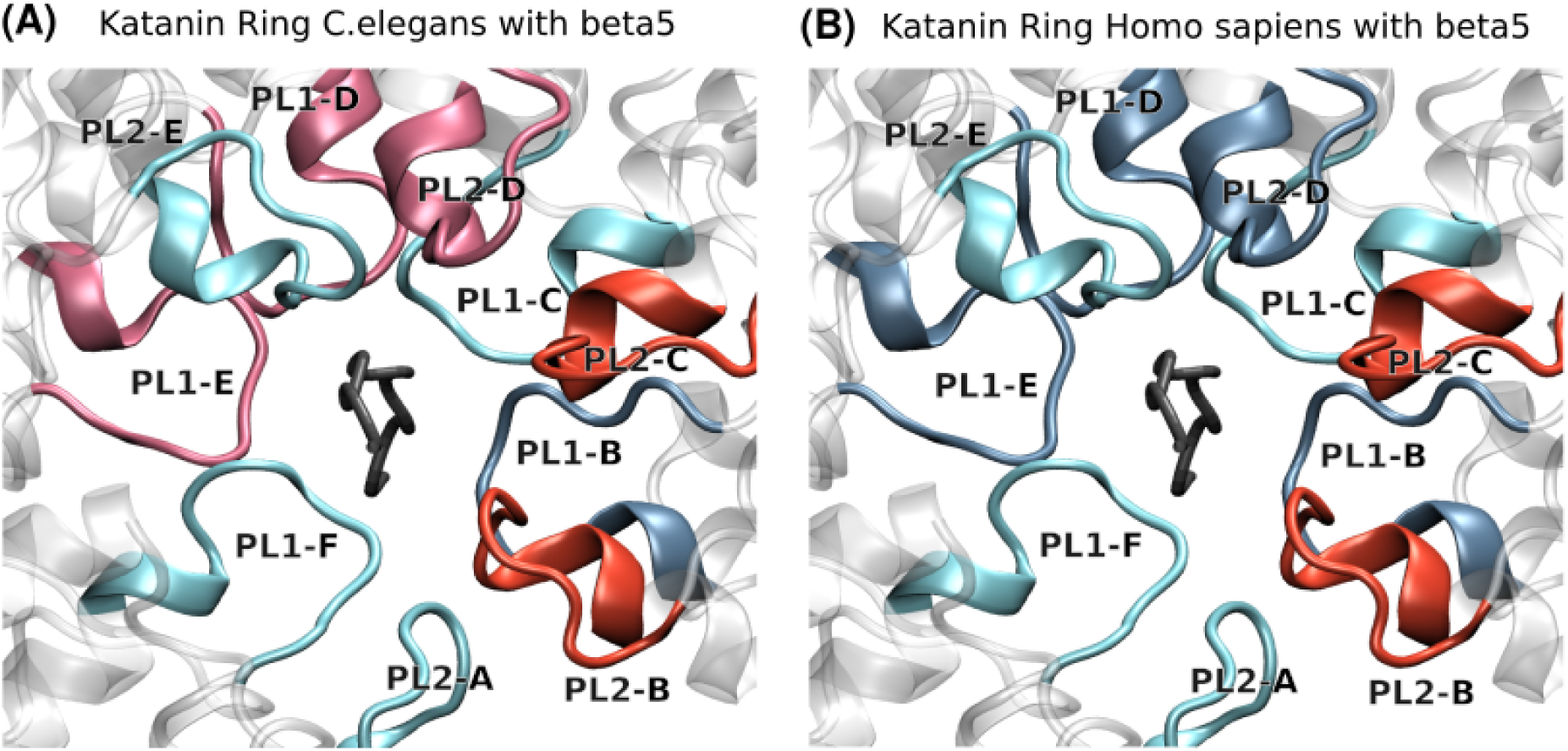
Contacts between the central pore and beta5 CTT in the katanin conformation in different species. (A) Katanin ring C.elegans with beta5 (B) Katanin ring Human with beta5. Here red indicates stronger contacts and blue indicates weaker contacts.

Finally, we analyzed the salt bridge networks to determine how the electrostatic interactions vary between species. In the *C. elegans*, we identified three significant inter-protomer salt bridges, namely ASP261-ARG275 and ASP269-LYS265 between the PL1 regions and GLU308-LYS314 between the PL2 regions. In comparison, the human ring system exhibited only one significant inter-protomer salt bridge (GLU285-LYS281) in the PL1 region, which was highly unstable compared to those observed in the *C. elegans*. These results further confirm that, in the presence of beta5, the pore loops lining the central pore in the *C. elegans* katanin ring are more strongly connected with each other and execute more coordinated motions than the loops in the human ring.

To further understand the species-specific differences in the dynamics and interaction patterns for the same CTT isotype, we performed a sequence alignment between the *C. elegans* and human katanin, specifically focusing on the pore loops and the CT-helix, the element important for the oligomerization of katanin. Although sequence differences are present in PL1 and PL3, most substitutions are conservative and preserve the chemical nature of the residues. In contrast, in human katanin the PL2 sequence contains more charged residues, and the CT-helix region shows a higher content of both hydrophobic and charged residues. Since PL2 is involved in interprotomer interactions and the CT-helix contributes to oligomerization, these differences may have an impact on how the hexameric structure is maintained. Consistent with these findings, the human katanin ring shows a tendency to form less stable assemblies, including dimers or trimers during dynamic motions, whereas the *C. elegans* ring remains intact in the presence of beta5. These observations suggest that sequence differences in regions important for interprotomer interactions may contribute to the distinct stability and dynamic behavior observed between the katanin rings of the two species.

### F. Human Katanin Ring Conformation Bound to Non-Inhibitory CTT isotypes

To investigate how the experimentally determined non-inhibitory CTT isotypes interact with the human katanin ring, we analyzed the essential dynamics and the interactions formed at the central pore for two CTT isotypes, beta3 and beta5-A+Y. Notably, these CTTs failed to dock into the *C. elegans* katanin ring, highlighting a species-specific difference in substrate recognition. PC analysis shows that, when human katanin is bound to beta3, the NBDs of the terminal protomers A and F move in the same direction, while the HBD of protomer F exhibits an up-and-down motion and the HBD of protomer A executes a horizontal movement. This motion pattern is distinct from that seen above in the presence of beta5, and also the overall motion occurs with less magnitude compared to the beta5-bound system. When katanin is bound to beta5-A+Y, PC1 reveals a different dynamic pattern. The HBD of protomer A moves downward while its NBD moves horizontally toward protomer F. At the same time, the HBD of protomer F moves vertically upward. This opposing vertical displacement between protomers suggests a loss of balanced axial coordination resulting in a stationary state where the ring gate neither opens nor closes.

At the pore loop level, when human katanin is bound to beta3, PC1 shows that the separation of the inner protomers is still observed, similar to that seen before with beta5. However, the coordination between protomers E and F is weaker compared to the beta5-bound system. When bound to beta5-A+Y PC1 motions show that PL1 of protomer F moves upward and horizontally, but without being accompanied by a coordinated motion across the other pore loops. Compared to beta5, these motions appear less synchronized and more asymmetric, suggesting weakened mechanical coupling within the pore in the presence of a non-inhibitor.

Analysis of contacts at the central pore revealed that, when the human katanin ring is bound to beta3, the substrate interacts more extensively with the PL1 than the PL2 loops of the inner protomers, and the PL2 of protomer F. This pattern differs from that observed with beta5, where only minimal contacts occur with these regions. Thus, the non-inhibitor beta3 is more strongly bound to the ring than the inhibitor beta5. When comparing beta5-A+Y with beta5, a similar contact pattern was observed for chain A. PL1 of chains E and F showed moderate contacts with beta5-A+Y, which are comparable to the beta5-bound human katanin ring. However, there are several notable differences: PL1 of chains B, C, and D contacted beta5-A+Y more frequently than beta5, indicating a shift in the interaction with the CTT preference toward more central protomers. Conversely, PL2 of chain B exhibited reduced contact frequency in the beta5-A+Y system compared to the beta5-bound complex. Therefore our results show that the artificial non-inhibitor beta5-A+Y behaves more similarly to beta3, the wild-type non-inhibitor, in the fact that it makes stronger contacts with the pore loops of the inner protomers than what we’ve found with beta5, while having only short-lived contacts with the terminal protomers. Furthermore, this recalls the behavior of the central pore of the spiral state in the presence of the non-inhibitors versus the inhibitors.

To understand how electrostatic interactions support CTT binding to the human katanin ring, we analyzed the salt bridges formed within the pore-loop regions in the human katanin ring conformation. In the beta5-bound human ring structure, we observed a well-connected network of intraprotomer salt bridges linking PL1 and PL2, including GLU287–ARG328 and GLU332–ARG291, the latter being a long-lived salt bridge. These interactions connect the two pore-loop regions and contribute to structural coordination within individual protomers. In addition, interactions between PL2 residues further stabilize the loop region, forming a salt-bridge network involving GLU322 with LYS330 and ARG327. Importantly, interprotomer salt bridges within the PL1 region are also present, indicating communication between neighboring protomers. Together, these salt bridges indicate coordinated electrostatic connections between PL1 and PL2, supporting stable pore organization when bound to beta5.

In contrast, when the human katanin ring is bound to either beta3 or beta5-A+Y, the overall salt-bridge pattern is altered. Although certain PL2 intra-protomer interactions such as GLU322–LYS330 remain present, the PL1–PL2 interactions observed in beta5 are weakened or absent; therefore, the salt-bridge network centered on GLU322 is not fully maintained. This difference suggests reduced electrostatic communication between the two pore loops within individual protomers. However, the inter-protomer connectivity within PL1 is long-lived in the human ring katanin bound to beta5-A+Y. While some local stabilization and inter-protomer connections are retained, the loss of coordinated PL1–PL2 interactions likely reduces the structural integration across the pore, which aligns well with our above findings from the essential dynamics.

## IV. CONCLUSIONS

The importance of the beta-CTT sequences and their regulatory behavior in MT severing by katanin has been experimentally recognized. However, a molecular-level understanding of how variations in the CTT sequence control substrate binding to katanin, pore engagement, and why these sequences show different behaviors toward human and *C. elegans* katanin were unclear. Specifically, in an experimental system containing katanin, MTs, and free carboxy-terminal tail sequences added one at a time, beta5, beta4b, and poly-E free CTT tails act as inhibitors, showing an enhanced tendency to bind to katanin and prevent MT severing. On the other hand, beta5-A+Y and beta3 show non-inhibitory behavior, meaning they are less likely to bind to katanin in solution, thus allowing katanin to bind and sever MTs. In this study, we combined docking, molecular dynamics simulations, interactions analysis, and dynamical analysis to address this question. We examined both natural and engineered CTTs in the spiral and ring states of katanin from *C. elegans* and human, focusing on how they interact with the pore loops, the pattern of salt bridges formed with the pore loops, and how these interactions affect the overall motion of the hexamer.

We found that CTT binding strongly depends on both the sequence of the tail and the structural state of katanin. In the spiral state, most CTTs can dock easily to the *C. elegans* spiral katanin, suggesting that this state is more flexible and less selective. The only type of beta CTT that doesn’t dock into this state is the beta5-Nterm, an artificial construct of the beta5 CTT with all the charges grouped at its N-terminal end. Intriguingly, the ring state behaves very differently: the binding becomes more selective toward katanin from the same species. Non-inhibitory CTTs, such as beta3 and beta5-A+Y, do not dock to the *C. elegans* ring, but they dock to the human ring. In contrast, inhibitory CTTs, such as beta5, beta4b, E14, and the inhibitor-like CTT beta5-Cterm bind to both species. This shows that it is not just the overall charge of the CTT that matters, but also where the charged residues are located along the tail to match the arrangement of the pore loops in the ring state.

When comparing the contacts between the *C. elegans* ring and the human ring in complex with beta5, we observed that the electronegative residues located in the latter part of the sequence form strong contacts with the PL2 region of the interior protomers (B, C, and D) in the *C. elegans* ring. However, in the human ring, these strong contacts are observed only in protomer B and are replaced by contacts with both PL1 and PL2 of terminal protomers E and F. We also found that the electronegative residues located in the middle part of the sequence form strong contacts with PL1 in protomer B. This suggests that, although the same CTT is involved in binding, the specific interactions between the CTT and the various protomers are different in the human vs. the *C. elegans* katanin, in that beta5 forms strong contacts with both central pore loops (PL1 and PL2) of the terminal protomers (B, E, and F) in human katanin.

When examining the non-inhibitors beta3 and beta5-A+Y, we found that the strong interactions between the PL2 region of the interior protomers in the human ring and the electronegative residues in the latter part of the inhibitory CTT sequences are no longer formed. Also, the strong contacts observed with the PL2 region of protomer B when bound to beta5 are significantly weakened for the non-inhibitors. The absence or weakening of these interactions may explain why the non-inhibitors fail to dock with the *C. elegans* ring. Overall, these results indicate that strong contacts between the electronegative runs at the C-terminal end of a CTT and the PL2 region of the interior protomers are critical for docking in the *C. elegans* ring, whereas the human ring appears to be less dependent on such interactions, relying instead on contacts made by the runs of electronegative residues throughout a CTT with both central pore loops of the terminal protomers. Also of note is the difference between beta5 and beta5-A+Y when bound to the human ring state: the electronegative part at the C-terminal end of beta5 makes strong contacts with both pore loops of protomers E and F, while for beta5-A+Y only contacts with PL1 survive, and even those are weakened compared to beta5. This behavior strongly suggests that the presence in beta5-A+Y of the bulky amino acid tyrosine (Y), as opposed to the small amino acid alanine (A) in beta5, is responsible for the loss of contacts with the PL2 loops. Based on this, we propose that beta5-A+G would be an inhibitor. Keeping in mind the selectivity of the ring for non-inhibitors, we posit that beta5-A+G would bind to both the *C. elegans* and the human ring. This is indeed what we found using the docking approach described in the Methods (see Fig. S21). Finally, our above finding about the PL2 contacts suggests that strong contacts between a CTT and the PL2 loops of both terminal protomers E and F are a characteristic of a katanin inhibitor.

Beyond these specific interaction patterns in the ring state, when we examined the spiral state of katanin, we found a clear difference in how inhibitory and non-inhibitory CTTs engage with the pore loops. Inhibitory CTTs form strong and stable contacts with the PL1 regions of terminal protomers, especially chains A, B, and F. At the same time, they also maintain stable interactions with PL2. These interactions are supported by long-lived salt bridges, both inter- and intra-protomer, which contribute to the stabilization of the substrate within the pore. In contrast, non-inhibitory CTTs do not show this extensive interaction pattern. Instead, their contacts are concentrated toward the inner protomers, and their salt bridges are weaker and less stable, especially in the PL2 regions. This leads to a less stable and less coordinated binding inside the pore.

The RMSD-based double clustering analysis further showed that different CTTs reshape different parts of the spiral katanin. Among the natural CTT isotypes, beta5 showed the largest global structural deviation relative to the ATP-bound state and induced the strongest rearrangements in the inner protomers, especially protomers C and D, which act as structural scaffolds. In contrast, beta3 produced the largest structural deviations in the terminal protomer F. At the level of the pore loops, beta5 caused larger changes in the PL1 and PL2 of the inner protomers, while beta3 produced smaller changes there and instead shifted the structural response toward the terminal region. The engineered constructs beta5-A+Y and beta5-midpoint also followed the same overall pattern as beta3 rather than beta5, showing reduced structural changes in the inner protomers but considerable deviations in protomer F. Together, these results indicate that inhibitory and non-inhibitory CTTs reshape the structure of the spiral katanin in fundamentally different ways.

These differences at the interaction level directly affect the essential dynamics of katanin. When inhibitory CTTs are bound to the spiral katanin, the terminal protomers show coordinated movements leading to the narrowing of the central pore. However, when non-inhibitory CTTs are bound, this coordinated motion is reduced. The movements become weaker and less organized, with terminal protomers exhibiting altered displacement patterns and the inner protomers contributing less effectively to the collective dynamics. Specifically, the loss of coordinated relative movements between the end protomers A and F, along with changes in the motions of their HBD and NBD domains, result in the opening of the pore. It is likely that the opening of the pore seen in the presence of non-inhibitory CTTs frees katanin to bind to the microtubule and sever it. In contrast, the strong grip exerted by inhibitory CTTs, which results in the narrowing of the central pore, arrests the hexamer on the free CTT thus preventing microtubule severing.

In summary, our results show that the binding of katanin to a CTT sequence depends not only on the overall charge but more importantly on how the charges are distributed along the sequence. This distribution determines how well the CTT can interact with pore loops, how stable those interactions are, and whether the katanin hexamer can maintain a coordinated motion between its protomers. Even the one amino acid in the C-terminal end of a sequence can affect the substrate recognition by katanin. In addition, the species-specific differences observed in the ring state suggest that even subtle structural differences between human and *C. elegans* katanin can strongly influence substrate recognition. Overall, by connecting sequence-level features to structural behavior and functional outcomes, this work helps explain the origin of inhibitory and non-inhibitory behavior in CTTs. These findings also suggest that it may be possible to design modified CTTs or peptide-based regulators that control katanin activity by targeting specific pore interactions and dynamic features. This will provide new directions for studying and potentially modulating MT severing.

## V. SUPPLEMENTARY MATERIAL

The supplementary material contains additional methods and analyses from the results: Tables S1 and S2: Summary of the MD simulation setups. Tables S3 and S4: Global average RMSD values. Table S5: Clustering cutoff analysis used to determine the optimal RMSD cutoff for the double clustering procedure. Table S6: List of functional residue regions in *C. elegans* and human katanin. Tables S7–S10: PCA variance coverage for global and PL1 motions. Tables S11 and S12: RMSIP comparisons between different setups and the beta5-bound system. Table S13: Center-of-mass distance changes between terminal protomers. Tables S14–S17: Inter- and intra-protomer salt bridge interactions in spiral conformations. Tables S18–S20: PCA and RMSIP analyses for ring conformations. Tables S21 and S22: Salt bridge interactions in ring conformations. Figures S1–S3: RMSD vs time plots for spiral and ring katanin systems. Figures S4 and S5: RMSF difference analyses of spiral katanin highlighting functional regions. Figures S6 and S7: RMSD comparisons of cluster 1 central structures for katanin spiral COMPLEX systems relative to the ATP-only state. Figures S8 and S9: PC1 porcupine plots of the global and pore-loop motions of the katanin spiral SUB state with different CTT isotypes. Figure S10: COM distance distributions and KL values between the terminal protomer regions. Figures S11 and S12: Inter- and intra-protomer salt bridge interactions. Figure S13: Contact maps between the CTTs and the pore-loop regions. Figures S14 and S15: PC1 porcupine plots of the global and PL1 motions of the katanin ring. Figure S16: COM distance distributions and KL values between *C. elegans* and human katanin ring conformations. Figure S17: Contact maps of katanin ring conformations. Figures S18 and S19: Inter- and intra-protomer salt bridge interactions in the katanin ring. Figure S20: Sequence alignments between *C. elegans* and human katanin rings. Figure S21: Docked katanin ring structures with CTT isotypes in *C. elegans* and human systems.

## ACKNOWLEDGMENTS

We thank Maria S. Kelly for their help with setting up the GROMACS simulations for the beta5-katanin spiral hexamer. This research was funded by the National Science Foundation (NSF) through grants MCB-1817948 and MCB-2527485 (both to RID). This work used the ACCESS program’s advanced computational resources through the allocation BIO210094 (to RID).

## AUTHOR DECLARATIONS

### Author Contributions

Madhavie Ranpati Dewage: Data curation (equal); Formal analysis (equal); Investigation (equal); Methodology (equal); Software (equal); Validation (equal); Visualization (equal); Writing – original draft (equal). Shehani Kahawatte: Data curation (equal); Formal analysis (equal); Investigation (equal); Methodology (equal); Software (equal); Validation (equal); Visualization (equal); Writing – original draft (equal). Jennifer L. Ross: Conceptualization (supporting); Writing – original draft (supporting). Ruxandra I. Dima: Conceptualization (lead); Data curation (equal); Formal analysis (equal); Funding acquisition (lead); Investigation (equal); Methodology (equal); Project administration (lead); Resources (lead); Supervision (lead); Validation (equal); Writing – original draft (equal).

## DATA AVAILABILITY

The data that support the findings of this study are available from the corresponding author upon reasonable request.

