## Supplementary Material for "Distinct Tubulin C-Terminal Tails Control the Efficiency of a Microtubule Severing Machine"

### Supplemental Information for Distinct Tubulin C-Terminal Tails Control the Efficiency of a Microtubule Severing Machine

*Authors: Madhavie Ranpati Dewage[1#], Shehani Kahawatte[1#], Jennifer L. Ross[2], Ruxandra I. Dima[1]**

***Affiliations**:*

1. *Department of Chemistry, University of Cincinnati, Cincinnati, Ohio 45221, United States*
2. *Department of Physics, Syracuse University, Syracuse, NY, United States*

**Corresponding author: Ruxandra I. Dima, Department of Chemistry, University of Cincinnati, Cincinnati, Ohio 45221.*

**#** *Authors with equal contribution*

### RMSD Analysis

We utilized the backbone RMSD of katanin with respect to the initial structure of each setup with different CTT isotypes to evaluate how the overall stability of katanin changes upon binding to various CTT isotypes. When comparing the global average RMSD of natural CTTs bound to katanin spiral conformation in the COMPLEX state (with ATP), we observed that katanin bound to the beta3 CTT exhibits reduced flexibility compared to beta5 and beta4b. Also, katanin bound to either beta5 or beta4b appears to be more flexible compared to E14, which serves as the substrate in the cryo-EM structure (Table S3 and Figure S1). When ATP is removed (SUB state), katanin becomes more flexible when bound to all natural CTT isotypes compared to the COMPLEX state. In this ATP-free state, beta4b, beta3, and E14 show similar high global average RMSD values, while beta5 maintains a global average RMSD value similar to the COMPLEX state (Table S3 and Figure S2). These results indicate that katanin becomes more flexible in the absence of ATP, regardless of the CTT isotype bound to it. However, beta5 CTT, which is an experimentally validated good target for katanin, helps stabilize the structure more effectively than other isotypes.[1] When comparing the stability of the katanin spiral hexamer in the presence of artificially constructed beta5 variants, in both states (COMPLEX and SUB), beta5-A+Y and beta5-midpoint have average RMSD values between 6–7 Å, which are close to the average RMSD of beta5 (6.15 Å). This indicates that beta5-A+Y and beta5-midpoint have a similar stability to the native beta5 CTT isotype (Table S3 and Figures S1-S2). However, beta5-cterm bound to katanin leads to the highest average RMSD of katanin among the beta5 variants. The ring conformation exhibits lower global average RMSD values compared to the spiral conformation, indicating greater structural stability. When comparing the ring conformation of katanin between the two species, *C. elegans* and *Homo sapiens*, we found that the beta5 CTT bound *Homo sapiens* katanin ring is more stable than the corresponding *C. elegans* ring.

### Equations

**Equation S1.** The equation used to calculate the root mean square fluctuation (RMSF) for each MD trajectory. Here $r_i(t_j)$ is the position of C-alpha atom of the $i^{th}$ residue at time $t_j$. $r_i^{ref}$ is the position of the $i^{th}$ residue of the reference structure. $N_{frames}$ is the total number of frames of each MD trajectory.

$$RMSF_i = \sqrt{\frac{1}{N_{frames}} \sum_{t_j=1}^{N_{frames}} |r_i(t_j) - r_i^{ref}|^2}$$

**Equation S2.** The equation used to average the RMSF values over the trajectories. $< RMSF_i >$ is the average RMSF of the i^th residue. $n_{traj}$ is the total number of trajectories.

$$< RMSF_i > \quad = \frac{1}{n_{traj}} \sum_{k=1}^{traj} (RMSF_i)_k$$

**Equation S3.** The root-mean-square inner product (RMSIP) used to measure the overlap between the extracted subspaces from the essential dynamics analysis (PCA). Here, the eigenvectors of the two subspaces, A and B are $\eta_i^A$ and $\nu_i^B$. The top 10 eigenvalues in the WT were found to cover around 90% of the overall variance so we set J to 10.[2]

$$RMSIP \; = \left(\frac{1}{J} \sum_{i,j=1}^{J} \left(\eta_i^A \bullet \nu_i^B\right)\right)^{1/2}$$

### Tables

**Table S1.** Summary of the MD setups for the Katanin Spiral conformation in various states

| State | Nucleotide | Substrate | #Atoms | #Residues | #Water | #Na | #Trajectories | Total Simulation time (ns) |
|---|---|---|---|---|---|---|---|---|
| COMPLEX | ATP | beta5 | 403019 | 1919 | 127888 | 70 | 5 (200ns) | 1000 |
| | ATP | beta4b | 282965 | 1920 | 87867 | 70 | 5 (200ns) | 1000 |
| | ATP | beta3 | 282924 | 1925 | 87837 | 70 | 5 (200ns) | 1000 |
| | ATP | beta5-A+Y | 282857 | 1919 | 87830 | 70 | 5 (200ns) | 1000 |
| | ATP | beta5_midpoint | 282935 | 1919 | 87860 | 70 | 5 (250ns) | 1250 |
| | ATP | E14 | 419068 | 1916 | 133241 | 74 | 5 (200ns) | 1000 |
| SUB | - | beta5 | 283040 | 1919 | 87989 | 52 | 5 (200ns) | 1000 |

| | | | | | | | | |
|---|---|---|---|---|---|---|---|---|
| | - | beta4b | 402965 | 1920 | 127961 | 52 | 5 (250ns) | 1250 |
| | - | beta3 | 282927 | 1925 | 87932 | 52 | 5 (250ns) | 1250 |
| | - | beta5-A+Y | 282965 | 1919 | 87960 | 52 | 5 (200ns) | 1000 |
| | - | beta5_Cterm | 403091 | 1919 | 128006 | 52 | 7 (200ns) | 1400 |
| | - | beta5_midpoint | 283001 | 1919 | 87976 | 52 | 5 (250ns) | 1250 |
| | - | E14 | 305368 | 1916 | 95435 | 56 | 5 (200ns) | 1000 |

**Table S2.** Summary of the MD setups for the Katanin Ring conformation in COMPLEX state

| Species | Nucleotide | Substrate | #Atoms | #Residues | #Water | #Na | #Trajectories | Total Simulation time (ns) |
|---|---|---|---|---|---|---|---|---|
| C.elegans | ATP | beta5 | 306282 | 1919 | 95658 | 67 | 5 (200ns) | 1000 |
| Homo sapiens | ATP | beta5 | 291414 | 1919 | 90582 | 91 | 5 (200ns) | 1000 |
| | ATP | beta3 | 291274 | 1925 | 90516 | 91 | 5 (200ns) | 1000 |
| | ATP | beta5-A+Y | 291327 | 1919 | 90549 | 91 | 6 (200ns) | 1200 |

**Table S3.** Global Average RMSD values for the Katanin Spiral conformation in various states

| | COMPLEX STATE | SUB STATE |
|---|---|---|
| | Global Average (Å) | Global Average (Å) |
| **beta5** | 6.15 ± 0.77 | 6.25 ± 0.68 |
| **beta4b** | 6.32 ± 0.55 | 7.38 ± 0.73 |
| **beta3** | 5.56 ± 0.61 | 7.35 ± 0.70 |
| **beta5-A+Y** | 6.24± 0.77 | 6.42 ± 0.70 |
| **beta5_Cterm** | - | 8.10 ± 1.15 |
| **beta5_midpoint** | 6.81± 0.92 | 6.76 ± 0.90 |
| **E14** | 5.78± 0.60 | 7.02 ± 0.73 |

**Table S4.** Global Average RMSD values for the Katanin Ring conformation in COMPLEX state

| | C.elegans | Homo sapiens |
|---|---|---|
| | Global Average (Å) | Global Average (Å) |

| | | |
|---|---|---|
| **beta5** | 5.91 ± 0.75 | 4.99 ± 0.35 |
| **beta3** | - | 4.91 ± 0.39 |
| **beta5-A+Y** | - | 5.40 ± 0.43 |

**Table S5.** Percentage of central structures from the first clustering step covered by the top two clusters in the second clustering step for each CTT isotype.This was used to determine the optimal cutoff for the second clustering step in the RMSD-based double clustering analysis. A cutoff of 0.55 (highlighted in bold) was selected, as it covers 70% of the structures in all setups.

| RMSD cutoff | **ATP State** | **beta5** | **beta4b** | **beta3** | **beta5-A+Y** | **beta5-mid point** | **E14** |
|---|---|---|---|---|---|---|---|
| 0.15 | 0.92% | 0.80% | 0.98% | 1.10% | 0.98% | 1.15% | 1.10% |
| 0.2 | 6.35% | 5.24% | 6.32% | 6.66% | 5.56% | 7.60% | 7.55% |
| 0.25 | 17.29% | 16.83% | 15.83% | 16.36% | 15.59% | 20.20% | 19.56% |
| 0.3 | 30.07% | 27.68% | 26.39% | 27.17% | 25.86% | 30.56% | 31.50% |
| 0.35 | 42.93% | 36.15% | 35.83% | 36.09% | 36.55% | 38.17% | 39.74% |
| 0.4 | 48.81% | 41.53% | 44.24% | 42.37% | 38.93% | 41.24% | 42.56% |
| 0.45 | 51.11% | 55.43% | 53.68% | 58.47% | 42.49% | 44.69% | 50.52% |
| 0.5 | 51.95% | 77.32% | 81.89% | 80.66% | 55.01% | 61.89% | 76.74% |
| 0.55 | 68.48% | 88.11% | 95.63% | 96.89% | 74.39% | 73.85% | 94.31% |

**Table S6.** The residue numbers of the functional regions of *C.elegans* and *H. Sapiens* katanin.[2,3,4,5]

| **Functional Region** | | **C.elegans Res. No** | **Human Res. No** |
|---|---|---|---|
| **ATP - Binding** | Walker A (WA) | 233 - 240 | 249 - 256 |
| | Walker B (WB) | 290 - 293 | 306 - 309 |
| | Arginine Fingers | 351, 352 | 371, 372 |
| **CTT - Binding** | Pore Loop 1 (PL1) | 263 - 272 | 280 - 289 |
| | Pore Loop 2 (PL2) | 301 - 312 | 317 - 328 |
| **Oligomerization** | Pore Loop 3 (PL3) | 339 - 346 | 359 - 366 |
| | CT-Helix | 456 - 472 | 475 - 491 |

**Table S7.** The proportion of variance covered (%) in the top PCs for the global motions of katanin spiral conformation with natural CTT isotypes. The corresponding PC motions are shown in Figure 3.

| Global | beta5 | beta4b | beta3 | E14 |
|---|---|---|---|---|
| *COMPLEX* | | | | |
| **PC 1** | 33.70% | 25.52% | 35.91% | 34.06% |
| **PC 2** | 21.52% | 18.90% | 16.94% | 17.08% |
| **Variance in top 2 PCs** | 55.22% | 44.42% | 52.84% | 51.14% |
| **Variance in top 10 PCs** | 86.39% | 85.22% | 86.18% | 86.32% |
| *SUB* | | | | |
| **PC 1** | 22.32% | 40.22% | 31.52% | 44.26% |
| **PC 2** | 17.60% | 19.72% | 26.96% | 14.87% |
| **Variance in top 2 PCs** | 39.93% | 59.94% | 58.48% | 59.13% |
| **Variance in top 10 PCs** | 86.35% | 89.23% | 90.25% | 89.14% |

**Table S8.** The proportion of variance covered (%) in the top PCs for the global motions of katanin spiral conformation with constructed CTT isotypes. The corresponding PC motions are shown in Figure 3.

| Global | beta5-A+Y | beta5_Cterm | beta5_midpoint |
|---|---|---|---|
| *COMPLEX* | | | |
| **PC 1** | 36.06% | - | 39.24% |
| **PC 2** | 18.15% | - | 17.57% |
| **Variance in top 2 PCs** | 54.21% | - | 56.81% |
| **Variance in top 10 PCs** | 89.45% | - | 89.26% |
| *SUB* | | | |
| **PC 1** | 40.26% | 33.00% | 37.52% |

| | | | |
|---|---|---|---|
| **PC 2** | 23.23% | 20.17% | 17.36% |
| **Variance in top 2 PCs** | 63.49% | 53.17% | 54.88% |
| **Variance in top 10 PCs** | 89.59% | 91.69% | 88.51% |

**Table S9.** The proportion of variance covered (%) in the top PCs for the PL1 motions of katanin spiral conformation with natural CTT isotypes. The corresponding PC motions are shown in Figure 4.

| **PL1** | **beta5** | **beta4b** | **beta3** | **E14** |
|---|---|---|---|---|
| *COMPLEX* | | | | |
| **PC 1** | 40.27% | 25.88% | 39.73% | 29.53% |
| **PC 2** | 14.25% | 21.80% | 19.30% | 14.97% |
| **Variance in top 2 PCs** | 54.52% | 47.98% | 59.03% | 44.50% |
| **Variance in top 10 PCs** | 84.32% | 82.87% | 86.12% | 80.00% |
| *SUB* | | | | |
| **PC 1** | 35.155 | 38.44% | 35.44% | 32.27% |
| **PC 2** | 18.43% | 22.81% | 26.64% | 21.06% |
| **Variance in top 2 PCs** | 53.58% | 61.25% | 62.09% | 53.33% |
| **Variance in top 10 PCs** | 84.45% | 87.47% | 89.13% | 84.41% |

**Table S10.** The proportion of variance covered (%) in the top PCs for the PL1 motions of katanin spiral conformation with constructed CTT isotypes. The corresponding PC motions are shown in Figure 4.

| **PL1** | **beta5-A+Y** | **beta5_Cterm** | **beta5_midpoint** |
|---|---|---|---|
| *COMPLEX* | | | |
| **PC 1** | 35.52% | - | 29.77 % |
| **PC 2** | 20.93% | - | 23.12 % |
| **Variance in top 2 PCs** | 56.45% | - | 52.89 % |

| | | | |
|---|---|---|---|
| **Variance in top 10 PCs** | 88.46% | - | 85.65% |
| *SUB* | | | |
| **PC 1** | 37.50% | 44.08% | 34.83% |
| **PC 2** | 19.58% | 18.26% | 23.36% |
| **Variance in top 2 PCs** | 57.08% | 62.34% | 58.19% |
| **Variance in top 10 PCs** | 87.54% | 92.35% | 87.66% |

**Table S11.** The RMSIP values between the top 10 PCs of katanin spiral confirmation with natural CTT isotypes compared the E14 bound system.

| RMSIP | COMPLEX | | SUB | |
|---|---|---|---|---|
| | Global | PL1 | Global | PL1 |
| **beta5** | 0.402 | 0.420 | 0.397 | 0.402 |
| **beta4b** | 0.417 | 0.393 | 0.226 | 0.415 |
| **beta3** | 0.413 | 0.406 | 0.395 | 0.414 |

**Table S12.** The RMSIP values between the top 10 PCs of katanin spiral confirmation with constructed CTT isotypes compared the beta5 bound system.

| RMSIP | COMPLEX | | SUB | |
|---|---|---|---|---|
| | Global | PL1 | Global | PL1 |
| **beta5-A+Y** | 0.412 | 0.428 | 0.397 | 0.416 |
| **beta5-midpoint** | 0.411 | 0.420 | 0.422 | 0.419 |
| **beta5-Cterm** | - | - | 0.356 | 0.393 |

**Table S13.** Center-of-mass distances between regions of the terminal protomers in the COMPLEX state of katanin spiral conformation bound to various CTT isotypes. Distances are reported in Å. Structures 1 and 2 correspond to the two extreme conformations along the PC1 space.

| CTT Isotype | | NBD (A) - NBD (F) | Difference | NBD (A) - HBD (F) | Difference |
|---|---|---|---|---|---|
| **beta5** | Structure 1 | 45.74 | -3.43 | 66.69 | -14.01 |
| | Structure 2 | 42.31 | | 52.68 | |
| **beta4b** | Structure 1 | 44.17 | 0.85 | 68.76 | -8.48 |
| | Structure 2 | 45.02 | | 60.28 | |
| **beta3** | Structure 1 | 39.11 | 8.68 | 47.43 | 19.75 |
| | Structure 2 | 47.79 | | 67.18 | |
| **beta5-A+Y** | Structure 1 | 43.05 | 6.30 | 51.05 | 27.57 |
| | Structure 2 | 49.35 | | 78.62 | |
| **beta5-midpoint** | Structure 1 | 42.04 | 6.08 | 20.03 | 52.44 |
| | Structure 2 | 48.12 | | 72.47 | |

**Table S14.** The list of inter-salt bridges involving pore loop 1 and 2 residues of katanin spiral conformation, when bound to natural CTT isotypes, includes those that were present in at least three protomers for more than 10 ns during MD simulations. The average persistence time and the participating protomers are indicated.

| Setup | Salt Bridge | beta5 | beta4b | beta3 | E14 |
|---|---|---|---|---|---|
| COMPLEX | ASP261_ARG275 | AB,BC,CD,DE,EF<br>$31 \pm 24.82$ | AB,BC,CD,DE,EF<br>$24.71 \pm 8.38$ | AB,BC,CD,DE,EF<br>$27.62 \pm 14.93$ | AB,BC,CD,DE,EF<br>$31.75 \pm 11.71$ |
| | ASP269_LYS265 | BA,CB,DC,ED,FE<br>$46.69 \pm 16.08$ | BA,CB,DC,ED,FE<br>$59.26 \pm 20.40$ | BA,CB,DC,ED,FE<br>$63.50 \pm 22.66$ | BA,CB,DC,ED,FE<br>$44.97 \pm 20.33$ |
| | ASP171_LYS265 | - | BA,CB,DC,ED,FE<br>$13.90 \pm 7.50$ | - | - |
| | GLU293_ARG301 | - | - | - | AB,BC,CD,DE,EF<br>$14.35 \pm 8.57$ |

| | | | | | |
|---|---|---|---|---|---|
| SUB | ASP261_ARG275 | AB,BC,CD,DE,EF $41.44 \pm 15.23$ | AB,BC,CD,DE,EF $31.75 \pm 11.21$ | AB,BC,CD,DE,EF $35.10 \pm 11.42$ | AB,BC,CD,DE,EF $53.82 \pm 17.80$ |
| | ASP269_LYS265 | BA,CB,DC,ED,FE $62.71 \pm 12.98$ | BA,CB,DC,ED,FE $63.11 \pm 18.80$ | BA,CB,DC,ED,FE $59.98 \pm 7.92$ | BA,CB,DC,ED,FE $41.37 \pm 9.25$ |
| | GLU308_LYS314 | - | DC,ED,FE $27.40 \pm 16.79$ | - | BA,DC,ED,FE $14.19 \pm 6.62$ |
| | ASP171_LYS265 | - | BA,CB,DC,ED $12.94 \pm 8.88$ | BA,CB,DC $31.74 \pm 6.43$ | BA,CB,DC,ED,FE $22.66 \pm 19.46$ |

**Table S15.** The list of inter-salt bridges involving pore loop 1 and 2 residues of katanin spiral conformation, when bound to constructed CTT isotypes, includes those that were present in at least three protomers for more than 10 ns during MD simulations. The average persistence time and the participating protomers are indicated.

| Setup | Salt Bridge | beta5-A+Y | beta5-midpoint | beta5-Cterm |
|---|---|---|---|---|
| COMPLEX | ASP261_ARG275 | AB,BC,CD,DE,EF $38.49 \pm 16.09$ | AB,BC,CD,DE,EF $20.57 \pm 14.23$ | - |
| | ASP269_LYS265 | BA,CB,DC,ED,FE $51.51 \pm 13.81$ | BA,CB,DC,ED,FE $63.10 \pm 29.76$ | - |
| | ASP171_LYS265 | - | BA,CB,DC,ED,FE $11.29 \pm 12.82$ | - |
| | GLU293_ARG301 | AB,BC,CD,DE,EF $11.22 \pm 7.06$ | AB,BC,CD,DE,EF $14.98 \pm 10.26$ | - |
| SUB | ASP261_ARG275 | AB,BC,CD,DE,EF $37.78 \pm 6.45$ | AB,BC,CD,DE,EF $29.20 \pm 15.53$ | AB,BC,CD,DE,EF $41.92 \pm 7.27$ |
| | ASP269_LYS265 | BA,CB,DC,ED,FE $74.73 \pm 18.66$ | BA,CB,DC,ED,FE $68.45 \pm 17.45$ | BA,CB,DC,ED,FE $59.12 \pm 25.46$ |
| | GLU308_LYS314 | BA,CB,DC,ED,FE $22.03 \pm 13.49$ | BA,CB,DC,ED,FE $27.57 \pm 11.05$ | - |

**Table S16.** The list of intra-salt bridges involving pore loop residues of katanin spiral conformation, when bound to natural CTT isotypes, includes those that were present in at least three protomers for more than 10 ns during MD simulations. The average persistence time and the participating protomers are indicated.

| Setup | Salt Bridge | beta5 | beta4b | beta3 | E14 |
|---|---|---|---|---|---|

| | | | | | |
|---|---|---|---|---|---|
| COMPLEX | GLU271_ARG312 | A,B,D,E,F<br>$10.92 \pm 12.90$ | - | - | A,B,C,D<br>$14.91 \pm 21.39$ |
| | GLU306_LYS314 | A,B,C,D,E,F<br>$63.13 \pm 33.24$ | A,B,C,D,E,F<br>$83.33 \pm 35.51$ | A,B,C,D,E,F<br>$83.73 \pm 10.83$ | A,B,C,D,E,F<br>$61.33 \pm 26.61$ |
| | GLU308_ARG312 | A,B,C,D,E,F<br>$10.42 \pm 6.81$ | A,B,C,D,E,F<br>$11.17 \pm 3.06$ | A,B,C,D,E,F<br>$16.51 \pm 7.97$ | A,B,C,D,E,F<br>$12.13 \pm 8.41$ |
| | GLU316_ARG275 | A,B,C,D,E,F<br>$84.17 \pm 15.89$ | A,B,C,D,E,F<br>$100.00 \pm 38.96$ | A,B,C,D,E,F<br>$81.27 \pm 31.68$ | A,B,C,D,E,F<br>$79.39 \pm 25.70$ |
| | ASP346_ARG301 | A,B,C,D,E,F<br>$19.61 \pm 10.02$ | A,B,C,D,E,F<br>$33.55 \pm 18.16$ | A,B,C,D,E,F<br>$27.10 \pm 14.68$ | A,B,C,D,E,F<br>$29.04 \pm 16.27$ |
| | ASP346_LYS314 | A,B,C,D,E,F<br>$64.78 \pm 28.76$ | A,C,D,E,F<br>$45.84 \pm 23.52$ | A,B,C,D,E,F<br>$33.92 \pm 10.49$ | A,B,C,D,E,F<br>$23.37 \pm 13.73$ |
| SUB | GLU271_ARG312 | A,C,D,E,F<br>$20.07 \pm 15.27$ | A,B,C,E,F<br>$18.61 \pm 14.21$ | A,B,C,D,E,F<br>$23.33 \pm 18.36$ | A,B,C,D,E,F<br>$13.64 \pm 17.55$ |
| | GLU306_LYS314 | A,B,C,D,E,F<br>$72.50 \pm 27.39$ | A,B,C,D,E,F<br>$53.33 \pm 36.79$ | A,B,C,D,E,F<br>$76.05 \pm 26.96$ | A,B,C,D,E,F<br>$45.12 \pm 29.90$ |
| | GLU308_ARG312 | A,B,C,D,E,F<br>$13.38 \pm 10.53$ | A,B,C,D,E,F<br>$16.71 \pm 7.87$ | A,B,C,D,E<br>$10.62 \pm 7.34$ | A,B,C,D,E,F<br>$7.69 \pm 2.63$ |
| | GLU316_ARG275 | A,B,C,D,E,F<br>$98.03 \pm 22.86$ | A,B,C,D,E,F<br>$74.49 \pm 41.68$ | A,B,C,D,E,F<br>$77.60 \pm 21.72$ | A,B,C,D,E,F<br>$101.55 \pm 27.51$ |
| | ASP346_ARG301 | A,B,C,D,E,F<br>$23.60 \pm 18.28$ | - | A,B,C,D,E,F<br>$13.96 \pm 6.05$ | A,B,C,D,E,F<br>$25.35 \pm 19.64$ |
| | ASP346_LYS314 | A,B,C,D,E,F<br>$50.28 \pm 23.48$ | A,B,C,D,E,F<br>$73.29 \pm 22.69$ | B,C,D,E,F<br>$65.20 \pm 10.91$ | A,B,C,D,E,F<br>$63.50 \pm 35.79$ |

**Table S17.** The list of intra-salt bridges involving pore loop residues of katanin spiral conformation, when bound to constructed CTT isotypes, includes those that were present in at least three protomers for more than 10 ns during MD simulations. The average persistence time and the participating protomers are indicated.

| Setup | Salt Bridge | beta5-A+Y | beta5-midpoint | beta5-Cterm |
|---|---|---|---|---|
| COMPLEX | GLU271_ARG312 | A,B,C,D,E,F<br>$10.86 \pm 11.28$ | A,B,C,D,E,F<br>$13.55 \pm 15.27$ | - |
| | GLU306_LYS314 | A,B,C,D,E,F<br>$67.52 \pm 34.89$ | A,B,C,D,E,F<br>$68.88 \pm 15.58$ | - |
| | GLU308_ARG312 | A,B,C,D,E,F<br>$11.26 \pm 5.82$ | A,B,C,D,E,F<br>$8.72 \pm 4.50$ | - |

| | | | | |
|---|---|---|---|---|
| | GLU316_ARG275 | A,B,C,D,E,F<br>$78.89 \pm 30.07$ | A,B,C,D,E,F<br>$90.27 \pm 34.19$ | - |
| | ASP346_ARG301 | A,B,C,D,E,F<br>$26.55 \pm 19.62$ | A,B,C,D,E,F<br>$34.35 \pm 18.61$ | - |
| | ASP346_LYS314 | A,B,C,D,E<br>$49.65 \pm 34.76$ | B,C,D,E,F<br>$62.61 \pm 32.94$ | - |
| SUB | GLU271_ARG312 | A,C,E,F<br>$12.60 \pm 13.12$ | A,B,C,D,E,F<br>$12.03 \pm 13.80$ | A,B,C,D,E,F<br>$16.77 \pm 12.58$ |
| | GLU306_LYS314 | A,B,C,D,E,F<br>$77.82 \pm 29.45$ | A,B,C,D,E,F<br>$81.47 \pm 29.89$ | A,B,C,D,E,F<br>$53.57 \pm 20.52$ |
| | GLU308_ARG312 | A,B,C,D,E,F<br>$8.56 \pm 6.13$ | A,B,C,D,E,F<br>$9.49 \pm 3.91$ | A,B,C,D,E,F<br>$7.06 \pm 2.84$ |
| | GLU316_ARG275 | A,B,C,D,E,F<br>$95.36 \pm 31.31$ | A,B,C,E,F<br>$87.59 \pm 26.59$ | A,B,C,D,E,F<br>$83.13 \pm 21.83$ |
| | ASP346_ARG301 | A,B,C,D,E,F<br>$25.02 \pm 12.18$ | A,B,C,D,E,F<br>$28.71 \pm 10.55$ | A,B,C,D,E,F<br>$15.61 \pm 12.66$ |
| | ASP346_LYS314 | A,B,C,D,E,F<br>$32.68 \pm 9.91$ | B,C,D,E,F<br>$30.21 \pm 18.87$ | A,B,C,D,E<br>$71.74 \pm 34.92$ |

**Table S18.** The proportion of variance covered (%) in the top PCs for the global motions of katanin ring conformation with different CTT isotypes. The corresponding PC motions are shown in Figure S18.

| Global | beta5 C.elegans | beta5 human | beta3 human | beta5-A+Y human |
|---|---|---|---|---|
| **PC 1** | 39.35% | 37.09% | 25.09% | 45.93% |
| **PC 2** | 16.44% | 18.22% | 20.19% | 13.58% |
| **Variance in top 2 PCs** | 55.79% | 53.31% | 45.29% | 59.51% |
| **Variance in top 10 PCs** | 87.47% | 86.89% | 83.58% | 88.57% |

**Table S19.** The proportion of variance covered (%) in the top PCs for the PL1 motions of katanin ring conformation with different CTT isotypes. The corresponding PC motions are shown in Figure S19.

| PL1 | beta5 C.elegans | beta5 human | beta3 human | beta5-A+Y human |
|---|---|---|---|---|

| | | | | |
|---|---|---|---|---|
| **PC 1** | 34.25% | 51.55% | 37.24% | 44.08% |
| **PC 2** | 28.34% | 17.86% | 22.37% | 15.35% |
| **Variance in top 2 PCs** | 62.59% | 69.41% | 59.61% | 59.43% |
| **Variance in top 10 PCs** | 91.44% | 92.74% | 91.29% | 91.11% |

**Table S20.** The RMSIP values between the top 10 PCs of katanin ring confirmation. Human ring conformation with beta5 was compared with C.elegans ring conformation with beta5. Human ring conformation with beta3 and beta5-A+Y were compared with Human ring conformation with beta5.

| **RMSIP** | **beta5-Human** | **beta3-Human** | **beta5-A+Y-Human** |
|---|---|---|---|
| **Global** | 0.331 | 0.343 | 0.374 |
| **PL1** | 0.372 | 0.395 | 0.396 |

**Table S21.** The list of intra and inter salt bridges involving pore loop 1 and 2 residues of C.elegans katanin ring bound to beta5 CTT. Includes those that were present in at least three protomers for more than 10 ns during MD simulations. The average persistence time and the participating protomers are indicated.

| | **Region** | **Salt Bridge** | **beta5 *C.elegans*** |
|---|---|---|---|
| INTER | PL1-PL1 | ASP261_ARG275 | BC,CD,DE,EF $24.55 \pm 7.52$ |
| | PL1-PL1 | ASP269_LYS265 | CB,DC,ED,FE $32.15 \pm 20.31$ |
| | PL2-PL2 | GLU308_LYS314 | CB,ED,FE $12.55 \pm 12.06$ |

| | | | |
|---|---|---|---|
| INTRA | PL1-PL1 | GLU271_ARG275 | A,B,C,D,E,F<br>$11.96 \pm 11.85$ |
| | PL2-PL2 | GLU306_LYS314 | A,B,C,D,E,F<br>$80.84 \pm 29.16$ |
| | PL2-PL2 | GLU308_ARG312 | A,B,C,D,E,F<br>$10.57 \pm 8.92$ |
| | PL2-PL2 | GLU316_ARG275 | A,B,C,D,E,F<br>$70.88 \pm 28.39$ |

**Table S22.** The list of intra and inter salt bridges involving pore loop 1 and 2 residues of human katanin ring bound to different CTT isotypes. Includes those that were present in at least three protomers for more than 10 ns during MD simulations. The average persistence time and the participating protomers are indicated.

| | **Salt Bridge** | **beta5**<br>***Human*** | **beta3**<br>***Human*** | **beta5-A+Y**<br>***Human*** |
|---|---|---|---|---|
| INTER | GLU285_LYS281 | CB,DC,ED,FE<br>$13.07 \pm 4.63$ | CB,DC,ED,FE<br>$32.7 \pm 20.45$ | CB,DC,ED,FE<br>$31.71 \pm 12.90$ |
| | GLU324_LYS330 | - | CD,DE,EF<br>$49.67 \pm 35.93$ | - |
| INTRA | GLU287_ARG328 | A,B,C,D,E,F<br>$13.74 \pm 8.86$ | A,B,C,D,E,F<br>$15.95 \pm 17.26$ | A,B,C,D,E,F<br>$10.06 \pm 7.04$ |
| | GLU322_ARG327 | B,C,D,E,F<br>$17.95 \pm 13.98$ | B,C,D,E,F<br>$22.88 \pm 9.12$ | - |
| | GLU322_LYS330 | A,B,C,D,E,F<br>$62.75 \pm 27.94$ | A,B,C,D,E,F<br>$73.93 \pm 35.71$ | A,B,C,D,E,F<br>$83.17 \pm 46.17$ |
| | GLU332_ARG291 | A,B,C,E<br>$71.17 \pm 26.22$ | A,B,C,D,E,F<br>$50.26 \pm 25.26$ | A,B,C,D,E,F<br>$42.99 \pm 27.44$ |

### Figures

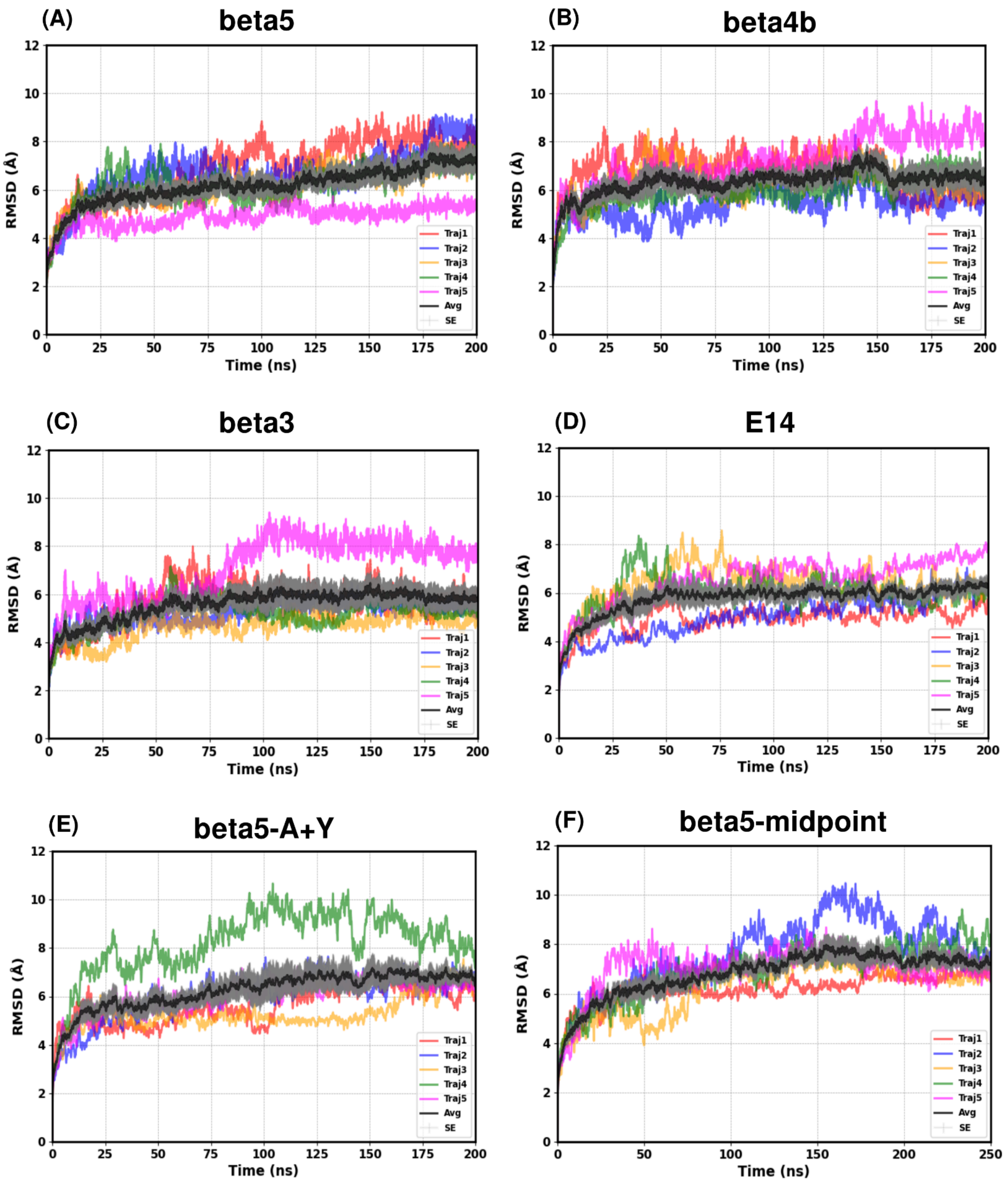

**Figure S1.** RMSD vs time plots, including ensemble averages (black line) and standard errors (gray) over multiple trajectories for katanin spiral conformation in the COMPLEX state with various CTT isotypes. (A) Katanin with beta5 (B) Katanin with beta4b (C) Katanin with beta3 (D) Katanin with E14 (E) Katanin with constructed beta5-A+Y (F) Katanin with constructed beta5-midpoint.

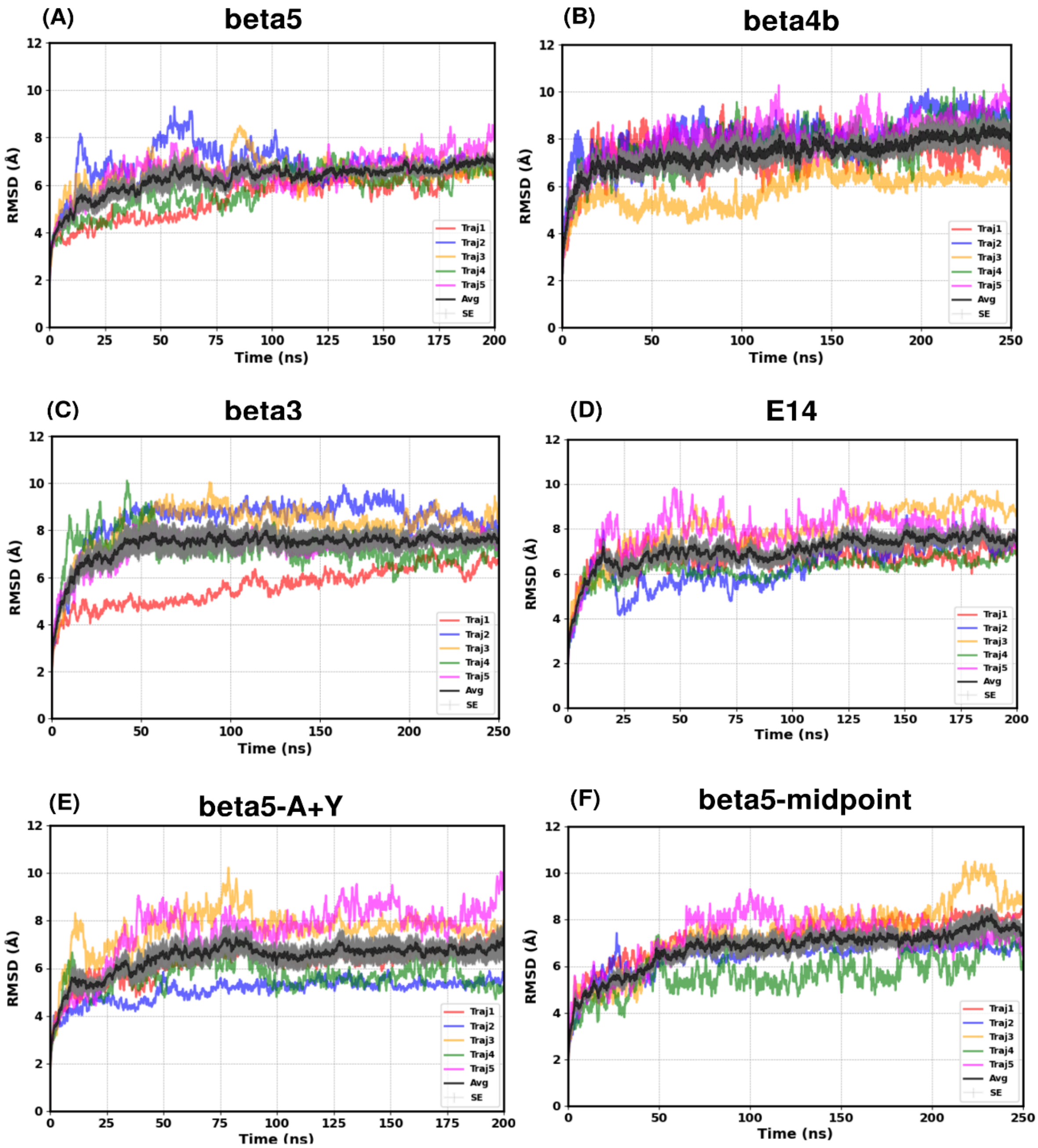

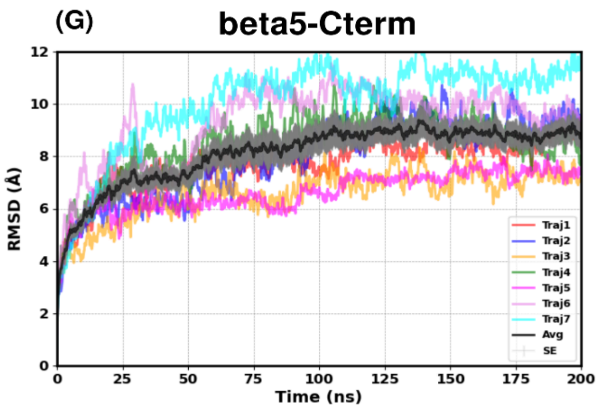

**Figure S2.** RMSD vs time plots, including ensemble averages (black line) and standard errors (gray) over multiple trajectories for katanin spiral conformation in the SUB state with various CTT isotypes. (A) Katanin with beta5 (B) Katanin with beta4b (C) Katanin with beta3 (D) Katanin with E14 (E) Katanin with beta5-A+Y (F) Katanin with beta5-midpoint CTT (G) Katanin with beta5-Cterm.

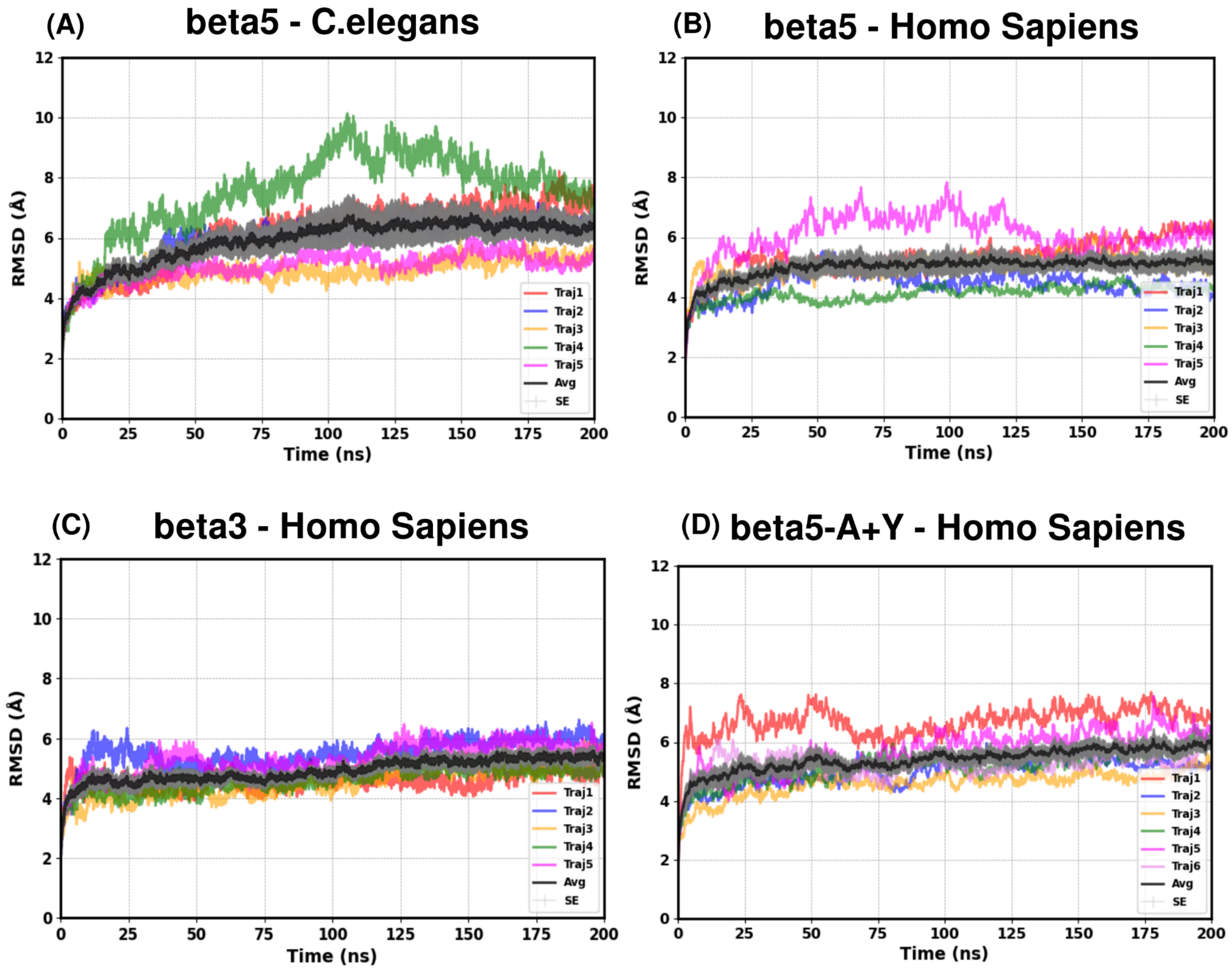

**Figure S3.** RMSD vs time plots, including ensemble averages (black line) and standard errors (gray) over multiple trajectories for katanin ring conformation in the COMPLEX state with various CTT isotypes. (A) *C. elegans* Katanin ring with beta5 (B) *Homo. Sapiens* Katanin ring with beta5 (C) *Homo. Sapiens* Katanin ring with beta3 (D) *Homo. Sapiens* Katanin ring with beta5-A+Y.

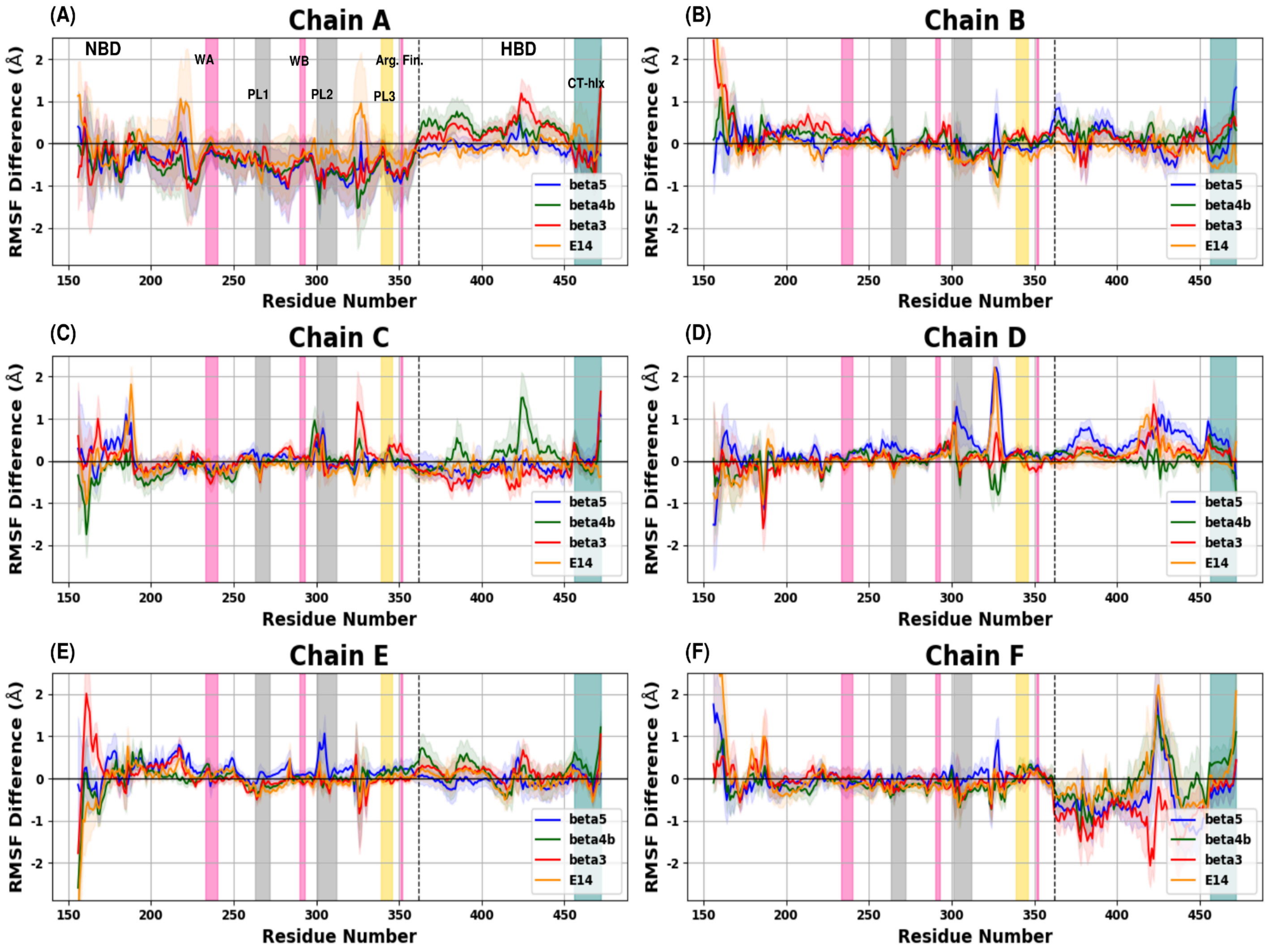

**Figure S4.** Differences between the average RMSF values of each natural CTT isotype in the COMPLEX state and the average RMSF values of the ATP state for the katanin spiral conformation. This plot shows the fluctuations when different natural CTT isotypes are bound to the ATP state (with only ATP). Each plot represents each protomer of the hexamer. (A) Protomer A (B) Protomer B (C) Protomer C (D) Protomer D (E) Protomer E (F) Protomer F. Vertical shaded regions in each plot indicate functionally important regions: WA, WB, Arg Finger (pink), PL1 and PL2 (gray), PL3 (yellow), and the CT-Helix (teal).

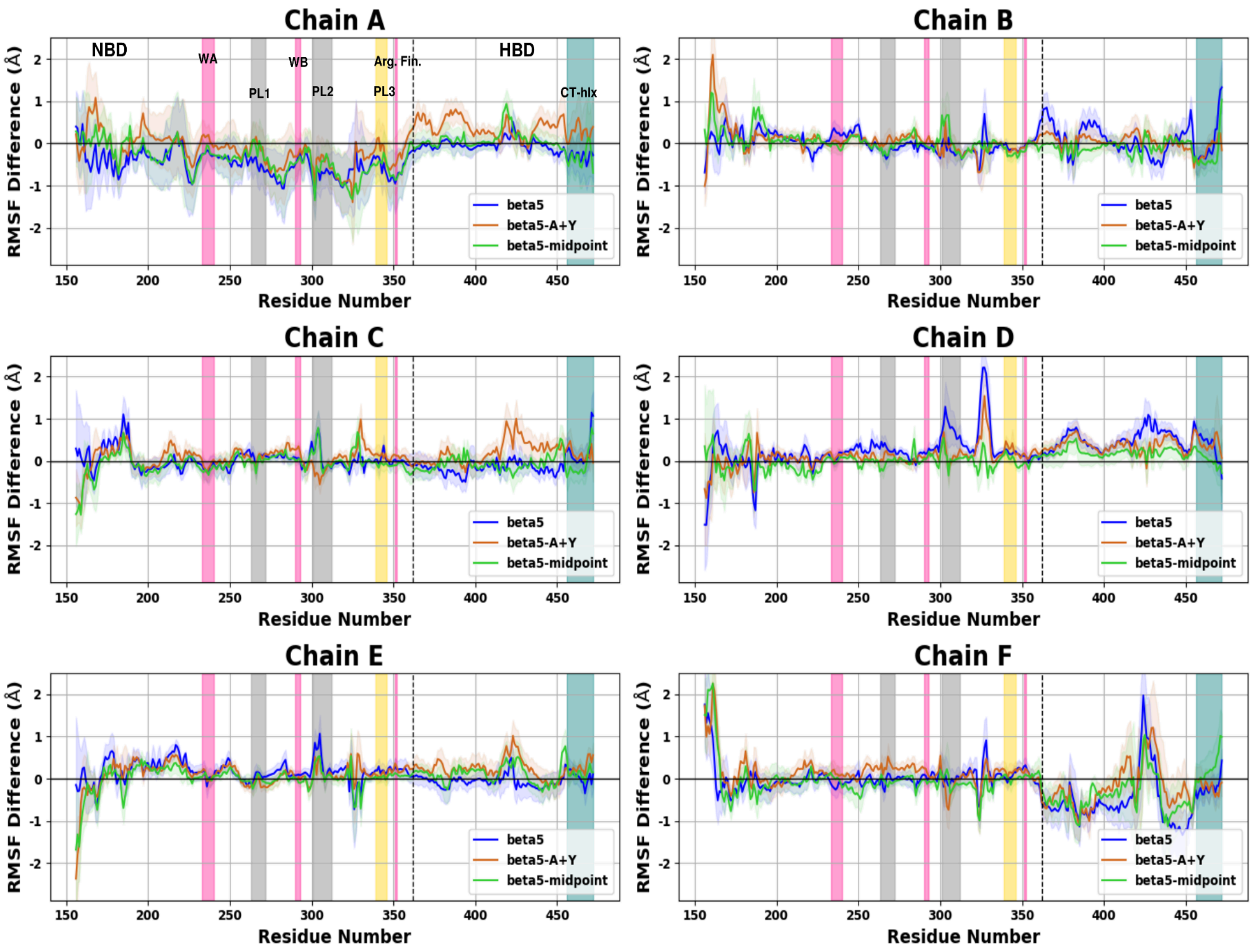

**Figure S5.** Differences between the average RMSF values of each constructed CTT isotype in the COMPLEX state and the average RMSF values of the ATP state for the katanin spiral conformation. This plot shows the fluctuations when different constructed CTT isotypes are bound to the ATP state (with only ATP). Each plot represents each protomer of the hexamer. (A) Protomer A (B) Protomer B (C) Protomer C (D) Protomer D (E) Protomer E (F) Protomer F. Vertical shaded regions in each plot indicate functionally important regions: WA, WB, Arg Finger (pink), PL1 and PL2 (gray), PL3 (yellow), and the CT-Helix (teal).

CLUSTER1

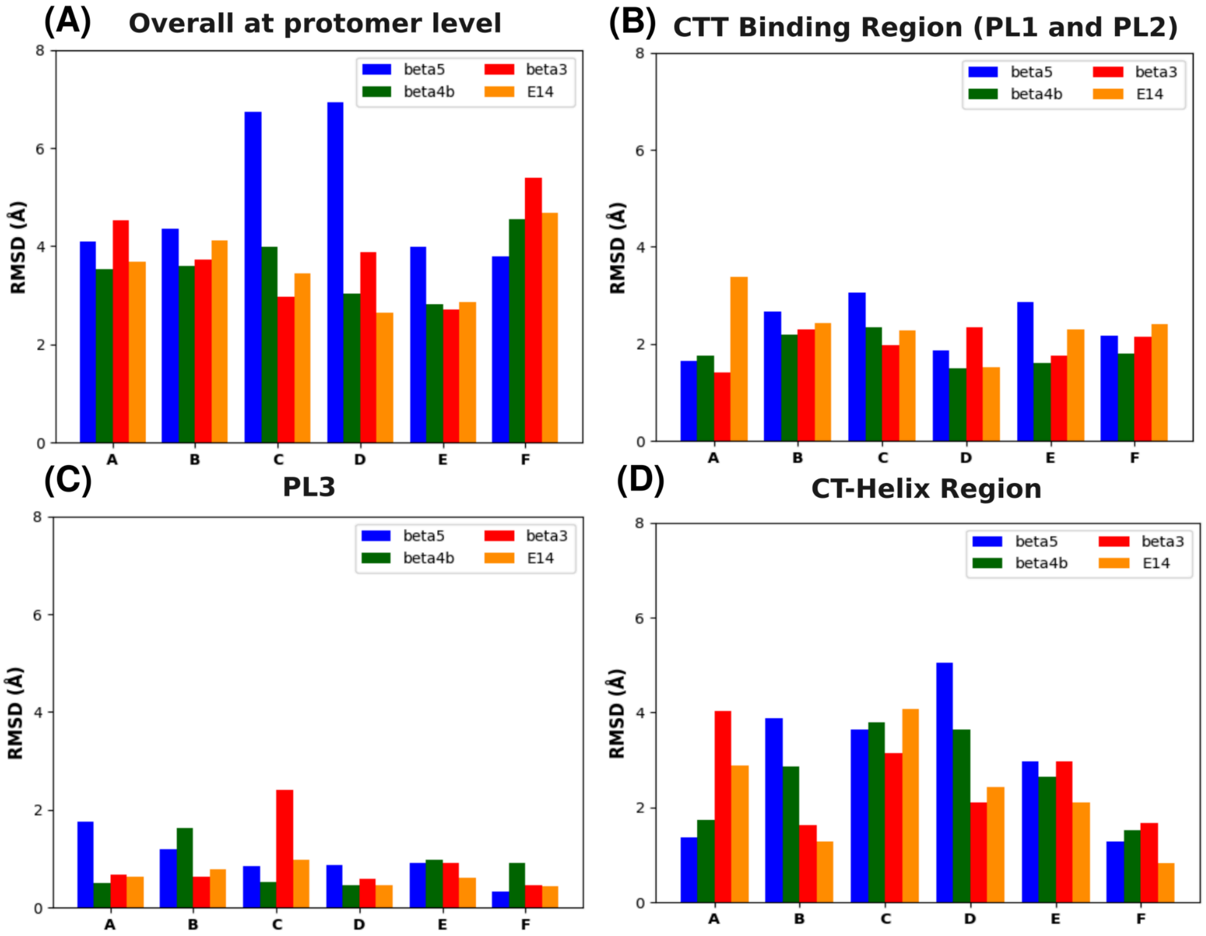

**Figure S6.** The RMSD values of the central structures of the katanin spiral COMPLEX with different natural CTT isotypes compared with the ATP only state for cluster1. (These values were obtained by aligning the functional regions separately). (A) Comparison of each protomer (B) Comparison of the CTT binding region (including both PL1 and PL2) (C) Comparison of the PL3 region (D) Comparison of the CT-helix region. (The residue numbers of the functional regions are listed in Table S6)

### CLUSTER1

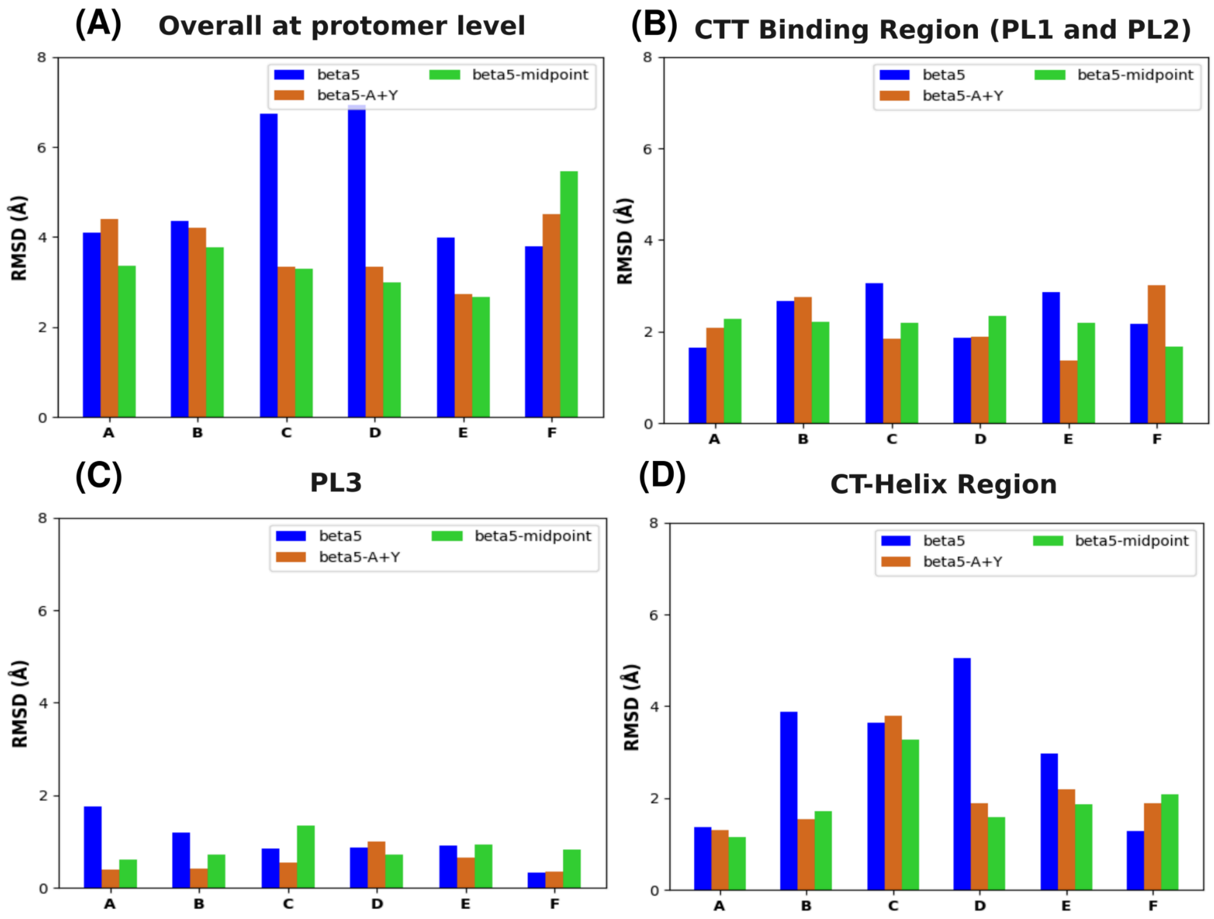

**Figure S7.** The RMSD values of the central structures of the katanin spiral COMPLEX with different constructed CTT isotypes compared with the ATP only state for cluster1. (These values were obtained by aligning the functional regions separately). (A) Comparison of each protomer (B) Comparison of the CTT binding region (including both PL1 and PL2) (C) Comparison of the PL3 region (D) Comparison of the CT-helix region. (The residue numbers of the functional regions are listed in Table S6)

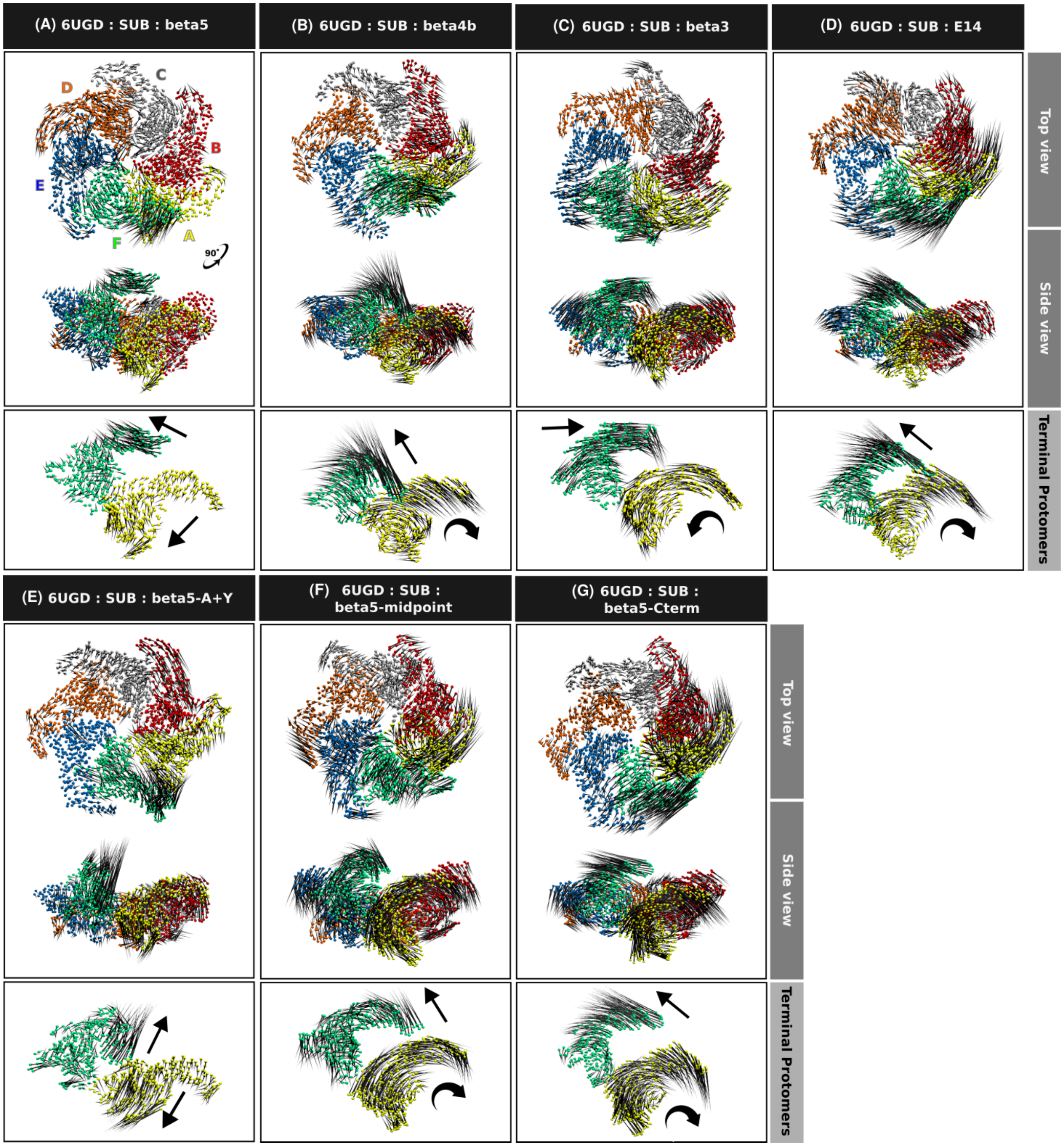

**Figure S8.** The porcupine plots illustrate the global motions corresponding to PC1 of the SUB state of Katanin spiral conformation with different CTT isotypes. (A) katanin with beta5 (B) katanin with beta4b (C) katanin with beta3 (D) katanin with E14 (E) katanin with constructed beta5-A+Y (F) katanin with constructed beta5-midpoint (G) katanin with constructed beta5-Cterm. The variance covered by PC1 for each of these systems is provided in Tables S7-S8.

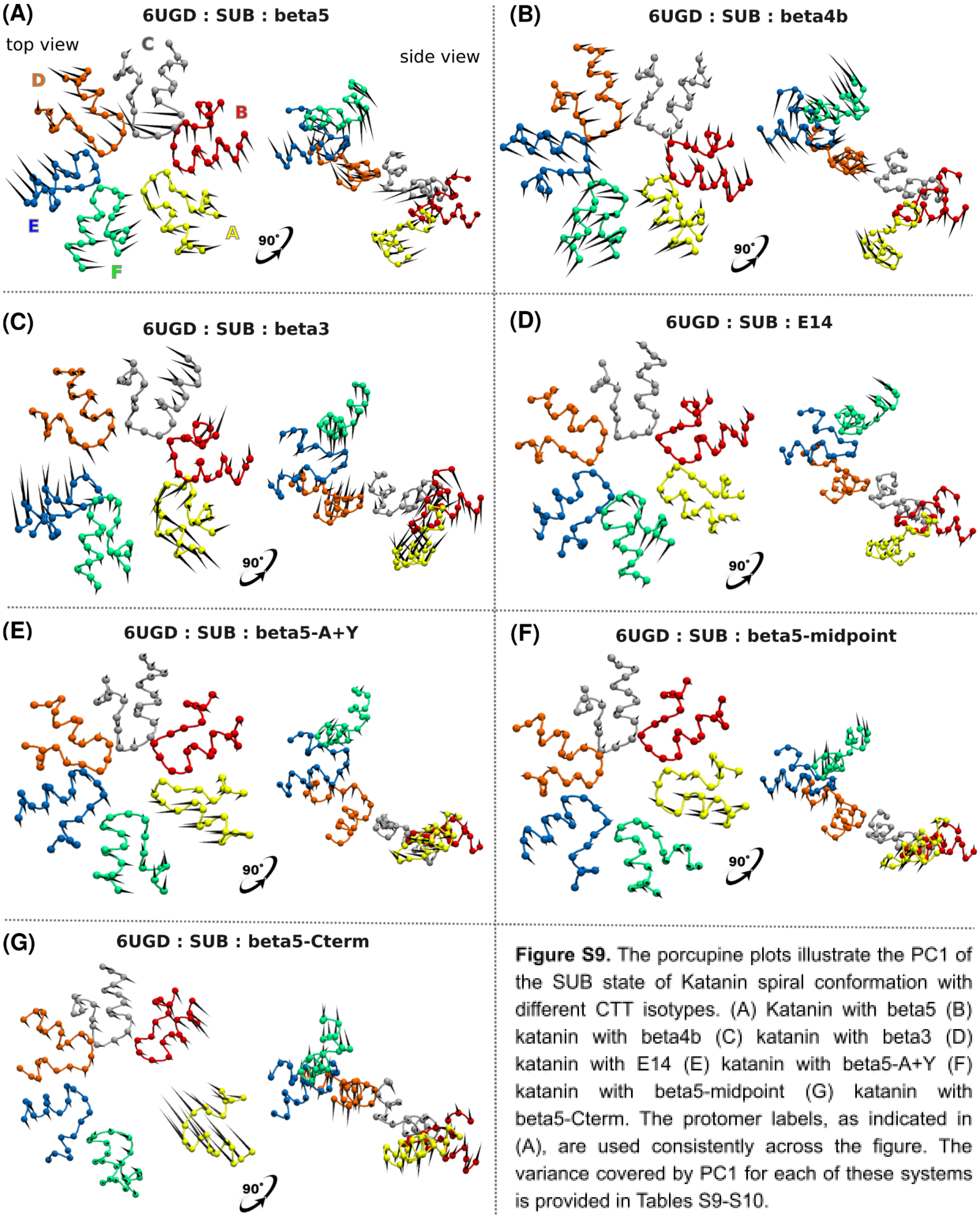

**Figure S9.** The porcupine plots illustrate the PC1 of the SUB state of Katanin spiral conformation with different CTT isotypes. (A) Katanin with beta5 (B) katanin with beta4b (C) katanin with beta3 (D) katanin with E14 (E) katanin with beta5-A+Y (F) katanin with beta5-midpoint (G) katanin with beta5-Cterm. The protomer labels, as indicated in (A), are used consistently across the figure. The variance covered by PC1 for each of these systems is provided in Tables S9-S10.

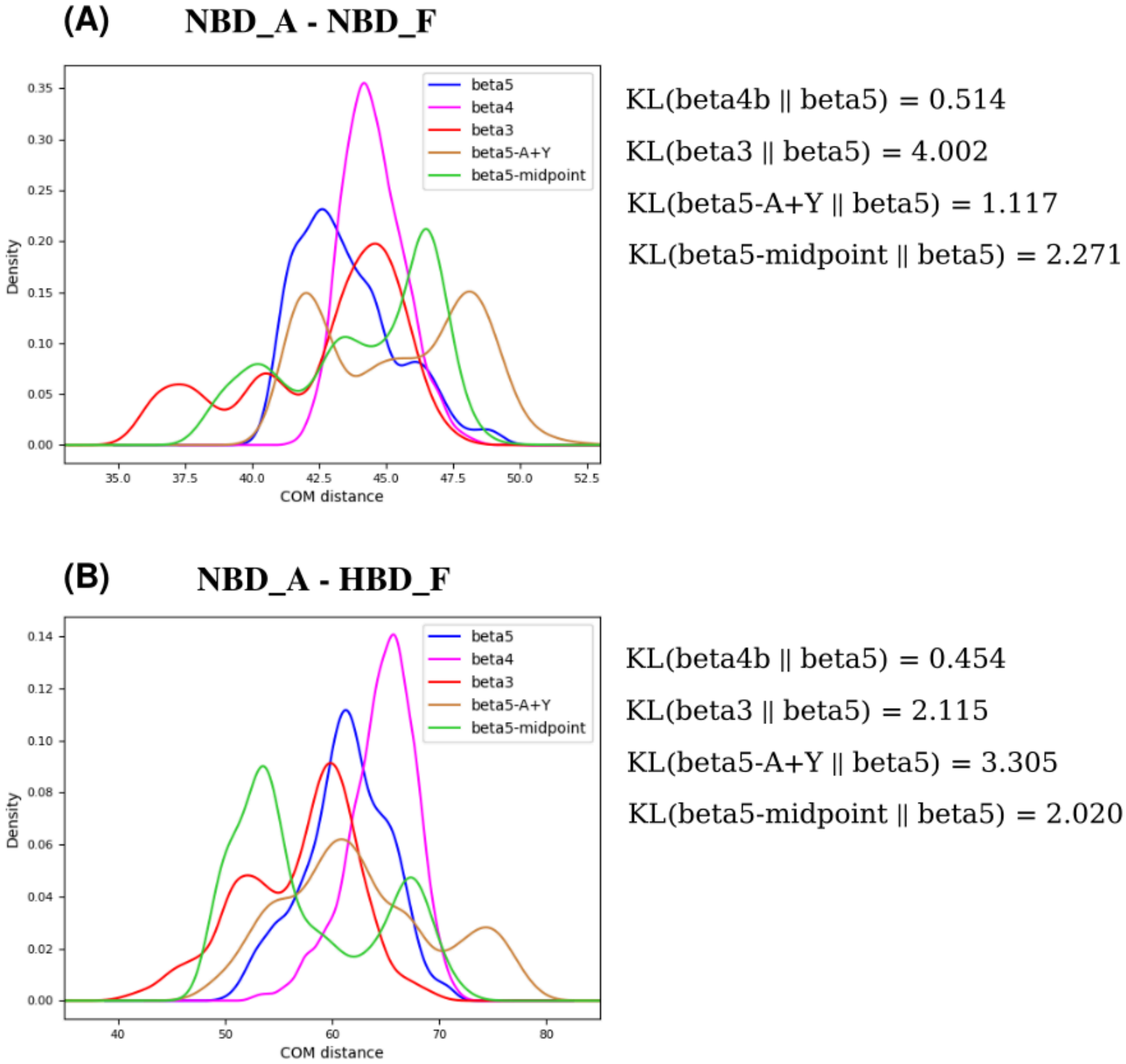

**Figure S10.** The center of mass (COM) distributions of katanin spiral conformation in the COMPLEX state when bound to various CTT isotypes. (A) The COM distance between NBD of protomer A and NBD of protomer F (B) The COM distance between NBD of protomer A and HBD of protomer F. The KL values were calculated to compare the distributions. KL closer to zero indicates similarities between the distributions.

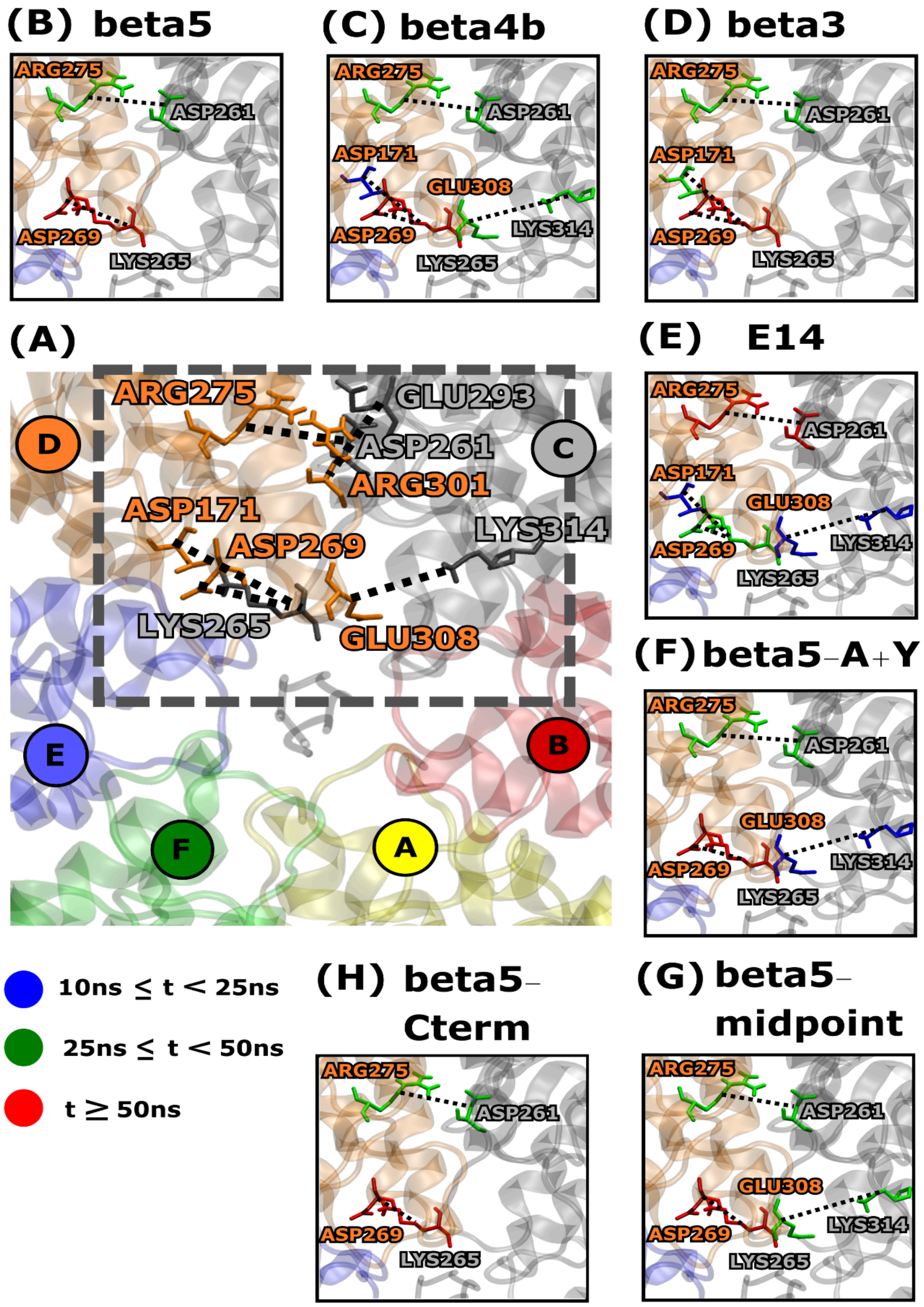

**Figure S11.** Interprotomer salt bridges observed for more than 10 ns in at least three protomer pairs of the katanin spiral conformation in the SUB state. For visualization, the interface between protomers C and D is shown. (A) All interprotomer salt bridges present for more than 10 ns. Residues in grey represent protomer *i–1*, and those in orange represent protomer *i*. Panels (B) – (G) show zoomed-in views of the interprotomer salt bridges specific to each CTT isotype. Colors (blue, green, red) indicate persistence time (*t*) as defined in the legend. If a residue forms multiple salt bridges, its color represents the salt bridge with the highest persistence time.

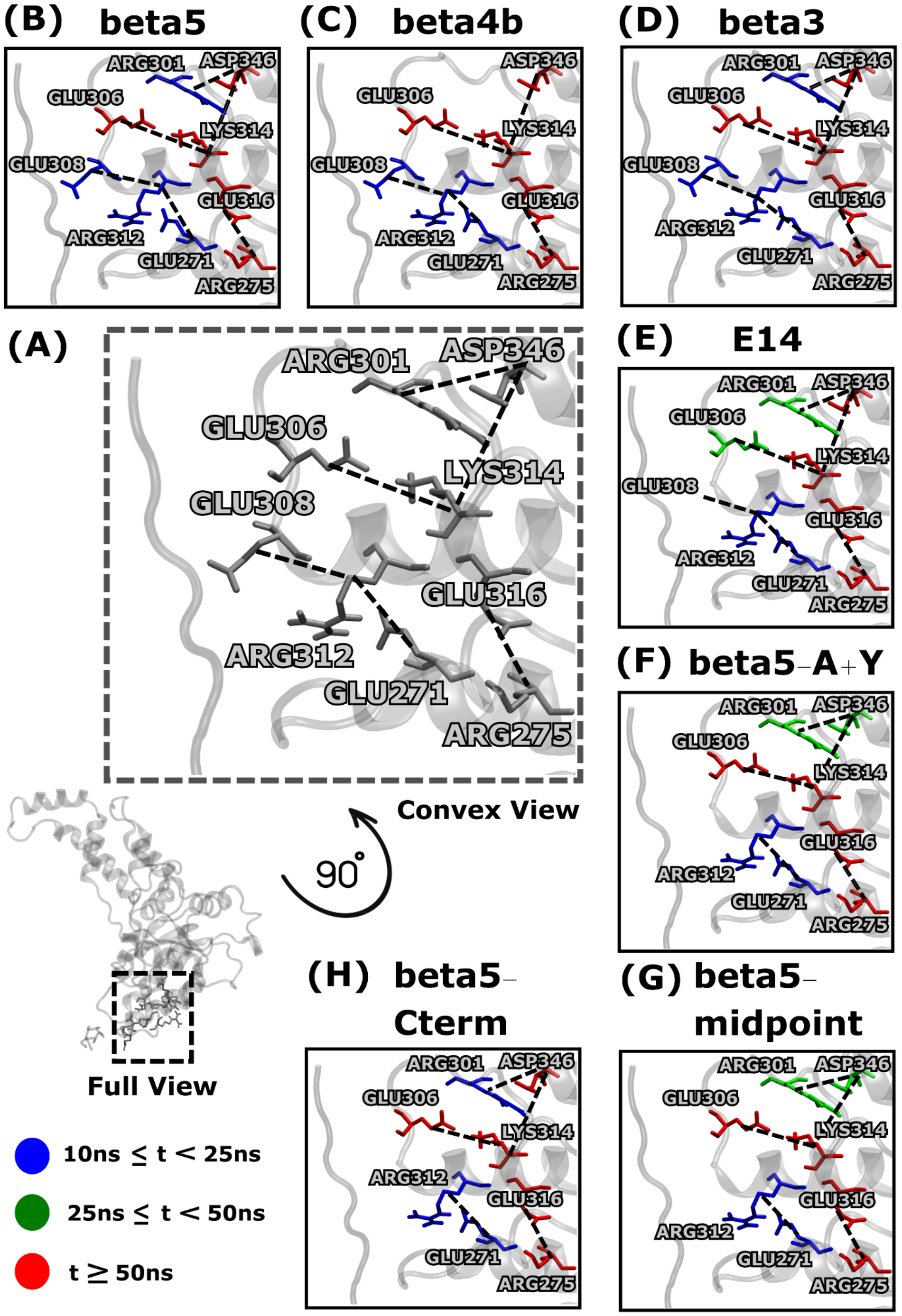

**Figure S12.** Intraprotomer salt bridges observed for more than 10 ns in at least three protomers of the katanin spiral conformation in the SUB state. (A) All intra protomer salt bridges present for more than 10 ns.The convex view is zoomed in to provide better visualization of the salt bridges. Panels (B) – (H) show zoomed-in views of the intra protomer salt bridges specific to each CTT isotype. Colors (blue, green, red) indicate persistence time (*t*) as defined in the legend. If a residue forms multiple salt bridges, its color represents the salt bridge with the highest persistence time.

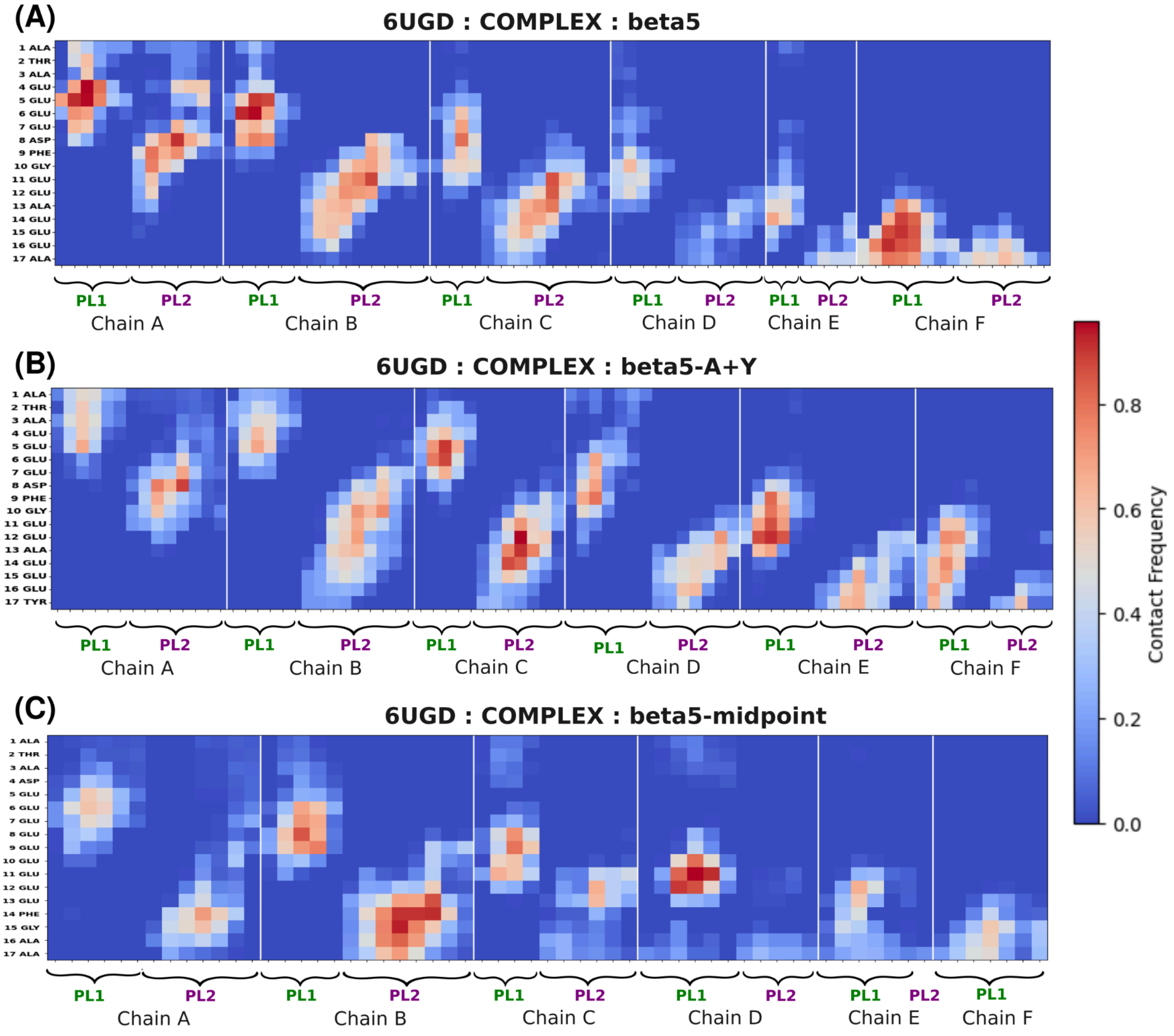

**Figure S13.** Contact maps showing the fraction of contacts between the CTT constructs and the pore loop region (PL1 and Pl2) of the katanin spiral conformation in the COMPLEX state. Here, red indicates that residues are highly in contact during the simulations and blue indicates less contacts. (A) Contact map with beta5 (for reference) (B) Contact map with beta5-A+Y (C) Contact map with beta5-midpoint.

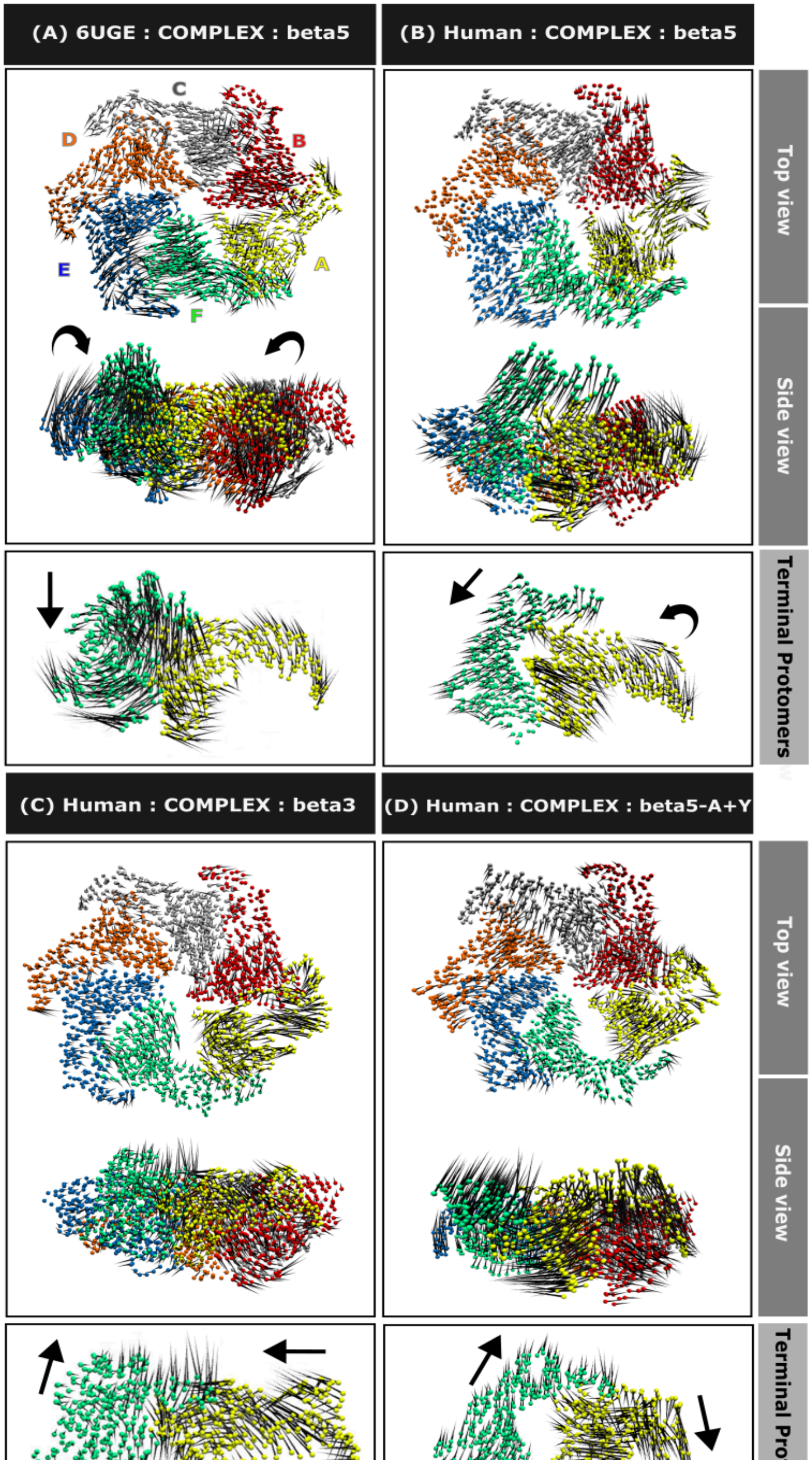

**Figure S14.** The porcupine plots illustrate the global motions corresponding to PC1 of the COMPLEX state of Katanin ring conformation with different CTT isotypes. (A) *C. elegans* katanin ring with beta5

(B) Human katanin ring with beta5 (C) Human katanin ring with beta3 (D) Human katanin ring with beta5-A+Y. The variance covered by PC1 for each of these systems is provided in Table S18.

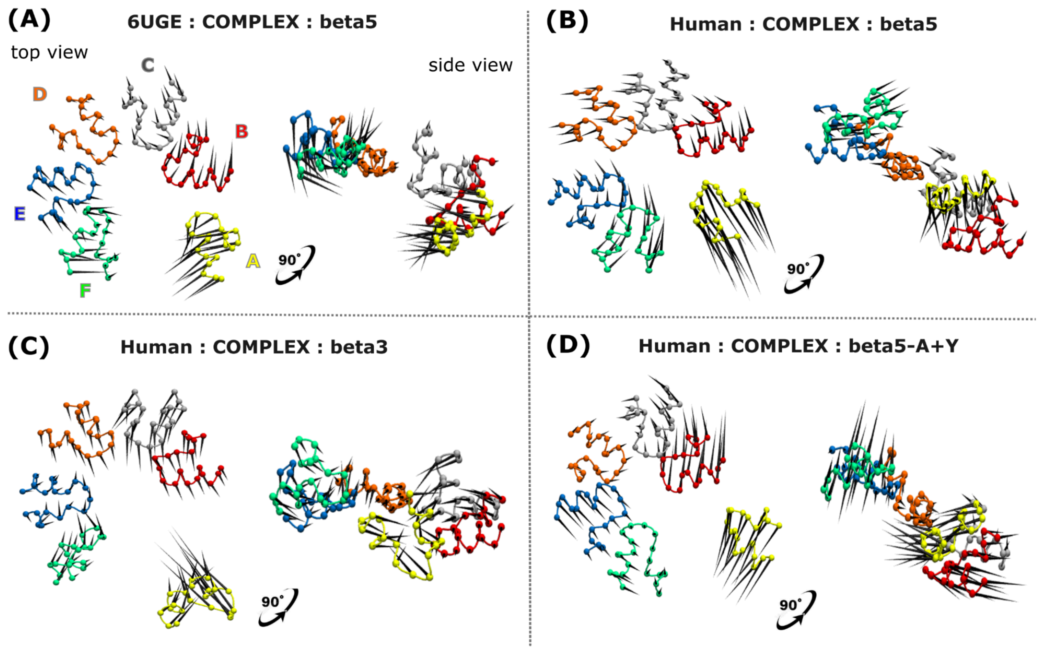

**Figure S15.** The porcupine plots illustrate the PL1 motions corresponding to PC1 of the COMPLEX state of Katanin ring conformation with different CTT isotypes. (A) *C.elegans* katanin with beta5; (B) Human katanin with beta5 CTT; (C) Human katanin with beta3; (D) Human katanin with beta5-A+Y. The protomer labels, as indicated in (A), are used consistently across the entire figure.  A side view of the protein is also included to clearly display the motions of the terminal protomers, A and F. The variance covered by PC1 for each of these systems is provided in Table S19.

**(A)** **NBD_A - NBD_F**

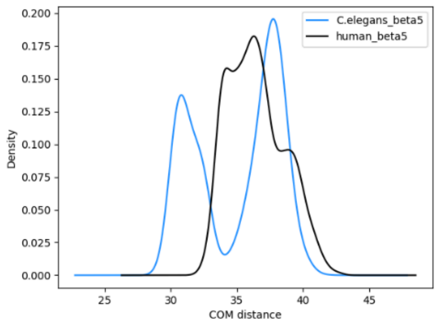

KL(C.ele_beta5 || Human_beta5) = 6.41

**(B)** **NBD_A - HBD_F**

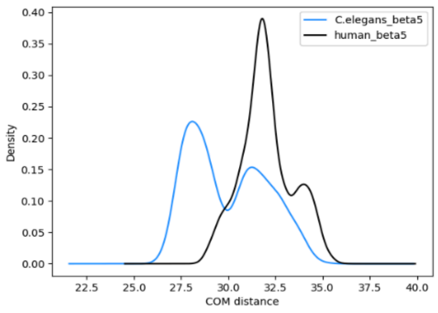

KL(C.ele_beta5 || Human_beta5) = 6.46

**Figure S16.** The center of mass distributions of katanin Ring conformation in different species when bound to beta5. (A) The COM distance between NBD A and NBD F (B) The COM distance between NBD A and HBD F. The KL values were calculated to compare the distributions. KL closer to zero indicates similarities between the distributions.

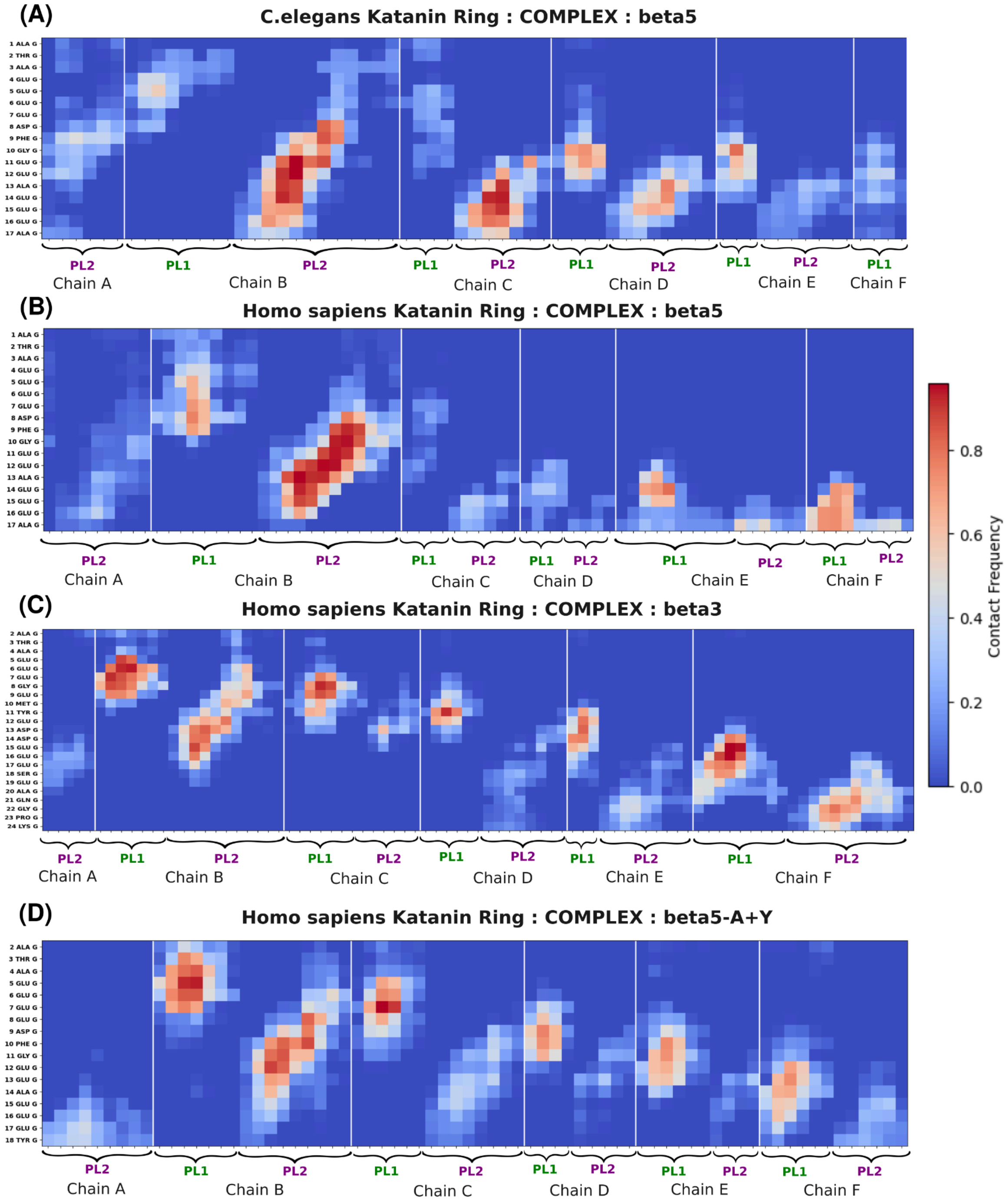

**Figure S17.** Contact maps showing the fraction of contacts between various CTT isotypes and the pore loop region of the katanin ring. Here, red indicates that residues are highly in contact during the simulations and blue indicates less contacts. (A) Contact map between beta5 vs katanin ring (*C.elegans*) (B) Contact map between beta5 vs katanin ring (*Homo sapiens*) (C)

Contact map between beta3 vs katanin ring (*Homo sapiens*) (D) Contact map between beta5-A+Y vs katanin ring (*Homo sapiens*) .

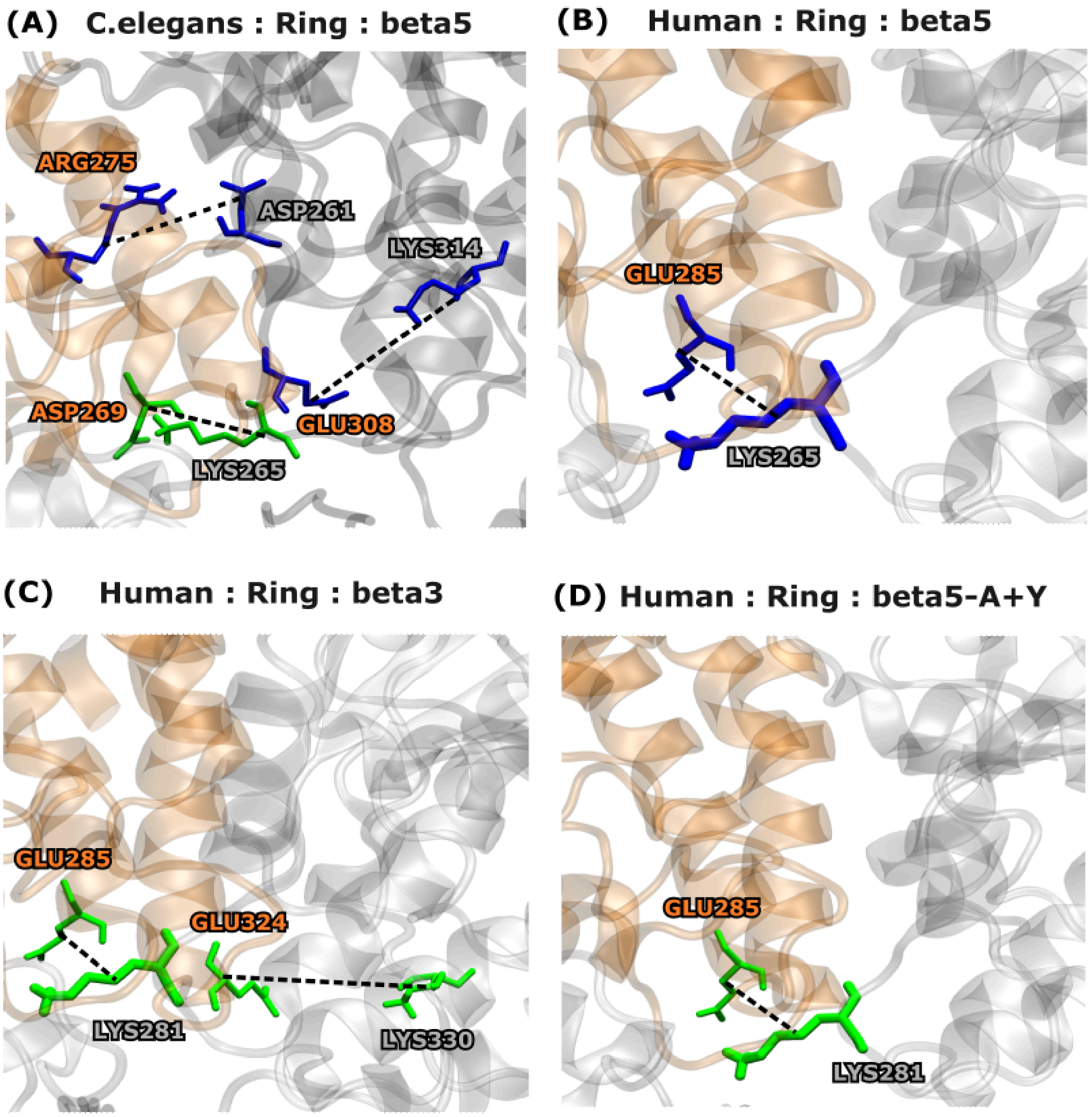

**Figure S18.** Interprotomer salt bridges observed for more than 10 ns in at least three protomer pairs of the katanin ring conformation. (A) *C. elegans* katanin ring with beta5, (B) human katanin ring with beta5, (C) human katanin ring with beta3, and (D) human katanin ring with beta5-A+Y. Salt bridges are colored according to their average persistence time throughout the trajectories, as listed in Tables S21–S24: blue, 10 ns < $t$ < 25 ns; green, 25 ns < $t$ < 50 ns; red, $t$ > 50 ns, where $t$ represents the average persistence time of the salt bridge. If a residue forms multiple salt bridges, its color corresponds to the salt bridge with the highest persistence time.

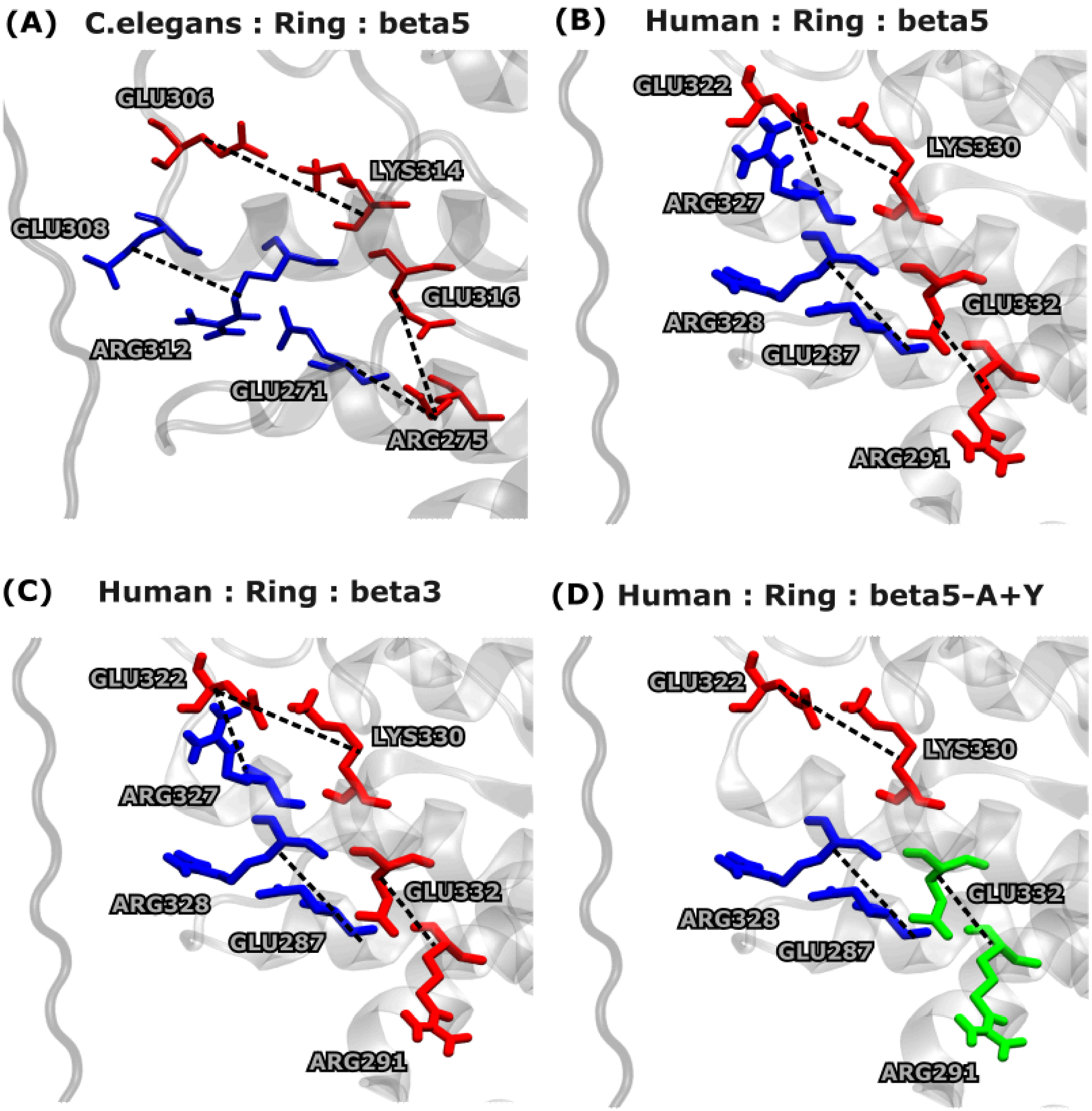

**Figure S19.** Intraprotomer salt bridges observed for more than 10 ns in at least three protomer pairs of the katanin ring conformation. (A) *C. elegans* katanin ring with beta5, (B) human katanin ring with beta5, (C) human katanin ring with beta3, and (D) human katanin ring with beta5-A+Y. Salt bridges are colored according to their average persistence time throughout the trajectories, as listed in Tables S21–S24: blue, 10 ns < $t$ < 25 ns; green, 25 ns < $t$ < 50 ns; red, $t$ > 50 ns, where $t$ represents the average persistence time of the salt bridge. If a residue forms multiple salt bridges, its color corresponds to the salt bridge with the highest persistence time.

(A) **PL1**

| | | | | | | | | | | | |
|---|---|---|---|---|---|---|---|---|---|---|---|
| *C.elegans* | S | S | K | W | R | G | D | S | E | K | I |
| Human | T | S | K | Y | R | G | E | S | E | K | K |

(B) **PL2**

| | | | | | | | | | | | | |
|---|---|---|---|---|---|---|---|---|---|---|---|---|
| *C.elegans* | R | G | N | S | G | E | H | E | A | S | R | R |
| Human | G | G | T | S | E | E | H | E | A | S | R | R |

(C) **PL3**

| | | | | | | | | |
|---|---|---|---|---|---|---|---|---|
| *C.elegans* | T | N | I | P | W | E | L | D |
| Human | T | N | F | P | W | D | I | D |

(D) **CT-Helix**

| | | | | | | | | | | | | | | | | | |
|---|---|---|---|---|---|---|---|---|---|---|---|---|---|---|---|---|---|
| *C.elegans* | P | D | T | M | L | K | C | K | E | W | C | D | S | F | G | A | M |
| Human | A | A | D | I | E | R | Y | E | K | W | I | F | E | F | G | S | C |

**Figure S20.** Sequence alignment of the pore loops and the CT-Helix between the *C.elegans* and the human katanin ring conformation. (A) PL1 (B) PL2 (C) PL3 (D) CT-helix. Conserved residues are highlighted in yellow, while non-conserved substitutions are shown in white.

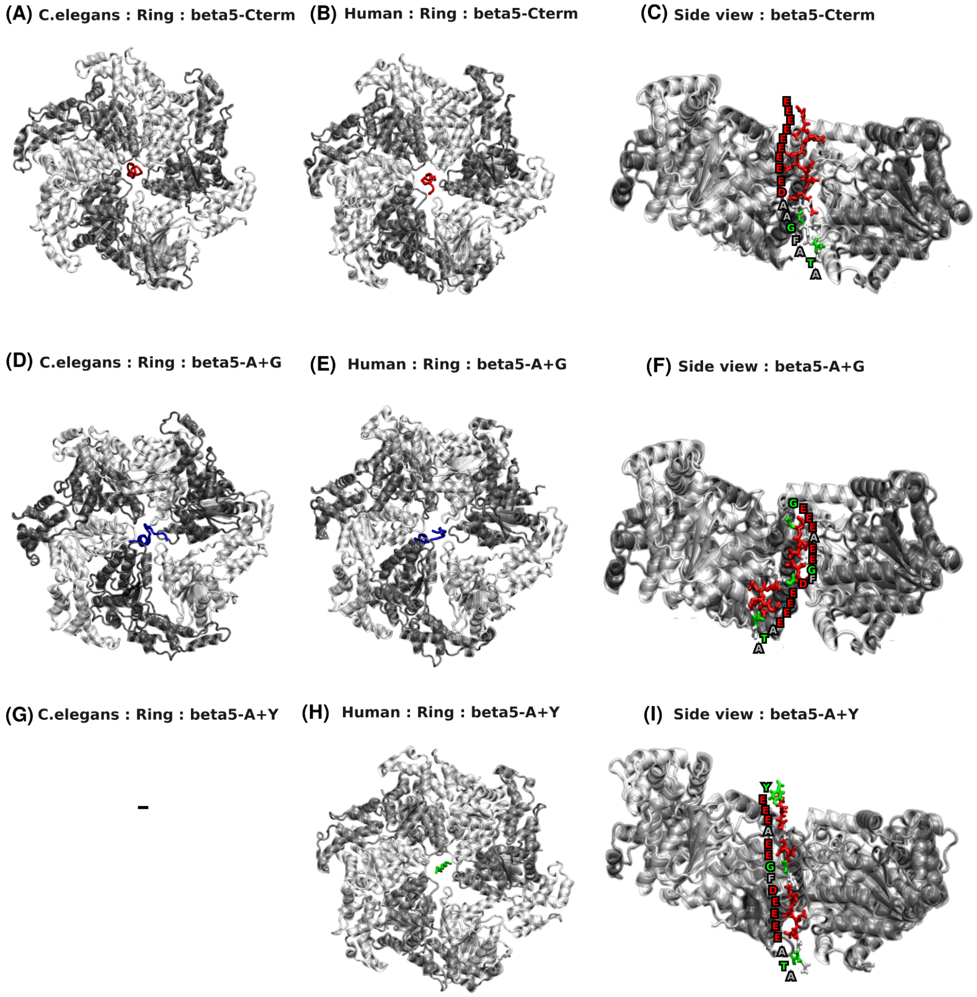

**Figure S21.** Docked katanin ring structures with beta5-Cterm, beta5-A+G and beta5-A+Y CTT isotypes. (A) *C.elegans* katanin ring with beta5-Cterm (B) Human katanin ring with beta5-Cterm (C) Side view of docked katanin structure with beta5-Cterm (D) *C.elegans*

katanin ring with beta5-A+G (E) Human katanin ring with beta5-A+G (F) Side view of docked katanin structure with beta5-A+G (G) beta5-A+Y did not docked into *C.elegans* katanin ring (H) Human katanin ring with beta5-A+Y (I) Side view of docked katanin structure with beta5-A+Y . In figures (C), (F) and (I), a side view is shown to illustrate the sequence of the CTT isotype. Here, red indicates negatively charged residues, green indicates polar uncharged residues, and grey indicates hydrophobic residues.
